# Effects of sub-lethal single, simultaneous, and sequential abiotic stresses on phenotypic traits of Arabidopsis thaliana

**DOI:** 10.1101/2021.12.10.472073

**Authors:** A. Morales, H. J. de Boer, J. C. Douma, S. Elsen, S. Engels, T. Glimmerveen, N. Sajeev, M. Huber, M. Luimes, E. Luitjens, K. Raatjes, C. Hsieh, J. Teapal, T. Wildenbeest, Z. Jiang, A. Pareek, S. L. Singla-Pareek, X. Yin, J.B. Evers, N.P.R. Anten, M. van Zanten, R. Sasidharan

## Abstract

Plant responses to abiotic stresses are complex and dynamic, and involve changes in different traits, either as the direct consequence of the stress, or as an active acclimatory response. Abiotic stresses frequently occur simultaneously or in succession, rather than in isolation. Despite this, most studies have focused on a single stress and single or few plant traits. To address this gap, our study comprehensively and categorically quantified the individual and combined effects of three major abiotic stresses associated with climate change (flooding, progressive drought and high temperature) on 12 phenotypic traits related to morphology, development, growth and fitness, at different developmental stages in four *Arabidopsis thaliana* accessions. Combined sub-lethal stresses were applied either simultaneously (high temperature and drought) or sequentially (flooding followed by drought). In total, we analyzed the phenotypic responses of 1782 individuals across these stresses and different developmental stages.

Overall, abiotic stresses and their combinations resulted in distinct patterns of effects across the traits analyzed, with both quantitative and qualitative differences across accessions. Stress combinations had additive effects on some traits, whereas clear positive and negative interactions were observed for other traits: 9 out of 12 traits for high temperature and drought, 6 out of 12 traits for post-submergence and drought showed significant interactions. In many cases where the stresses interacted, the strength of interactions varied across accessions. Hence, our results indicated a general pattern of response in most phenotypic traits to the different stresses and stress combinations, but it also indicated a natural genetic variation in the strength of these responses.

Overall, our study provides a rich characterization of trait responses of Arabidopsis plants to sub-lethal abiotic stresses at the phenotypic level and can serve as starting point for further in-depth physiological research and plant modelling efforts.

## INTRODUCTION

Climate change has resulted in an overall increase in temperature and increased the likelihood of extreme weather events such as floods, drought episodes, and heat waves (Stott, 2016, Schiermeier, 2011, Meehl et al., 2000). Such events often negatively affect the performance of plants and have a significant impact on food production (Suzuki et al., 2014, Mittler et al., 2012). Extreme weather events also occur simultaneously or sequentially, such as high temperature combined with drought during summer heat waves, or sequential combinations of flooding and drought (Mittler, 2006, Miao et al., 2009). Improvement of plant tolerance and/or ability to recover from these stresses is critical to efforts towards safeguarding global food security in the foreseeable future (Fedoroff et al., 2010).

In recent decades, knowledge on the effect of abiotic stresses on plant survival (tolerance) has advanced significantly but is primarily based on studies focusing on a single stress often using severe treatments such as submergence in darkness (Vashisht et al., 2011). Less is known about the performance of plants under sub-lethal stresses, despite being common in the field, such as mild supra-optimal temperatures, shallow submergence in the light or mild drought (Blum and Jordan, 1985, Chapin, 1991, Zanten et al., 2013). A comprehensive understanding of plant performance under sub-lethal abiotic stresses requires analyzing coordinated changes in functional traits both during the occurrence of stress and upon stress recovery, as opposed to one trait at a time (Thoen et al., 2017, Pandey et al., 2017, Tardieu and Tuberosa, 2010).

In *Arabidopsis thaliana* (Arabidopsis), a mild increase in temperature is known to affect multiple traits such as leaf angle, petiole length, leaf shape, specific leaf area or flowering time (Quint et al., 2016, Zanten et al., 2013, Casal and Balasubramanian, 2019, Jagadish et al., 2016). Drought may affect allocation of assimilates to roots, leaf relative water content and leaf expansion (Tardieu et al., 2018, Chaves et al., 2003). Submergence also affects leaf angles and petiole elongation in some rosette plant species (Voesenek and Bailey-Serres, 2015, van Veen et al., 2016, Sasidharan and Voesenek, 2015). Also, hypoxia and energy impairment associated with submergence can severely reduce growth in many species (Pierik et al., 2005, Sasidharan and Voesenek, 2015, Bailey-Serres and Voesenek, 2008, Vashisht et al., 2011). The recovery phase following water recession presents a different set of stressors for plants. The return to aerobic conditions can cause oxidative stress accompanied by drought-like symptoms, a condition called ‘physiological drought’ (Yeung et al., 2019, Yeung et al., 2018).

Regarding multiple abiotic stresses, studies in Arabidopsis have revealed distinct responses matching specific stress combinations at the metabolic and molecular level. These responses to multi-stress environments were not just a summation of the single stress responses (Mittler, 2006, Rizhsky et al., 2004, Suzuki et al., 2014). Similarly, high temperature and mild drought had interacting effects on some organ- and plant-level physiological and morphological traits in Arabidopsis (Vile et al., 2012). Despite the studies mentioned above, significant gaps remain in our knowledge on the effects of sub-lethal stresses and their combinations on phenotypic traits at the plant level across different developmental stages.

The goals of the current study were to (i) categorically quantify the dynamic effects of several sub-lethal abiotic stresses and their combinations on a wide range of phenotypic traits in Arabidopsis, and (ii) analyze to what extent these effects are conserved/differ across a selection of different natural accessions (Col-0, Bay-0, An-1 and Lp2-6). Three single sub-lethal abiotic stresses and two sub-lethal combinations were used: (i) transient submergence followed by de-submergence, (ii) continuous high temperature, (iii) progressive drought, (iv) progressive drought under continuous high temperature and (v) transient submergence followed by de-submergence and progressive drought.

## MATERIALS AND METHODS

For clarity of presentation, an overview of the different experiments in this study and how they are interrelated is given in the next section. This is followed by a description of the plant growth conditions, the experimental protocols, the measurement protocols with a description of every phenotypic trait studied and a final section on the statistical analysis of the measured traits.

### Overview of experiments

#### Experiment I

This experiment quantified the effects of high temperature, drought and their combination on plant growth and a series of whole-plant developmental and morphological traits of four natural accessions of Arabidopsis. Ten phenotypic traits were measured at three different timepoints (Figure 1).

**Figure 1:**
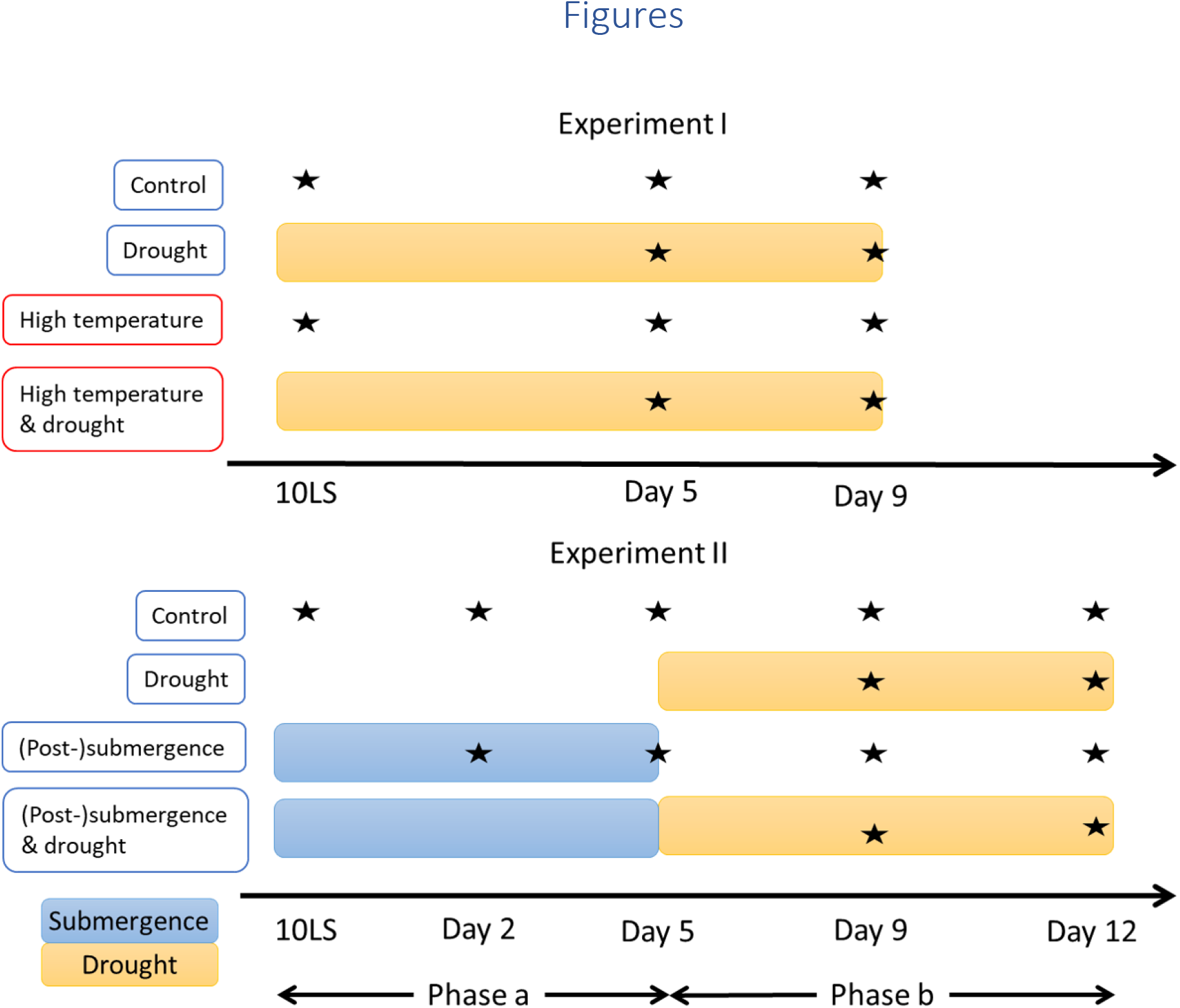
Schemes of experiments I and II showing treatments and harvest timepoints. Each row represents a temporal overview of the treatment or control. Each star represents a harvest timepoint (with days counted from the 10-leaf stage or 10LS). The same temporal schemes were used in experiment III for applying the treatments, but plants were harvested at the time of seed maturity instead, after rewatering to control conditions from respectively day 9 and 12 onwards in experiments I and II.

#### Experiment II

Similar to experiment I, but focused on the effects of submergence, drought, and their sequential combination. The same traits as in experiment I were measured, but there were five timepoints across sequential stresses involving submergence and drought (Figure 1). A separate analysis was performed of the first three timepoints (i.e., when some of the plants were submerged, “phase a” in Figure 1) and the last three timepoints (where none of the plants were submerged, “phase b” in Figure 1).

#### Experiment III

This experiment was performed using the same growth conditions and abiotic stress protocols as experiments I and II but focused on traits related to plant fitness measured at the end of the plant’s life cycle.

### Plant growth conditions

For all experiments, seeds of natural *Arabidopsis thaliana* accessions Col-0 (N1092), Bay-0 (N954), An-1 (N944) and Lp2-6 (N22595) were used (Arabidopsis stock accession numbers between brackets, from www.arabidopsis.info). Plants were sown on a moist soil:perlite mix 1:2 (Primasta BV, Asten, The Netherlands) and stratified in darkness at 4 °C for four days. After stratification, pots were placed in a climate-controlled room at 21 °C (day and night), 70% relative humidity, 120 μmol m^−2^ s^−1^ light intensity (PAR) at plant height provided by fluorescent tubes and 8 h photoperiod. When plants reached the two-leaf stage (ca. two weeks after sowing), the seedlings were transplanted to Jiffy 7c coco pellets (Jiffy Products International BV, Zwijndrecht, The Netherlands).

Prior to transfer of the seedlings, the pellets were soaked in lukewarm water and 50 mL of Hoagland solution until saturation (soaked pellets reached a height of ca. 20 cm). Plants were then cultivated at 21 °C in above-mentioned conditions or in a second climate-controlled room with the same conditions except for a temperature of 27 °C (day and night). Additional Hoagland solution was applied two, six and eight days after transplanting (10 mL, 20 mL, and 10 mL, respectively). Plants were watered every two days except during the application of progressive drought or submergence.

### Experimental protocols

#### Experiment I

As described above, the high-temperature treatment started at the two-leaf stage by moving the plants to the climate-controlled room at 27 °C, while the plants remaining in the original room at 21 °C served as controls. When plants reached the 10-leaf stage, drought was imposed by transferring plants to an empty tray and withholding watering for the duration of the treatment (same procedure in both temperature regimes) under otherwise identical conditions as the well-watered plants. Soil water content was estimated by monitoring the weight of the pellets in which the plants were grown, expressed as pellet weight relative to control conditions. On average, 50% relative pellet weight was reached at day 4 after stopping the watering, 25% at day 7 and 20 % at day 9 (Figure S1).

Plants were harvested at three timepoints corresponding to the 10-leaf stage and 5 and 9 days after the 10-leaf stage, respectively (Figure 1). At the first timepoint, only plants from the control and high temperature groups were harvested as no drought was yet imposed, whereas at the second and third timepoint, plants from the control, high temperature, drought and high temperature and drought groups were harvested.

Typically, six biological replicates were randomly sampled at each combination of accession, treatment and timepoint. However, this was not always possible (e.g., not enough plants might have been available due to mortality), so the realized number of biological replicates was slightly lower (see Table S1 for details). The experiment was executed in eight batches (each batch consisted of one accession subject to all treatments, with two batches per accession).

#### Experiment II

Experiment II was performed entirely in the climate-control room set at 21 °C. When plants reached the 10-leaf stage, randomly selected plants were subjected to complete submergence for five days whereas others remained in control (non-flooded) conditions (Figure 1) under otherwise identical conditions.

To submerge the plants, large containers (54 cm length × 27 cm width × 37 cm depth) were prepared two days prior to the anticipated 10-leaf-stage timepoint. The containers were disinfected with a chlorine tablet and lukewarm water and subsequently drained after at least two hours of disinfection and rinsed thoroughly with water. The containers were then moved to the climate-controlled rooms and filled with water a day before the experiments were started, to allow the water to reach the temperature of the climate room. The submergence was restricted to five days to avoid lethal effects or significant leaf senescence during post-submergence recovery. During submergence, plants were exposed to the same light intensity and quality as in other treatments.

After five days, submerged plants were gently taken out of the water and excess water in the pellet was drained by placing on absorbent paper, until the pellets reached the same weight as those of the well-watered control plants. A subset of the de-submerged plants was then subjected to progressive drought by withholding water (same protocol as in experiment I) while the others received regular watering. A group of plants that had not been submerged were also subjected to progressive drought. Altogether, experiment II thus had two phases:

– Phase a: Started at the 10-leaf stage and lasted for 5 days. This phase only contained control plants and submerged plants. Plants were harvested at three timepoints: at the 10-leaf stage (start of submergence), and two and five days after 10-leaf stage (Figure 1).
– Phase b: Started five days after the 10-leaf stage. There were four groups of plants: control, drought, post-submergence and post-submergence drought. The first harvest timepoint of this phase corresponds to the last timepoint of phase a, and it was followed by two additional harvest timepoints, 9 and 12 days after the 10-leaf stage (Figure 1), which corresponded to 4 and 7 days after the end of submergence, respectively.

Experiment II also aimed at six biological replicates in each combination of accession, treatment and timepoint but the realized number of biological replicates was slightly lower (see Table S1 for details). This experiment was also executed in eight batches (each batch consisted of one accession subject to all treatments, with two batches per accession).

#### Experiment III

In experiment III, plants were grown as in experiment I and II and subjected to the same stress treatments and duration. However, instead of harvesting them for phenotypic analysis, the plants were labelled and allowed to develop further while being monitored until seed harvest. Plants that were subjected to drought treatment (including those in which drought was combined with other stresses) were returned to well-watered conditions after the drought period. In the case of high temperature and high temperature and drought, plants were kept in the climate-controlled chamber at 27 °C up to the seed harvest.

In all cases, as soon as plants bolted, an Aracon system (Betatech BVBA) was fitted around the plant and kept until harvest. The base of the Aracon system collected the seeds from open siliques, while the Aracon tube prevented contamination from – and dispersal to – neighboring plants. When plants completely senesced, the inflorescence was cut at the base and all seeds attached to the plant were removed and sieved (Retsch Gmbh, mesh size: 425 μM), combined with the material in the Aracon base and collected in a paper bag.

### Trait measurement protocols

In the following, the measured traits are indicated in *italics*. For each harvested plant in experiment I and II, a lateral picture was taken using a conventional photo camera of the first or second youngest leaf with a petiole length larger than 1 cm at rosette base height, to determine the petiole insertion angle with respect to the horizontal plane (*leaf angle*). The rosette was then directly harvested using a forceps and razor blade, weighed to determined fresh weight, and dissected leaf-by-leaf with a razor blade. The harvested leaves were laid out on a plastic overhead projector sheet according to their developmental age and digitized with a flatbed scanner HP ScanJet G3110 (Hewlett-Packard, Palo Alto, CA, USA) at 600 dpi. The leaves were then oven-dried at 70 °C for at least 72 h and weighed on a precision scale to determine the *rosette dry weight* and *relative water content*. In addition, directly following the harvesting of above-ground parts, the primary roots were retrieved from the growth substrate by rinsing the substrate off with tap water and the maximum depth reached by the root system recorded (*root length*).

All the imaged true leaves were analyzed with ImageJ v. 1.52a (National Institutes of Health, USA) to determine total plant area, number of rosette leaves (*early rosette leaf number*), average ratio between blade width and length as an index of blade shape (*blade shape*) and average ratio between petiole length and total leaf (i.e., petiole + blade) length (*petiole ratio*). From the plant area and *early rosette leaf number*, the *leaf size* was derived and from the *rosette dry weight* and the plant area, the average *specific leaf area* including petioles was calculated.

In experiment III, the total amount of seeds retrieved from each individual plant was weighed on a precision scale to calculate the seed yield of each individual plant (*yield*) following a sedimentation-based cleaning method described in Morales et al. (2020). In addition, the time to opening of the first flower (*flowering time*) and the total number of rosette leaves produced at the moment of flowering (*total rosette leaf number*) were recorded per plant.

### Statistical analysis

A linear mixed model (random intercept and slopes) was fitted to each trait in experiments I and II (separating phase a and b, Figure 1) using the lme function in the R package nlme (Pinheiro et al., 2021). Either a logarithmic (for positive traits) or logit (for traits between 0 and 1) transformation was used on each trait (Table S1). Each model assumed a linear response of the transformed trait to time where the slope and intercepts were estimated as a function of treatment and accession (fixed effects), and a random effect was used to account for variation across batches.

In each model, the time variable was set to zero at the first measurement timepoint. This means that in experiment I and phase a of experiment II the intercepts of the models corresponded to trait values at the 10-leaf stage, whereas for phase b of experiment II it corresponded to trait values five days after the 10-leaf stage (Figure 1).

The six possible treatment groups (Figure 1) were encoded as the product of three binary variables representing whether the plants were (i) grown under high temperature or not, (ii) subjected to drought or not, (iii) subjected to submergence or not. The experimental design was not full factorial as high temperature and submergence were not combined. The effect of accession was encoded as a categorical variable representing the four possible accessions (An-1, Bay-0, Col-0 or Lp2-6). All main effects and interactions were included in the models.

Once a model was fitted, F tests (significance level of 5%) were performed on each main effect and interaction, using marginal sum of squares. In addition, the following post-hoc tests at 5% significance level were performed with the *emmeans* R package (Lenth, 2021), using Tukey’s method:

i. differences in mean values across treatments at the 10 leaf-stage of experiment I, tested separately for each accession,
ii. differences in slopes across all treatments, tested separately for each accession (in experiments I and II, both phases). These slopes represent the rates of change over time of transformed valued of the traits and hence capture the dynamic effects of transient treatments (drought, submergence and combinations thereof).

The analysis described above emphasizes the trait responses to drought and submergence as a function of time (slopes in the mixed effect models) rather than their effects on mean trait values at the end of the experimental period. We do this because values of traits at the end of the treatment are determined by the length of the treatment and the starting values of the traits (especially for those traits associated to growth or development of the plants). Such analysis would thus not generalize well, as they would be highly sensitive to the duration of the drought or submergence treatment period and the developmental age of the plants at the beginning of the treatment. However, the dynamic effects (i.e., rates of change, represented in this study by slopes) was considered more useful as it would be more sensitive to the intensity of the stress imposed, the physiological response of the plant and potential differences among accessions.

The analysis of experiment III also used linear mixed effect models on transformed trait values (Table S1) to account for the effects and interactions of treatment and accession (fixed effects) and experiment batch (random effect) but, unlike for experiments I and II, there was no time component in this experiment and hence slopes were not computed. Differences in mean values across all accessions and treatment combinations were tested using the same methodology described above for experiment I.

## RESULTS

Overall, abiotic stresses and their combinations resulted in distinct patterns of effects across the traits analyzed, with both quantitative and qualitative differences across accessions. Stress combinations had additive effects on some traits, whereas clear positive and negative interactions were observed for other traits: 9 out of 12 traits for high temperature and drought, 6 out of 12 traits for post-submergence and drought showed significant interactions. The detailed results of the different experiments are described below trait-by-trait, in alphabetical order.

For conciseness, while all the results are shown in Figures 2 – 6, only those effects that were statistically and physiologically significant are reported in the text unless it is pertinent to emphasize a lack of effect. Whenever the statistical significance of the results was not consistent (e.g., a significant effect of drought in experiment I but not in experiment II), the most stringent result was chosen to avoid reporting excessive false positives. The individual measurements for all the experiments and traits together with the fitted models are reported graphically in supplemental Figures S2 – S28. The P values of the F tests for each fitted model are provided in Tables S2 – S5.

**Figure 2:**
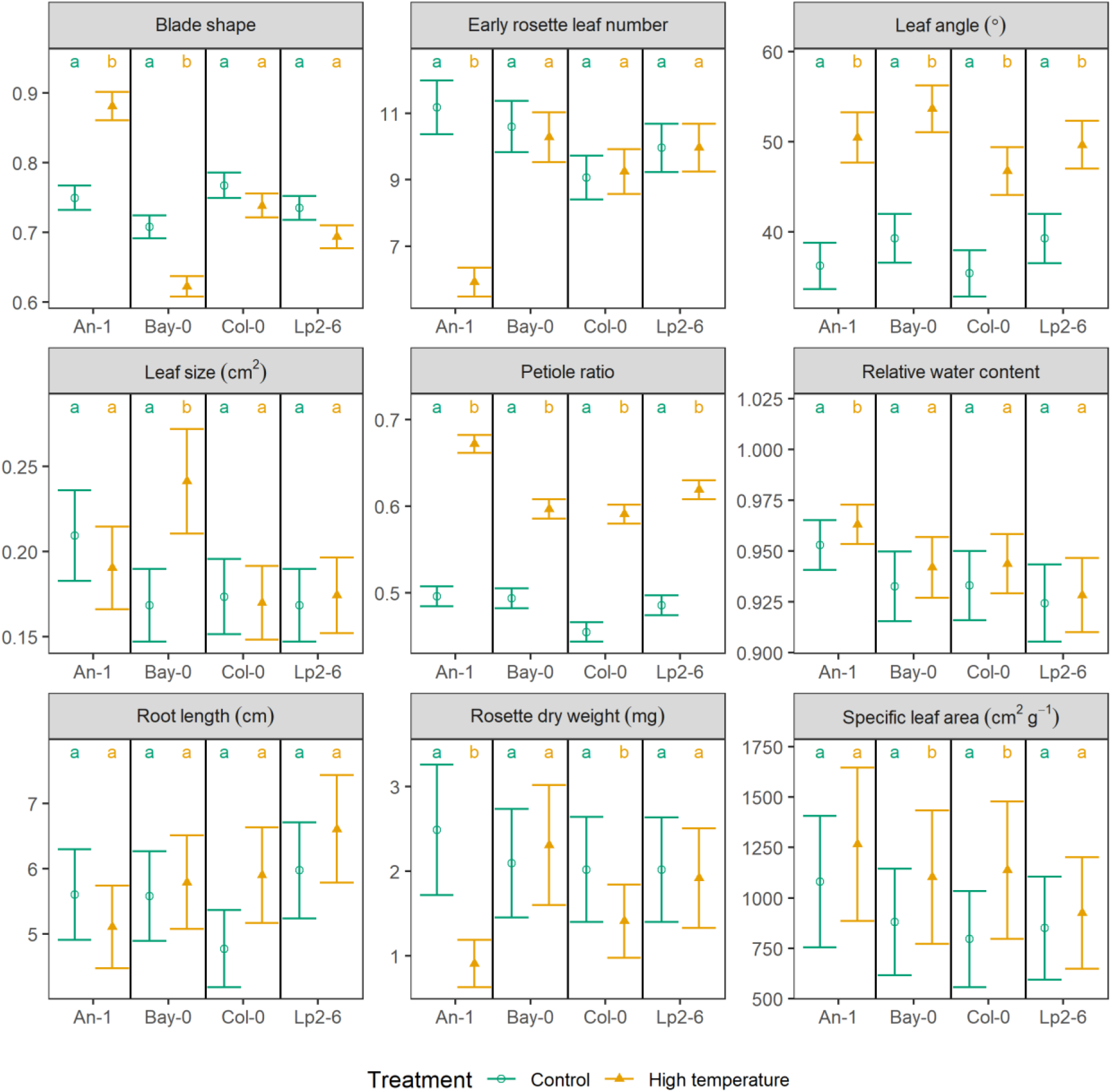
Estimated average values of each trait in control and high temperature conditions at the 10-true leaf stage of experiment I (see text and Figure 1 for details) for each accession and treatment as predicted by the linear mixed models fitted to the data on each trait. Within each panel and accession, if two groups share the same letters it implies no significant difference at the 95% confidence level (the significant tests were performed in the transformed scale, the means are reported in the original scale, Tukey correction for multiple comparison was applied). Units of measure are displayed in the heading of each panel if applicable. Whiskers indicate standard errors of the means.

### Blade shape

The *blade shape* corresponds to the average ratio between leaf blade width and length and decreased over time under control conditions (Figures 3 – 5). High temperature (with or without drought) accelerated this decrease in An-1 (Figures 3 and S2), although the value at the 10-leaf stage was higher when compared to control conditions (Figures 2 and S2). On the other hand, Bay-0 had systematically lower *blade shape* values under high temperature (Figure 2) but with the same dynamics as in control conditions.

**Figure 3:**
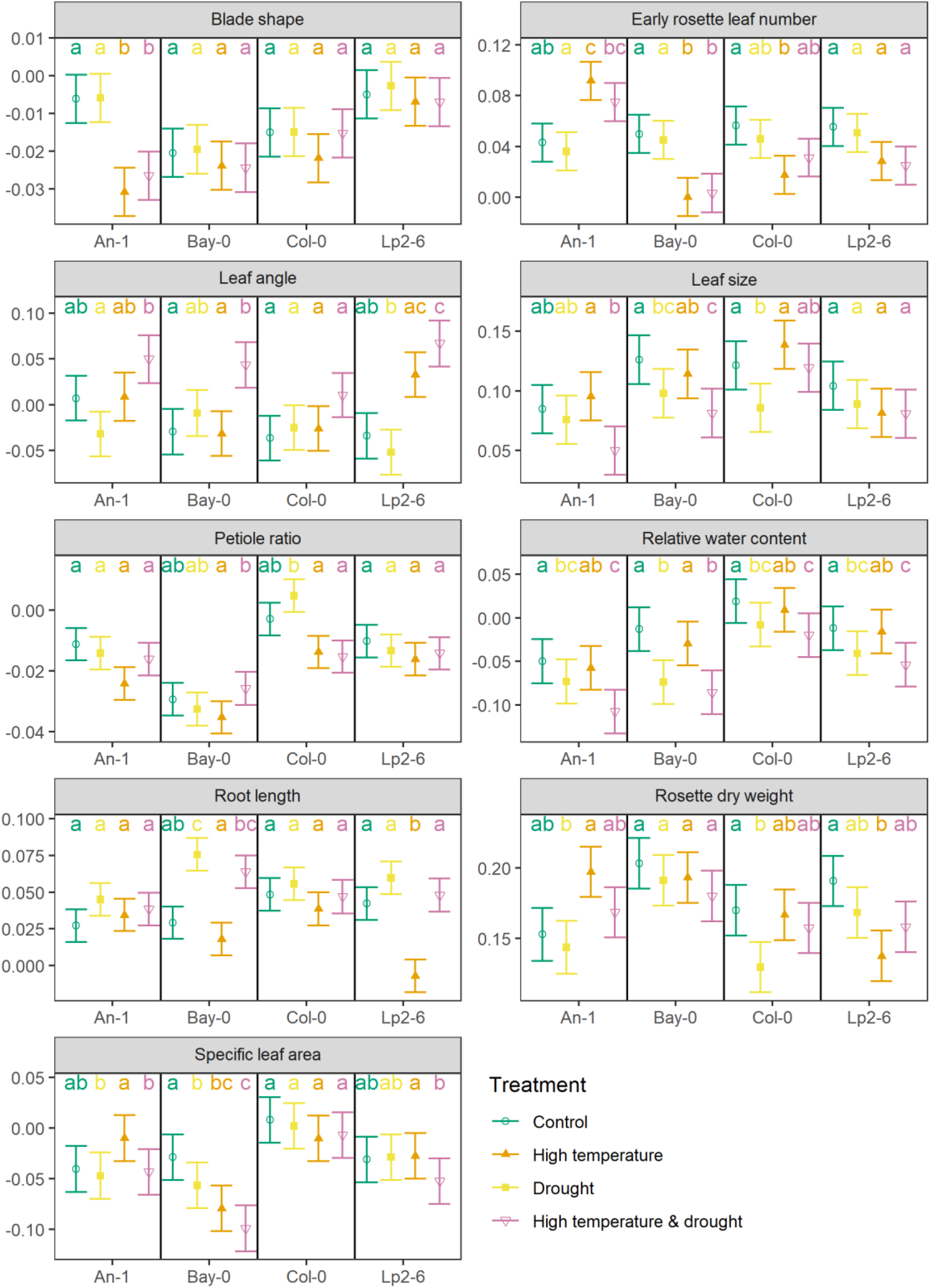
Estimated rate of change over time of each trait (after being transformed, see Table S1, with units of 1/day) in experiment I (see text and Figure 1 for details) for each accession and treatment (combination) as predicted by the linear mixed models fitted to the data on each trait. Within each panel and accession, if two groups share the same letters it implies no significant difference at the 95% confidence level (Tukey correction for multiple comparison was applied). Whiskers indicate standard errors of the slopes.

In experiment II, the leaf blades of Lp2-6 became more elongated when submerged (Figures 4 and S3) with a strong recovery towards control values during post-submergence with or without drought (Figure 5 and S4). Overall, no effect of drought was detected for *blade shape*.

**Figure 4:**
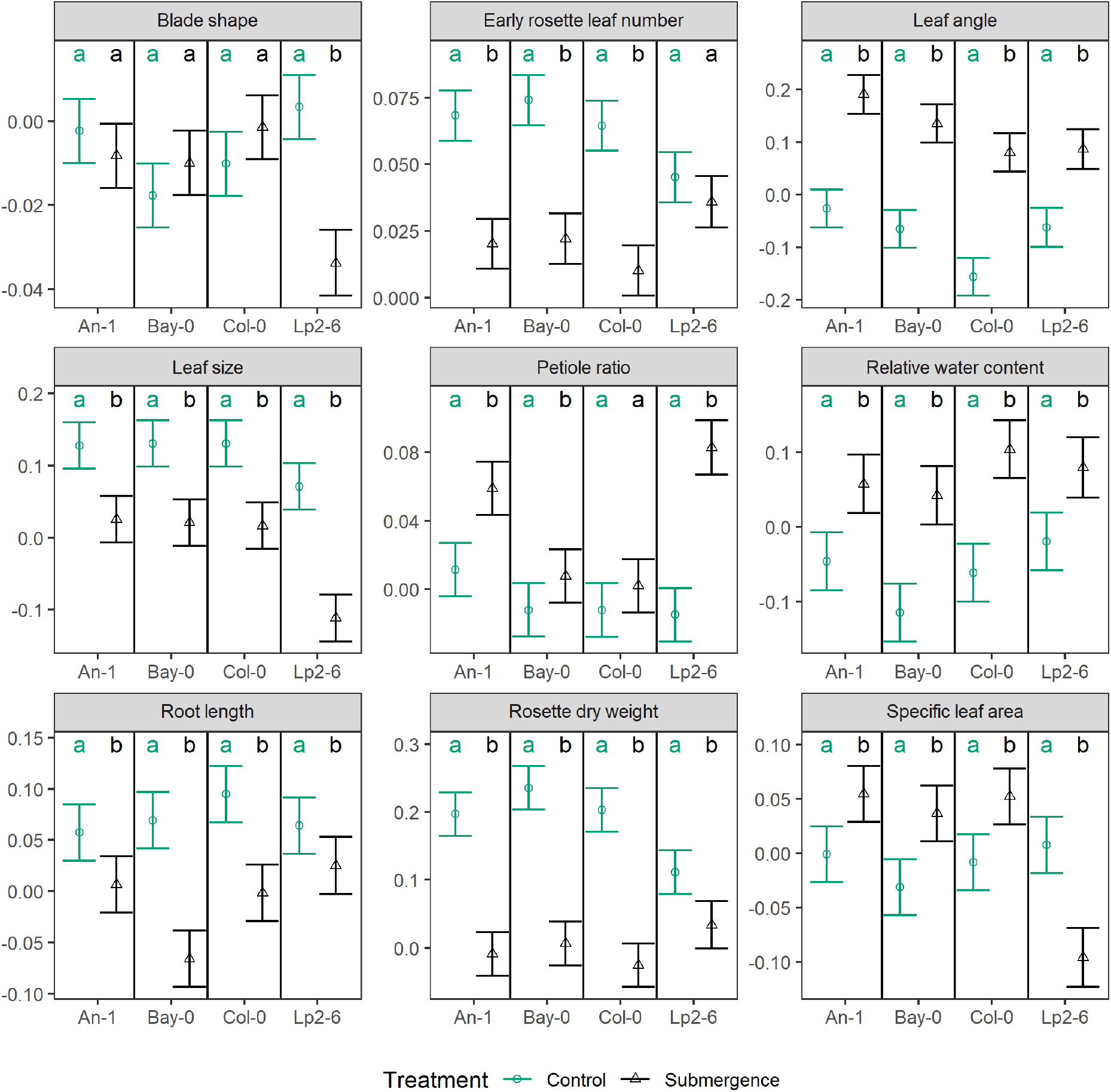
Estimated rate of change over time of each trait (after being transformed, see Table S1, with units of 1/day) in experiment II, phase a (see text and Figure 1 for details) for each accession and treatment as predicted by the linear mixed models fitted to the data on each trait. Within each panel and accession, if two groups share the same letters it implies no statistically significant difference at the 95% confidence level (Tukey’s HSD correction for multiple comparison was applied). Whiskers indicate standard errors of the slopes.

**Figure 5:**
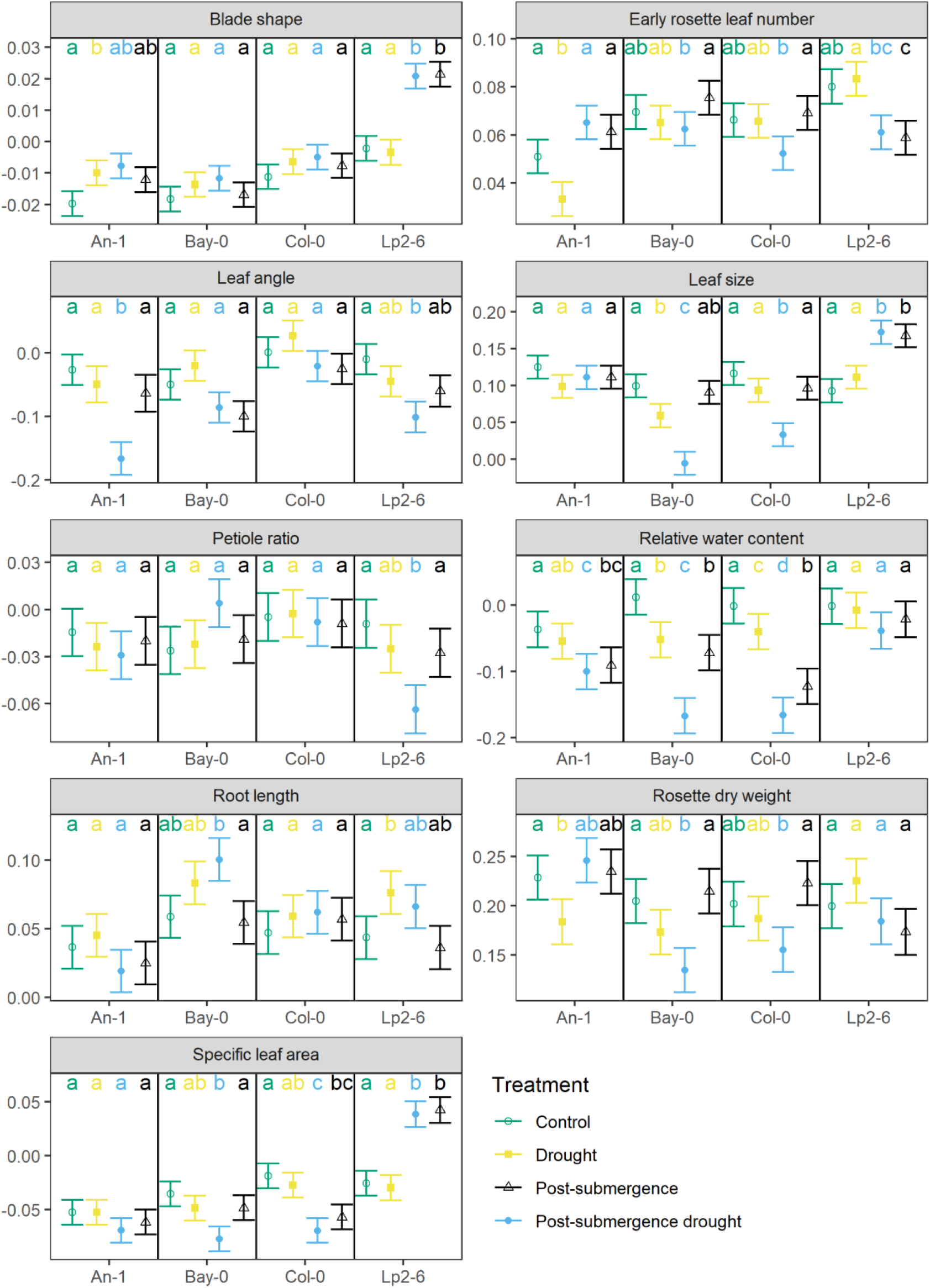
Estimated rate of change over time of each trait (after being transformed, see Table S1, with units of 1/day) in experiment II, phase b (see text and Figure 1 for details) for each accession and treatment as predicted by the linear mixed models fitted to the data on each trait. Within each panel and accession, if two groups share the same letters it implies no statistically significant difference at the 95% confidence level (Tukey’s correction for multiple comparison was applied). Whiskers indicate standard errors of the slopes.

### Flowering time

High temperature (with or without drought) reduced the time to first flower opening (*flowering time*) in all accessions except Lp2-6, while neither submergence nor drought had any effect on this trait (Figure 6). The effect of high temperature varied across the affected accessions, with the smallest effect on An-1 and the largest on Col-0 (Figure 6).

**Figure 6:**
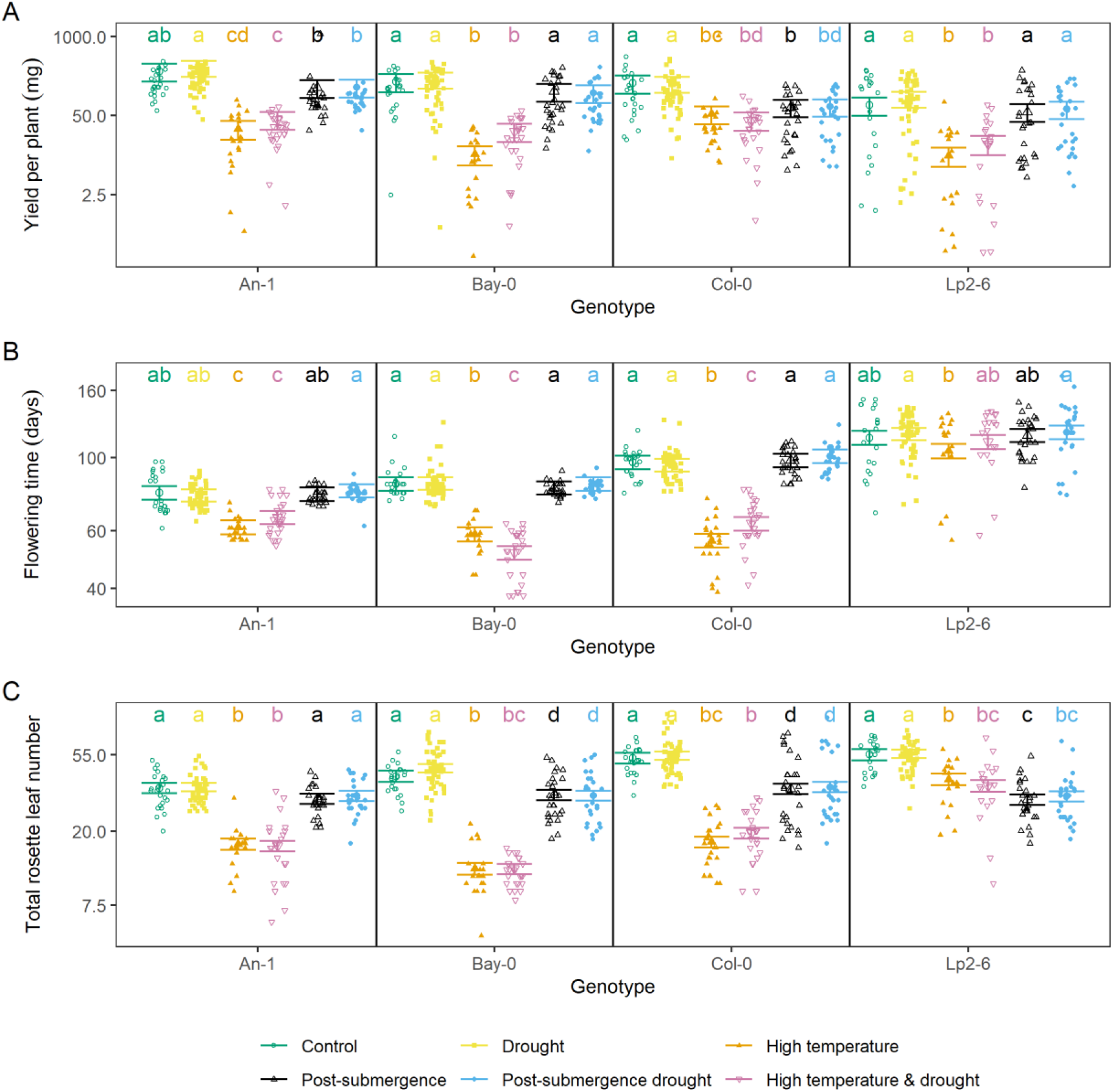
Estimated average values of each trait measured in experiment III for each accession and treatment as predicted by the linear mixed models fitted to the data on each trait. Within each panel and accession, if two groups share the same letters it implies no statistically significant difference at the 95% confidence level (the significant tests were performed in the transformed scale, the means are reported in the original scale, Tukey’s correction for multiple comparison was applied). Whiskers indicate standard errors of the means. Original data points each representing an individual plant are shown as dots.

### Leaf angle

The insertion angle of leaves with respect to the horizontal plane (*leaf angle*) was strongly increased by high temperature (Figures 2 and S5), though no dynamic effect was observed (Figures 3 and S5). Drought did not affect the values of this trait but there was a strong positive interaction with high temperature which led to further increases in *leaf angle* (Figures 3 and S5).

*Leaf angle* also increased during submergence (Figures 4 and S6) and did not decrease towards control values during post-submergence, except when combined with drought for An-1 and Lp2-6 (Figure 5 and S7).

### Leaf size

The average area of a rosette leaf (*leaf size*) of Bay-0 was increased by high temperature (Figures 2 and S8) but the dynamics were not affected (Figures 3 and S8). In contrast, drought always led to smaller leaves in Bay-0 and the effect was enhanced when combined with high temperature (Figure 3).

Submergence also led to smaller leaves in all accessions (Figures 4 and S9) and no recovery was observed during post-submergence except for Lp2-6 (Figures 5 and S10). The application of drought during post-submergence did not affect the dynamics of *leaf size* in Lp2-6 or An-1, but strongly suppressed the trait values in Bay-0 and Col-0 (Figures 5 and S10).

### Petiole ratio

The ratio between petiole length and leaf length (*petiole ratio*) was increased by high temperature in all accessions (Figures 2 and S11) but the dynamics were not affected (Figures 3 and S11). Submergence also led to an increase in *petiole ratio*, especially for An-1 and Lp2-6 (Figures 4 and S12), though the post-submergence recovery phase did not differ from control conditions (Figures 5 and S13). Drought did not affect this trait except when combined with post-submergence where it led to a strong decrease in *petiole ratio* in Lp2-6 (Figure 5).

### Relative water content

The relative water content of leaves (*relative water content*) was not affected by high temperature (Figures 2, 3 and S14), but it decreased with drought (Figures 3, 5, S14 and S16) and the effect of drought was slightly enhanced by combining it with high temperature or post-submergence (Figures 3, 5, S14 and S16). Submergence led to an increase in *relative water content* (Figures 4 and S15), but this was followed by a decrease during post-submergence, except in Lp2-6 (Figures 5 and S16).

### Root length

The maximum depth reached by the root system (*root length*) did not differ between control and high temperature at the 10-leaf stage (Figures 2 and S17), but the stress had a negative effect on the growth of *root length* of Lp2-6 (Figures 3 and S17). *Root length* strongly decreased during submergence, especially for Bay-0 and Col-0 (Figures 4 and S18) but there were no differences noted in its dynamics between control and post-submergence (Figures 5 and S19). There were some effects of drought on *root length* (on its own or when combined with post-submergence) for Bay-0 and Lp2-6, though they were not always reproduced across the experiments (Figures 3, 5, S16 and S19).

### Rosette dry weight

The effect of high temperature on rosette biomass (*rosette dry weight*) at the 10-leaf stage and its rate of increase varied across accessions. Both An-1 and Col-0 had a lower *rosette dry weight* at the 10-leaf stage under high temperature (Figures 2 and S20), but the stress accelerated the accumulation of biomass in An-1, had no dynamic effect on Col-0 and slowed down growth in Lp2-6 (Figures 3 and S20).

The *rosette dry weight* of all accessions decreased under submergence with Lp2-6 being the accession least affected (Figures 4 and S21). However, no differences were detected between post-submergence and control in the rate of *rosette dry weight* increase (Figures 5 and S22).

All accessions continued to accumulate *rosette dry weight* under drought and while this stress reduced growth in Col-0 under experiment I (Figures 3 and S20) and An-1 under experiment II (Figures 5 and S22), these results were not consistent across the two experiments. Drought countered the dynamics effects of high temperature on *rosette dry weight* control (Figures 3 and S20), whereas the rate of *rosette dry weight* increase was strongly suppressed by post-submergence drought in Bay-0 and Col-0 (Figures 5 and S22).

### Rosette leaf number

The number of rosette leaves at each harvest timepoint was counted in all the experiments in this study. In experiment III, the rosette leaf number is referred to as *total rosette leaf number* and it represents the total number of true leaves produced by the plants up to the moment of bolting (used as a measure of fitness), whereas in experiments I and II, it is denoted as *early rosette leaf number* and represents the number of true leaves present in a plant at harvest. The rate of change of the transformed values of *early rosette leaf number* over time is denoted as “rate of leaf appearance”.

High temperature had a positive effect on the rate of leaf appearance of An-1 (Figures 3 and S23). On the other hand, the rate of leaf appearance was decreased by high temperature in Bay-0 and Col-0. The *total rosette leaf number* at bolting was also decreased by high temperature in all accessions except Lp2-6 (Figure 6). The effect of high temperature on Bay-0 was so strong that they bolted before the last harvest of experiment I (Figure S23).

The rate of leaf appearance decreased strongly during submergence in all accessions except Lp2-6 (Figures 4 and S24) but returned to the same rate as in control conditions during post-submergence, except for Lp2-6 where it decreased during post-submergence (Figures 5 and S25). In all accessions, the *total rosette leaf number* was lower in plants that had been submerged, although the effect was much smaller than for high temperature (Figure 6).

Drought did not have any effect on *total rosette leaf number,* or the rate of leaf appearance and no clear interactions were observed between high temperature or post-submergence and drought.

### Specific leaf area

The average specific leaf area of the rosettes (*specific leaf area*) increased slightly for Bay-0 and Col-0 at the 10-leaf stage (Figures 2 and S26). This trait decreased over time as adult leaves had lower *specific leaf area* compared to juvenile leaves and high temperature accelerated this trend in Bay-0 (Figures 3 and S26). A decrease in *specific leaf area* of Bay-0 was also observed for drought (on its own or when combined with the other stresses).

Submergence increased *specific leaf area* in all accessions except for Lp2-6 where it decreased (Figures 4 and S27). On the other hand, there were no differences between control and post-submergence, except for Lp2-6 where a strong increase in *specific leaf area* was observed (Figures 5 and S28). No interactions with drought were observed except for Bay-0 (see above).

### Yield

The seed yield per plant (*yield*) decreased under high temperature for all accessions, with Bay-0 being the most sensitive accession (Figure 6). Submergence also negatively affected the *yield* of An-1 and Col-0, though the effect was smaller than for high temperature (Figure 6). Drought did not have any effect on *yield* nor did it interact with other treatments.

## DISCUSSION

### Effects of high temperature

Our results confirm previous studies on thermomorphogenesis that reported significant increases in *leaf angle*, *petiole ratio* and *specific leaf area* (Zanten et al., 2013, Quint et al., 2016, Ibañez et al., 2017, Vile et al., 2012, Vasseur et al., 2011), though we observed a very weak effect on *specific leaf area* compared to previous studies (Vile et al., 2012, Vasseur et al., 2011). The discrepancy on *specific leaf area* may be due to these previous studies using a higher temperature difference (30 °C – 20 °C) and different developmental stage (first silique shattered).

Less is known about the effects of high temperature on *root length* of Arabidopsis. Although Ibañez et al. (2017) and Hanzawa et al. (2013) observed an increase in *root length* under high temperature, these measurements were performed on young seedlings rather than established plants. On the other hand, Vile et al. (2012) reported no effect of high temperature on biomass allocation of roots which would be in agreement with our results.

Despite a relatively small impact on the dry weight of the young rosettes in experiment I (Figure 3), the high temperature treatment had a negative effect on *yield* and *total rosette leaf number* (Figure 6). Such a reduction could not always be explained by a shorter vegetative cycle only, as there was no effect of high temperature on the *flowering time* (measured in days) of Lp2-6 but still *yield* was strongly affected in this accession (Figure 6).

Besides the expected high temperature phenotype, our measurements also revealed that high temperature also reduces the rate of leaf appearance for some accessions (Figure 3), in contradiction with previous experiments in Arabidopsis (Granier et al., 2002) and the thermal time paradigm (Trudgill et al., 2005). Since the leaf appearance rate in Arabidopsis and other dicots is known to be affected by the daily light integral when plants are grown at low light intensities (Chenu et al., 2005, Savvides et al., 2014), we speculate that the temperature optimum for leaf appearance may also depend on the light intensity or photoperiod, but this needs to be tested in future experiments.

### Effects of submergence

The present study provides an extensive documentation of the effect of submergence on a suite of traits in Arabidopsis. The increase in leaf angle and petiole elongation under submergence has been reported previously for flood-adapted species with aerenchyma-rich tissues, such as species from the genus *Rumex* and *Rorippa* (Pierik et al., 2005, Sasidharan and Voesenek, 2015, Cox et al., 2003) as well as in Arabidopsis, which lacks aerenchyma (van Veen et al., 2016). However, unlike most previous submergence research on Arabidopsis, our submergence treatments were performed in the light and the decline in internal oxygen concentrations might have been more gradual, as light availability during submergence allows for some underwater photosynthesis and supports oxygen production and carbon assimilation, making the stress milder than in the absence of light (Vashisht et al., 2011). This may also explain why we do not see a strong evidence of a physiological drought during the post-submergence phase in our experiments, in contrast to what has been reported previously (Sasidharan and Voesenek, 2015, Tamang and Fukao, 2015, Yeung et al., 2019, Fukao et al., 2011).

The significant negative effect of submergence on *yield* for An-1 and Col-0 and on *total rosette leaf number* for all accessions was unexpected (Figure 6). Such an effect could not be explained by the direct impact of submergence on *rosette dry weight* or *leaf initiation rate* (Figure 4), nor by a post-submergence effect, as experiment II revealed no differences between control conditions and the post-submergence recovery of *rosette dry weight* or *early rosette leaf number* (Fig. 4). This implies that the negative effects of submergence on the growth and development of the plants were either delayed (from the moment of de-submergence) or became stronger over time and hence could not be detected during the first nine days of post-submergence that experiment II covered.

A decrease in *total rosette leaf number* but no effect on *flowering time* (as was the case for submergence, Figure 6) is also not common in Arabidopsis. There are examples of experimental manipulations that either increase flowering time and total number of leaves, such as the application of nitric oxide (He et al., 2004) or sucrose (Ohto et al., 2001), or reduce flowering time and total number of leaves, such as vernalization (Martinez-Zapater and Somerville, 1990) or gibberellic acid (Bagnall, 1992). This correlation between *flowering time* and *total rosette leaf number* is also observed across accessions of Arabidopsis and in mutants that affect either of the processes (Méndez-Vigo et al., 2010), suggesting that the average rate of leaf appearance is highly conserved. One example of decoupling the two traits is the application of nitrogen dioxide, which reduces *flowering time* but does not affect *total rosette leaf number* (Takahashi and Morikawa, 2014). However, to our knowledge, such decoupling has not been reported before for an abiotic stress in Arabidopsis.

In terms of *rosette dry weight* and *early rosette leaf number*, Lp2-6 appears to be more tolerant to submergence stress than the other accessions (Figure 4). This is in accordance with previous studies characterizing its relatively high survival and recovery following submergence in darkness (Vashisht et al. 2011; Yeung et al., 2018). The minimal submergence effects on Lp2-6 biomass coincided with a deviant effect (compared to other accessions) on many of the traits such as *blade shape*, *leaf size* or *petiole ratio* (Figure 4), followed by a strong recovery towards control levels during post-submergence (Figure 5). Since these traits were averaged at the plant level, these rapid changes observed in Lp2-6 (Figures 4 and 5) reflect a strong contrast between the morphology of leaves generated during submergence vs non-submerged conditions. Specifically, Lp2-6 grew more elongated, thinner leaves with a longer petiole and a smaller blade, that would have reduced the distance to the water surface, relative to the other accessions. This could result in more access to light and oxygen and hence enable some growth and development under water. Interestingly, Lp2-6 did not increase its *leaf angle* during submergence as much as other accessions. These results suggest that *blade shape*, *leaf size* and *petiole ratio* are traits of interest when evaluating the tolerance to submergence of Arabidopsis.

### Effects of drought and its interactions with high temperature and post-submergence

Overall, drought tended to have weaker (or no) effect on the different traits in our study and had no effect on yield (thus, plants were able to fully recover from the drought stress). Indeed, the drought stress imposed was only mild, as dry weight accumulation during the drought was barely affected in most cases, despite the gravimetric pellet water content decreasing by 75% after 9 days of no watering (Figure S1). However, it is possible that the small transpiration rates of the young Arabidopsis rosettes and the structure of the coconut fibers in the pellets allowed plants to maintain a relatively favorable hydraulic status under such conditions.

Despite the overall weak effects of drought, we identified several instances where drought interacted in a complex manner with high temperature or post-submergence, especially when variation across accessions is considered. Specifically, there were statistically significant interactions between drought and high temperature (for at least one accession) in 9 out of 12 traits (Table S2) and between post-submergence and drought for half of the traits (Table S4). For example, drought increased the effects of high temperature and post-submergence on *leaf angle* for most accessions (Figure 3 and 5). Similarly, post-submergence drought led to small *leaf size* in Bay-0 and Col-0 even though drought and post-submergence had little effect on their own (Figure 5). Vile et al. (2012) reported mostly additive effects of mild drought and high temperature combinations for similar traits to the ones in our experiment, but they also identified some interactions. Strong interactions have also been reported for molecular and metabolic traits in response to combinations of stresses (Mittler, 2006, Rizhsky et al., 2004, Suzuki et al., 2014).

### Conclusions

All tested sub-lethal abiotic stresses and their combinations affected a wide range of traits in Arabidopsis. Moreover, each stress led to a different pattern of effects reflecting the distinct challenges imposed by the different stresses, but there were differences among accessions for some of the traits. Also, there were interactions between drought and other treatments and, when present, the interactions tended to be accession specific.

This study contributes with a rich characterization of the response of Arabidopsis to sub-lethal stresses at the level of organ and plant, providing a starting point for further in-depth physiological research and mechanistic modelling efforts. Mechanistic plant models aid in understanding the feedbacks and trade-offs acting on plant growth under different abiotic stresses, but only if they are well calibrated and validated against a comprehensive quantitative description of plant physiology, morphology and development such as the one provided in this study.

## ACKNOWLEDGEMENTS

We thank Ankie Ammerlaan, Jolanda Schuurmans and Evelien Stouten for their excellent technical support during the experiments.

