## Supplementary material for "Effects of sub-lethal single, simultaneous, and sequential abiotic stresses on phenotypic traits of Arabidopsis thaliana": Supplemetary Figures and Tables

### Supplemental tables

**Table S1:** For each trait and experiment (I-II), the type of non-linear transformation of the response variable in the model (Transform), total number of measurements for a given trait and experiment ($N_{total}$), total number of batches used for a given experiment ($N_{batch}$), and average, standard deviation, maximum and minimum number of biological replicates for each combination of accession, treatment, batch and timepoint ($\overline{N}$ and $\sigma_N$, $N_{max}$ and $N_{min}$, respectively) are indicated.

| Trait | Transform* | $N_{total}$ | $N_{batch}$ | $\overline{N}$ | $\sigma_N$ | $N_{max}$ | $N_{min}$ |
|---|---|---|---|---|---|---|---|
| **Experiment I** | | | | | | | |
| ***Blade shape*** | log | 478 | 8 | 6.0 | 0.2 | 6 | 5 |
| ***Early rosette leaf number*** | log | 478 | 8 | 6.0 | 0.2 | 6 | 5 |
| ***Leaf angle*** | logit | 458 | 8 | 5.7 | 0.7 | 6 | 3 |
| ***Leaf size*** | log | 478 | 8 | 6.0 | 0.2 | 6 | 5 |
| ***Petiole ratio*** | logit | 478 | 8 | 6.0 | 0.2 | 6 | 5 |
| ***Relative water content*** | logit | 477 | 8 | 6.0 | 0.3 | 6 | 3 |
| ***Root length*** | log | 470 | 8 | 5.9 | 0.4 | 6 | 4 |
| ***Rosette dry weight*** | log | 477 | 8 | 6.0 | 0.3 | 6 | 3 |
| ***Specific leaf area*** | log | 475 | 8 | 5.9 | 0.4 | 6 | 3 |
| **Experiment II (phase a)** | | | | | | | |
| ***Blade shape*** | log | 239 | 8 | 6.0 | 0.2 | 6 | 5 |
| ***Early rosette leaf number*** | log | 239 | 8 | 6.0 | 0.2 | 6 | 5 |
| ***Leaf angle*** | logit | 237 | 8 | 5.9 | 0.3 | 6 | 5 |
| ***Leaf size*** | log | 239 | 8 | 6.0 | 0.2 | 6 | 5 |
| ***Petiole ratio*** | logit | 239 | 8 | 6.0 | 0.2 | 6 | 5 |
| ***Relative water content*** | logit | 235 | 8 | 5.9 | 0.4 | 6 | 4 |
| ***Root length*** | log | 239 | 8 | 6.0 | 0.2 | 6 | 5 |
| ***Rosette dry weight*** | log | 236 | 8 | 5.9 | 0.4 | 6 | 4 |
| ***Specific leaf area*** | log | 236 | 8 | 5.9 | 0.4 | 6 | 4 |
| **Experiment II (phase b)** | | | | | | | |
| ***Blade shape*** | log | 467 | 8 | 6.0 | 0.1 | 6 | 5 |
| ***Early rosette leaf number*** | log | 467 | 8 | 6.0 | 0.1 | 6 | 5 |
| ***Leaf angle*** | logit | 460 | 8 | 5.9 | 0.4 | 6 | 4 |
| ***Leaf size*** | log | 467 | 8 | 6.0 | 0.1 | 6 | 5 |
| ***Petiole ratio*** | logit | 467 | 8 | 6.0 | 0.1 | 6 | 5 |
| ***Relative water content*** | logit | 474 | 8 | 5.9 | 0.3 | 6 | 4 |
| ***Root length*** | log | 476 | 8 | 6.0 | 0.2 | 6 | 5 |
| ***Rosette dry weight*** | log | 476 | 8 | 5.9 | 0.3 | 6 | 4 |
| ***Specific leaf area*** | log | 464 | 8 | 6.0 | 0.3 | 6 | 4 |
| **Experiment III** | | | | | | | |
| ***Flowering time*** | log | 678 | 3 | 9.4 | 3.9 | 19 | 3 |
| ***Total rosette leaf number*** | log | 678 | 3 | 9.4 | 3.9 | 19 | 3 |
| ***Yield*** | log | 678 | 3 | 9.4 | 3.9 | 19 | 3 |

* log stands for a natural logarithmic (i.e. $\log_e(x)$) transformation where x is the trait value; logit stands for a $\log_e[x/(1 – x)]$ transformation.

**Table S2:** Significance of F tests expressed as $\log_{10}(P)$ on the effects of the accession (A), high temperature (HT) and drought (D) and their interactions (denoted with ":") on either the intercepts (initial value) or the slopes (rate of change) of the linear mixed models fitted to the measured traits in experiment I (see text and Figure 1 for details). P values that are lower than 0.05 (i.e., $\log_{10}(P) < -1.3$) are highlighted in bold.

| Traits | Initial value | | | Rate of change | | | | | | |
|---|---|---|---|---|---|---|---|---|---|---|
| | A | HT | HT:A | G | HT | D | HT:D | HT:G | D:A | HT:D:A |
| ***Blade shape*** | -0.6 | **-9.2** | **-13.3** | -0.6 | **-7.4** | 0.0 | -0.5 | **-3.1** | 0.0 | -0.5 |
| ***Early rosette leaf number*** | -0.5 | **-14.0** | **-9.1** | 0.0 | **-3.4** | -0.4 | -0.4 | **-6.0** | 0.0 | -0.6 |
| ***Leaf angle*** | -0.2 | **-4.3** | -0.1 | -0.3 | 0.0 | **-1.6** | **-2.7** | -0.6 | -1.2 | -0.3 |
| ***Leaf size*** | -0.2 | -0.6 | **-3.3** | -0.3 | -0.4 | -0.5 | **-2.4** | -0.8 | -0.8 | **-1.9** |
| ***Petiole ratio*** | -0.8 | **-16.0** | **-9.4** | **-2.4** | **-1.7** | -0.4 | **-1.5** | -0.1 | -1.1 | **-2.0** |
| ***Relative water content*** | -0.2 | **-2.6** | -0.4 | -0.5 | -0.2 | **-2.1** | **-1.6** | 0.0 | **-2.1** | -0.5 |
| ***Root length*** | -0.2 | -0.6 | -1.1 | -0.3 | -0.2 | -1.3 | -0.6 | **-1.4** | **-1.7** | **-1.5** |
| ***Rosette dry weight*** | 0.0 | **-15.4** | **-10.1** | -0.7 | **-1.6** | -0.3 | -0.6 | **-2.2** | -0.6 | **-1.3** |
| ***Specific leaf area*** | 0.0 | -1.2 | -1.0 | -0.4 | **-1.5** | -0.3 | **-1.5** | **-3.4** | -1.0 | **-1.4** |

**Table S3**: Significance of F tests expressed as $\log_{10}(P)$ on the effects of the accession (A), submergence (S) and their interactions (denoted with ":") on either the intercepts (initial value) or the slopes (rate of change) as captured by the linear mixed models fitted to the measured traits in experiment II - phase a (see text and Figure 1 for details). P values that are lower than 0.05 (i.e., $\log_{10}(P) < -1.3$) are highlighted in bold.

| Trait | Initial values | Rate of change | | |
|---|---|---|---|---|
| | A | A | S | S:A |
| ***Blade shape*** | **-1.6** | -0.6 | -0.4 | **-5.1** |
| ***Early rosette leaf number*** | **-2.2** | -0.8 | **-7.5** | **-3.3** |
| ***Leaf angle*** | 0.0 | -1.1 | **-8.8** | -0.5 |
| ***Leaf size*** | -0.2 | -0.3 | **-5.1** | **-1.3** |
| ***Petiole ratio*** | -0.1 | -0.2 | **-5.7** | **-8.2** |
| ***Relative water content*** | -0.4 | -0.4 | **-4.0** | -0.8 |
| ***Root length*** | -1.0 | -0.1 | **-2.3** | **-3.3** |
| ***Rosette dry weight*** | -0.4 | **-1.4** | **-10.3** | **-2.9** |
| ***Specific leaf area*** | -0.1 | -0.1 | **-2.1** | **-7.9** |

**Table S4:** Significance of F tests expressed as $\log_{10}(P)$ on the effects of the accession (A), post-submergence (PS), drought (D) and their interactions (denoted with ":") on either the intercepts (initial value) or the slopes (rate of change) as captured by the linear mixed models fitted to the measured traits in experiment II – phase b (see text and Figure 1 for details). P values that are lower than 0.05 (i.e., $\log_{10}(P) < -1.3$) are highlighted in bold.

| Trait | Initial value | | | Rate of change | | | | | | |
|---|---|---|---|---|---|---|---|---|---|---|
| | A | PS | PS:A | A | PS | D | PS:D | PS:A | D:A | PS:D:A |
| ***Blade shape*** | **-1.7** | -0.9 | **-9.6** | **-2.2** | -1.0 | **-2.9** | -0.7 | **-2.4** | -1.0 | -0.1 |
| ***Early rosette leaf number*** | **-2.0** | **-14.6** | **-6.2** | **-1.5** | -0.7 | **-3.5** | **-2.7** | **-1.7** | **-1.7** | **-3.0** |
| ***Leaf angle*** | -0.8 | **-7.8** | -0.3 | -0.3 | -0.5 | -0.4 | **-1.6** | 0.0 | -1.1 | -0.4 |
| ***Leaf size*** | -0.3 | **-11.1** | **-1.9** | -0.3 | -0.3 | **-1.6** | -0.9 | **-3.2** | **-2.3** | **-2.6** |
| ***Petiole ratio*** | -0.2 | **-2.1** | **-2.1** | -0.1 | -0.1 | -0.4 | 0.0 | -0.1 | -0.3 | -0.5 |
| ***Relative water content*** | -0.1 | **-9.4** | **-2.7** | -0.2 | **-2.6** | -0.9 | -0.2 | **-3.3** | **-2.8** | -0.4 |
| ***Root length*** | -1.0 | **-4.7** | **-4.7** | -0.1 | -0.3 | -0.3 | -0.4 | -0.1 | -0.4 | -0.4 |
| ***Rosette dry weight*** | -0.8 | **-16.0** | **-3.8** | -0.1 | -0.1 | **-2.2** | **-1.8** | -0.2 | **-1.9** | **-2.6** |
| ***Specific leaf area*** | -0.3 | **-7.4** | **-11.9** | -0.7 | -0.3 | 0.0 | -0.3 | **-5.8** | -0.1 | -0.1 |

**Table S5:** Significance of F tests expressed as $\log_{10}(P)$ on the effects of the accession (A), high temperature (HT), post-submergence (PS), drought (D) and their interactions (denoted with ":") as captured by the linear mixed models fitted to the measured traits in experiment III (see text and Figure 1 for details). P values that are lower than 0.05 (i.e., $\log_{10}(P) < -1.3$) are highlighted in bold.

| Trait | A | HT | PS | D | HT:D | PS:D | HT:A | PS:A | D:A | HT:D:A | PS:D:A |
|---|---|---|---|---|---|---|---|---|---|---|---|
| ***Flowering time*** | **-16.0** | **-9.4** | -0.1 | -0.2 | -1.1 | -0.4 | **-14.8** | -0.1 | -0.2 | **-3.2** | -0.1 |
| ***Total rosette leaf number*** | **-7.1** | **-16.0** | -1.1 | -0.1 | -0.1 | -0.1 | **-12.1** | **-3.3** | -0.1 | -0.2 | -0.1 |
| ***Yield*** | **-3.5** | **-12.0** | **-1.6** | -0.2 | -0.2 | -0.1 | **-2.6** | -0.4 | 0.0 | -0.5 | 0.0 |

### Supplemental figures

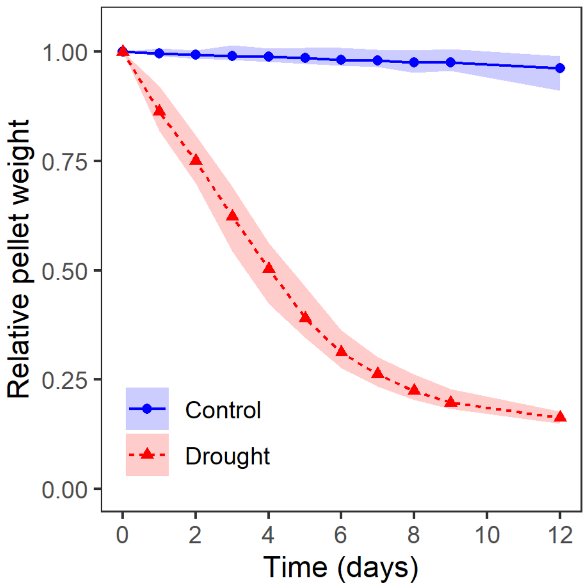


**Figure S1:** Relative pellet weight (with respect to the first timepoint, Time = 0) over time under control (well-watered) and progressive drought conditions (no watering). Lines and symbols indicate the estimated mean relative weight from multiple pellets and the shaded areas represent the 95% confidence intervals of the mean trend.

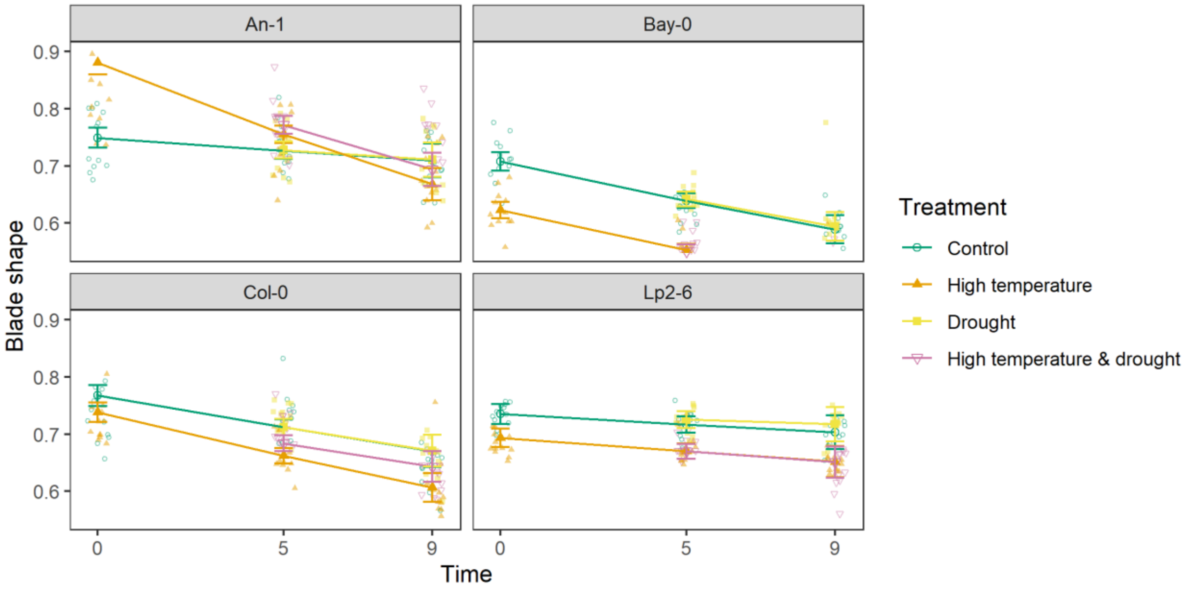


**Figure S2:** Measured blade shape in experiment I and estimated means for each combination of accession (panel), treatment (dots and lines) and time (days) using the fitted linear mixed models. Whiskers represent standard errors of the means, dots in the background indicate individual measurements. Time = 0 corresponds to the 10 true leaf stage where treatments were initiated.

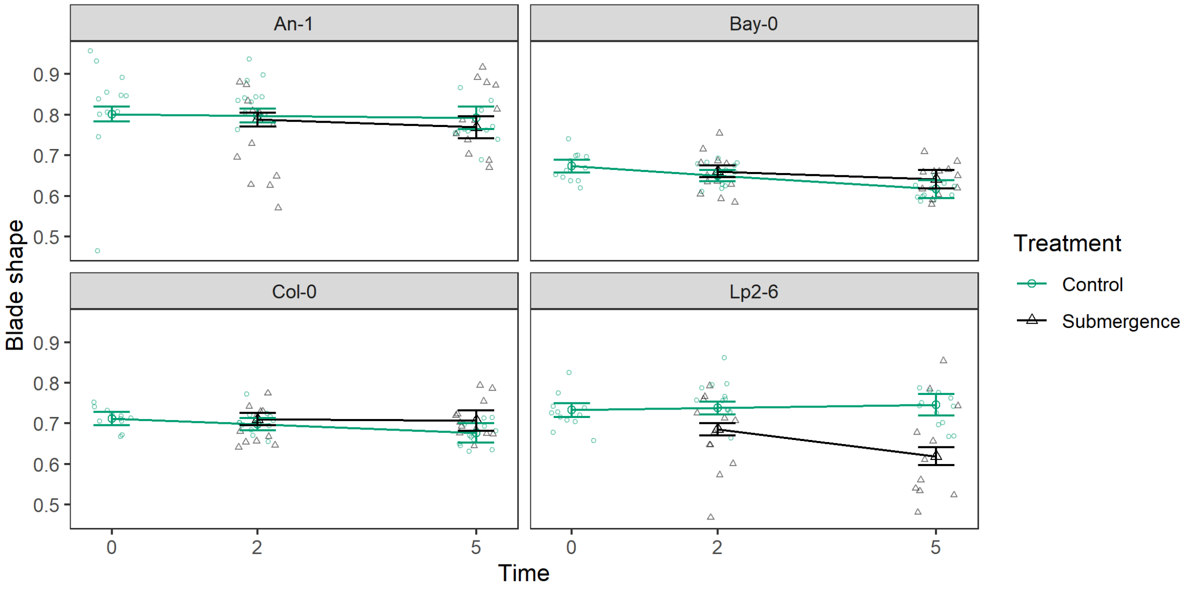


**Figure S3:** Measured blade shape in experiment II (phase a) and estimated means for each combination of accession (panel), treatment (dots and lines) and time (days) using the fitted linear mixed models. Whiskers represent standard errors of the means, dots in the background indicate individual measurements. Time = 0 corresponds to the 10 true leaf stage where treatments were initiated.

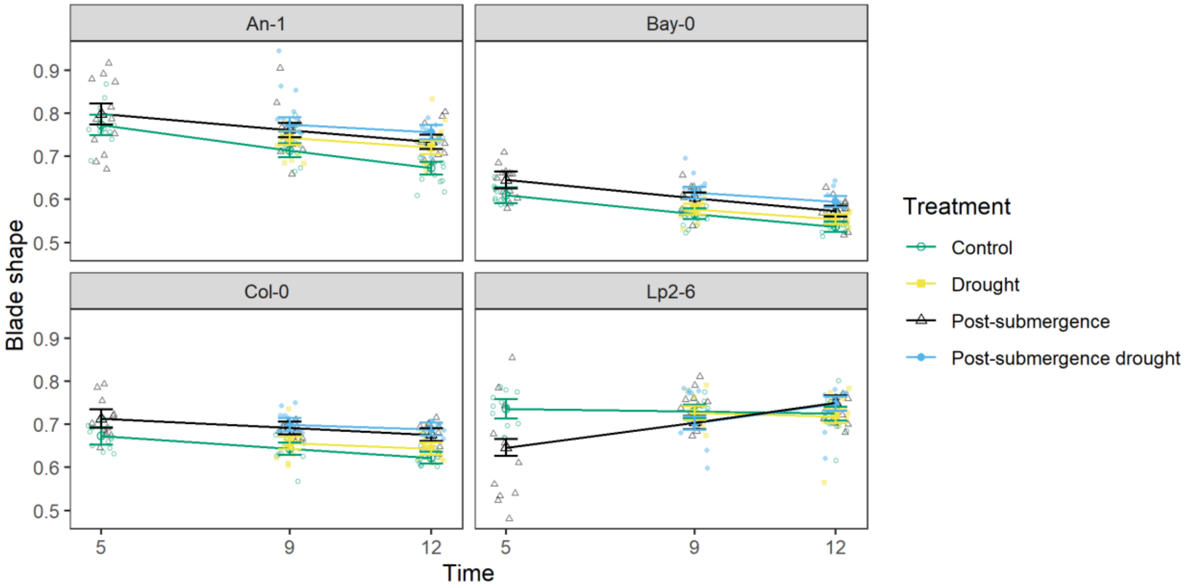


**Figure S4:** Measured blade shape in experiment II (phase b) and estimated means for each combination of accession (panel), treatment (dots and lines) and time (days) using the fitted linear mixed models. Whiskers represent standard errors of the means, dots in the background indicate individual measurements. Time = 5 corresponds to 5 days after the 10 true leaf stage was reached.

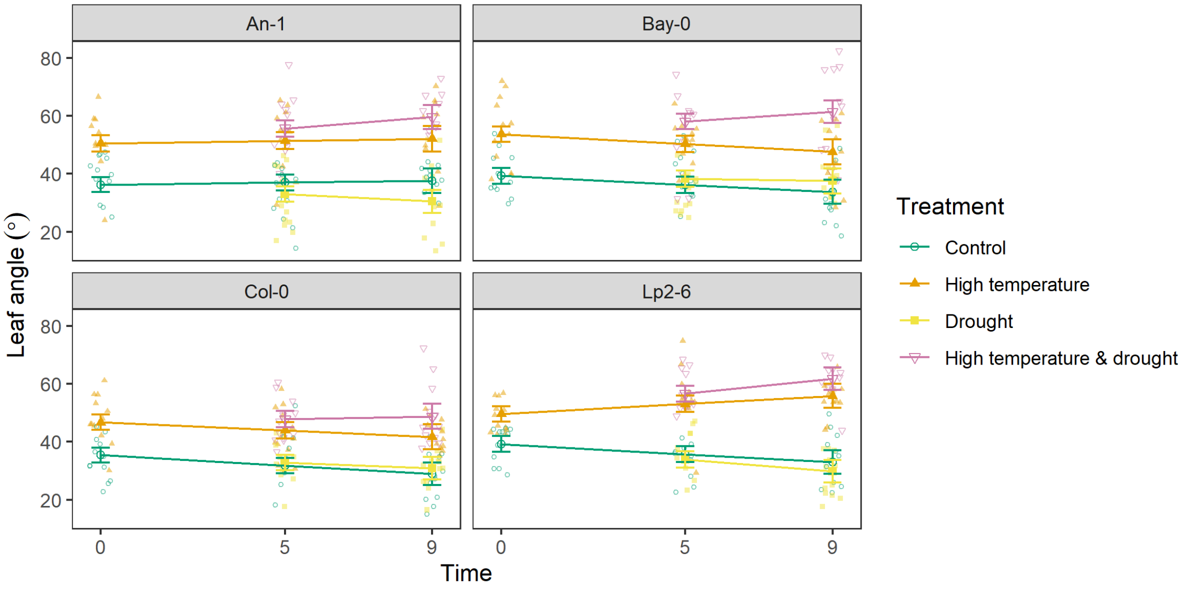


**Figure S5**: Measured leaf angle in experiment I and estimated means for each combination of accession (panel), treatment (dots and lines) and time (days) using the fitted linear mixed models. Whiskers represent standard errors of the means, dots in the background indicate individual measurements. Time = 0 corresponds to the 10 true leaf stage where treatments were initiated.

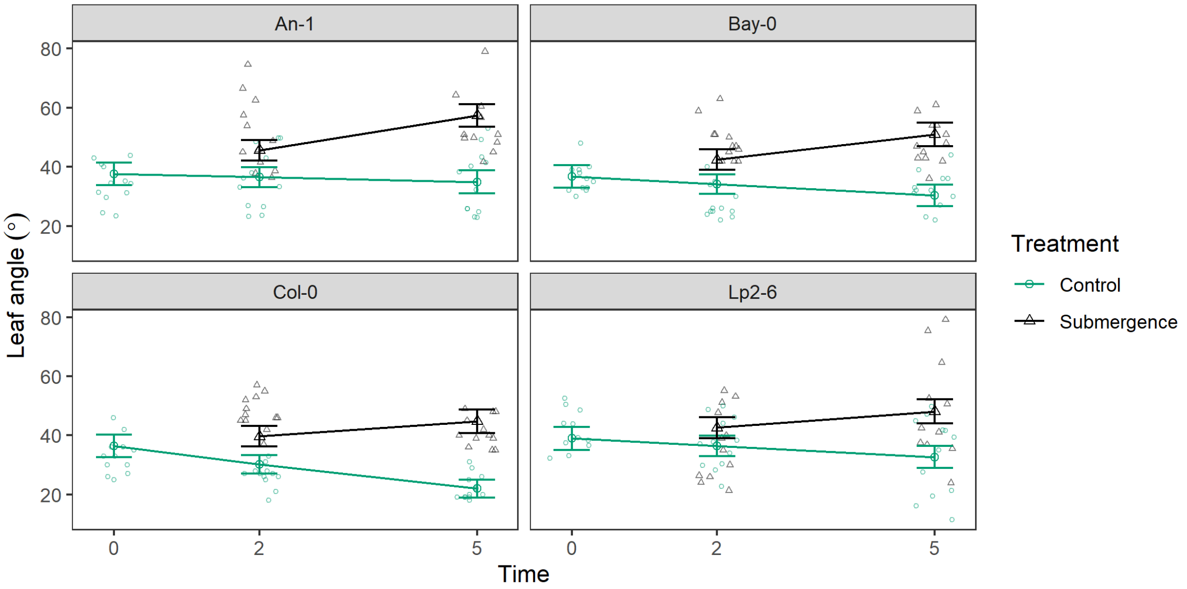


**Figure S6:** Measured leaf angle in experiment II (phase a) and estimated means for each combination of accession (panel), treatment (dots and lines) and time (days) using the fitted linear mixed models. Whiskers represent standard errors of the means, dots in the background indicate individual measurements. Time = 0 corresponds to the 10 true leaf stage where treatments were initiated.

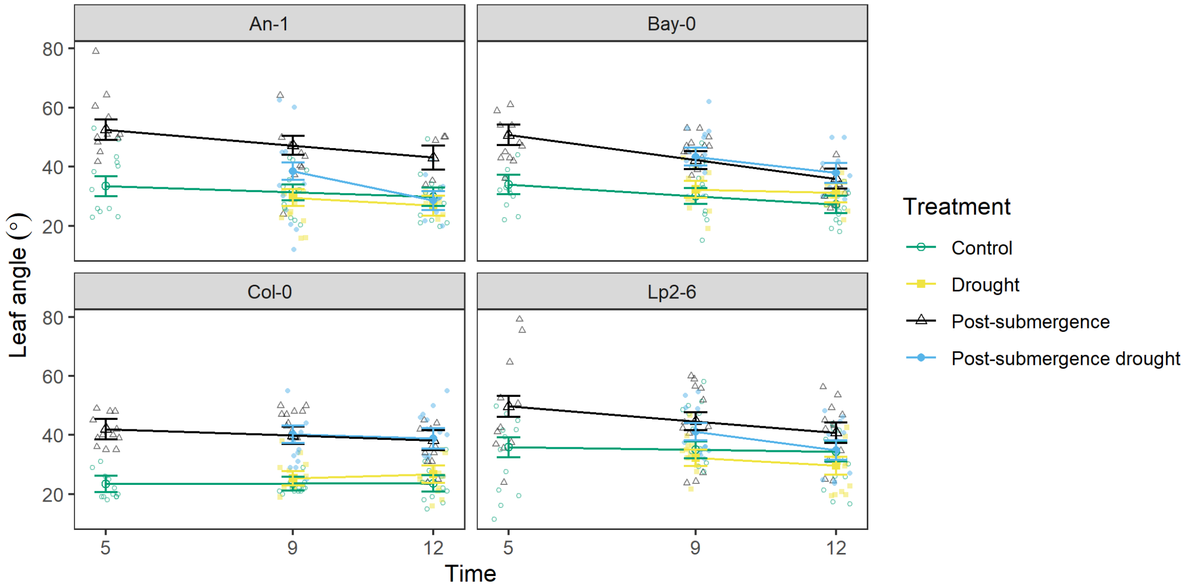


**Figure S7:** Measured leaf angle in experiment II (phase b) and estimated means for each combination of accession (panel), treatment (dots and lines) and time (days) using the fitted linear mixed models. Whiskers represent standard errors of the means, dots in the background indicate individual measurements. Time = 5 corresponds to 5 days after the 10 true leaf stage was reached.

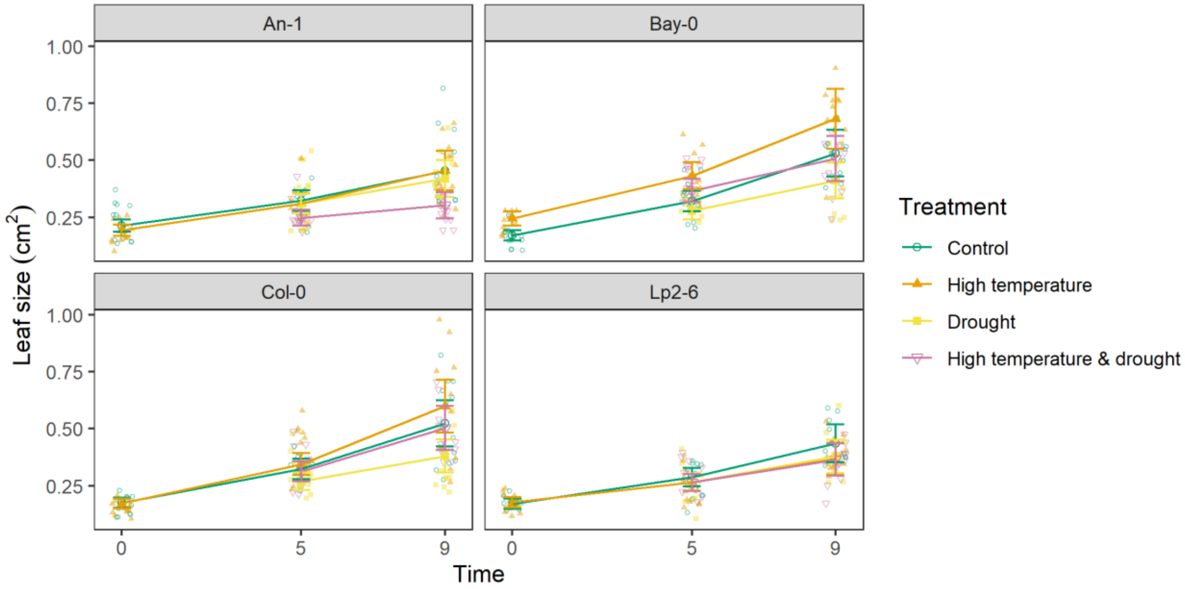


**Figure S8:** Measured leaf size in experiment I and estimated means for each combination of accession (panel), treatment (dots and lines) and time (days) using the fitted linear mixed models. Whiskers represent standard errors of the means, dots in the background indicate individual measurements. Time = 0 corresponds to the 10 true leaf stage where treatments were initiated.

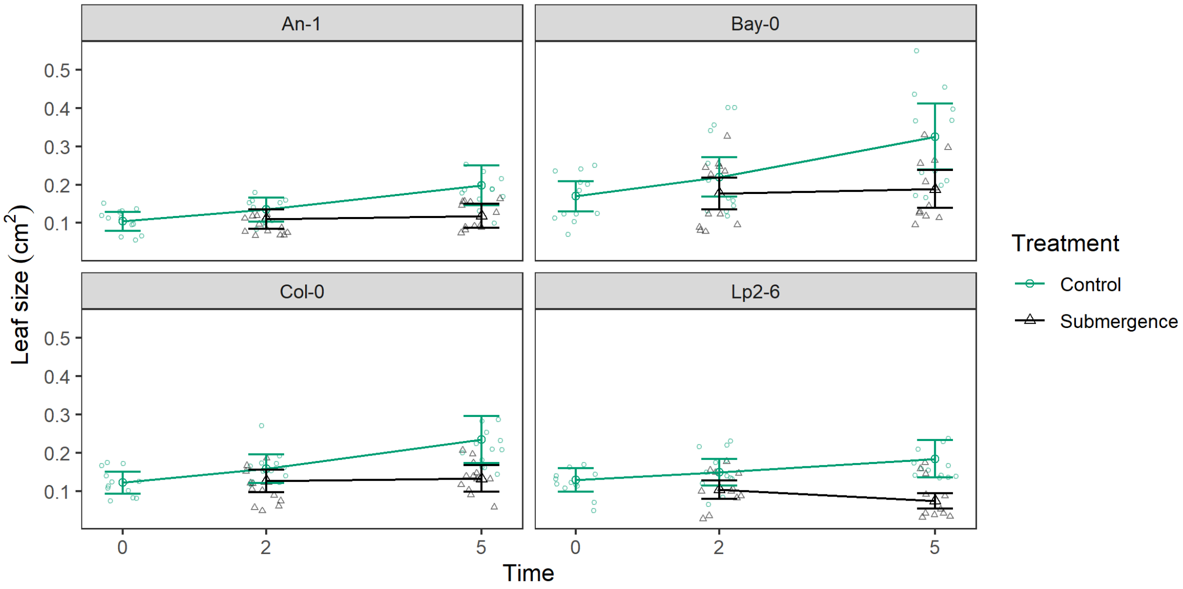


**Figure S9:** Measured leaf size in experiment II (phase a) and estimated means for each combination of accession (panel), treatment (dots and lines) and time (days) using the fitted linear mixed models. Whiskers represent standard errors of the means, dots in the background indicate individual measurements. Time = 0 corresponds to the 10 true leaf stage where treatments were initiated.

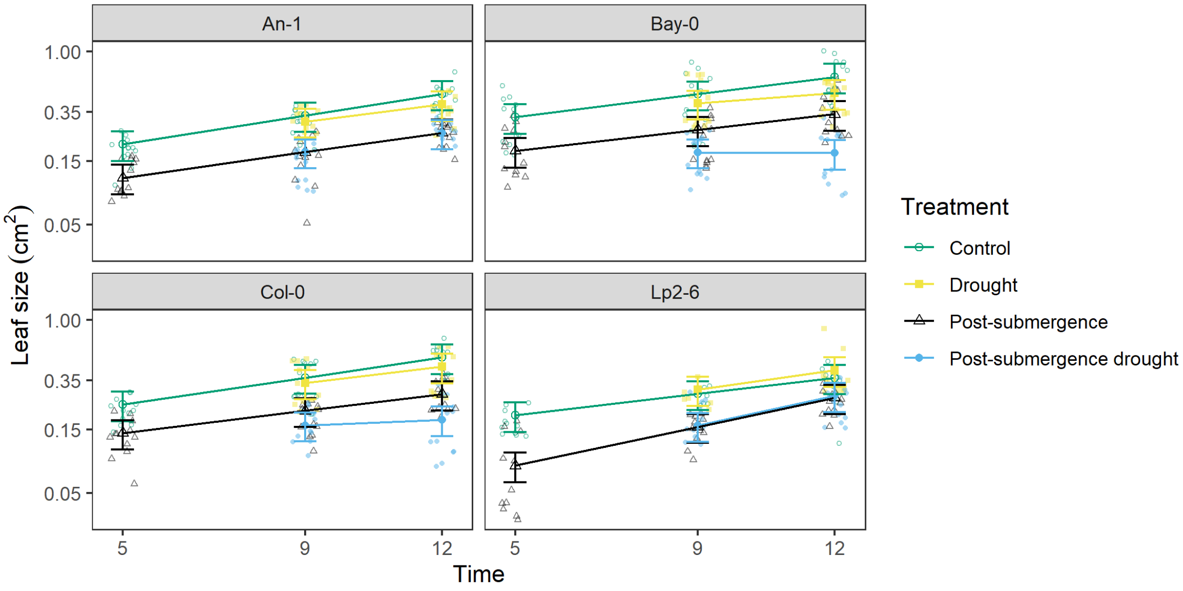


**Figure S10:** Measured leaf size in experiment II (phase b) and estimated means for each combination of accession (panel), treatment (dots and lines) and time (days) using the fitted linear mixed models. Whiskers represent standard errors of the means, dots in the background indicate individual measurements. Time = 5 corresponds to 5 days after the 10 true leaf stage was reached.

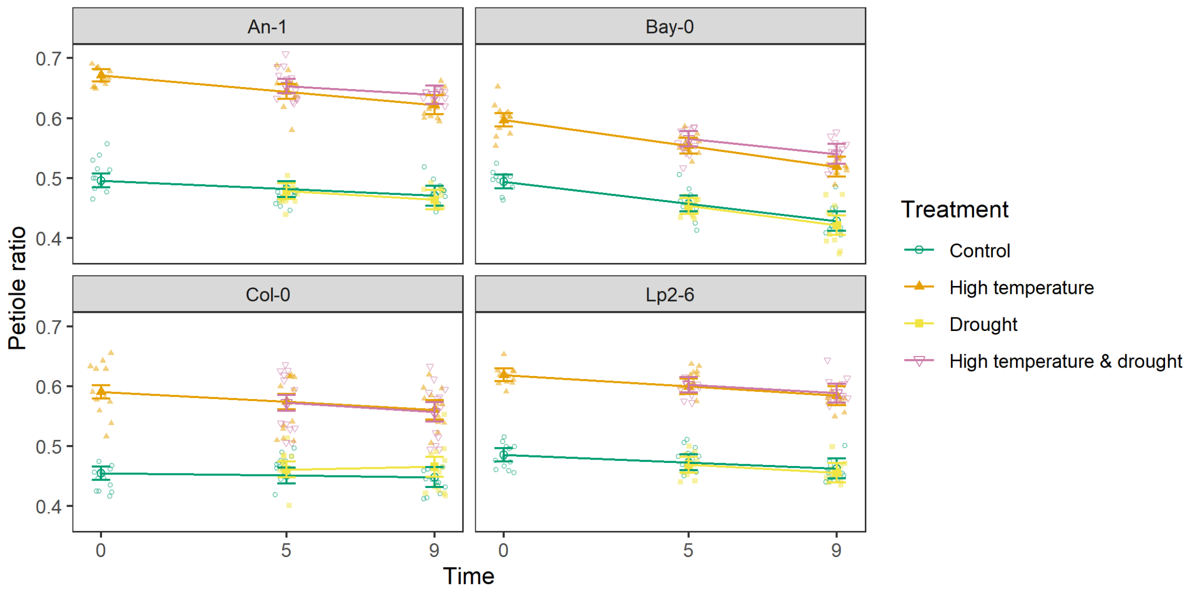


**Figure S11:** Measured petiole ratio in experiment I and estimated means for each combination of accession (panel), treatment (dots and lines) and time (days) using the fitted linear mixed models. Whiskers represent standard errors of the means, dots in the background indicate individual measurements. Time = 0 corresponds to the 10 true leaf stage where treatments were initiated.

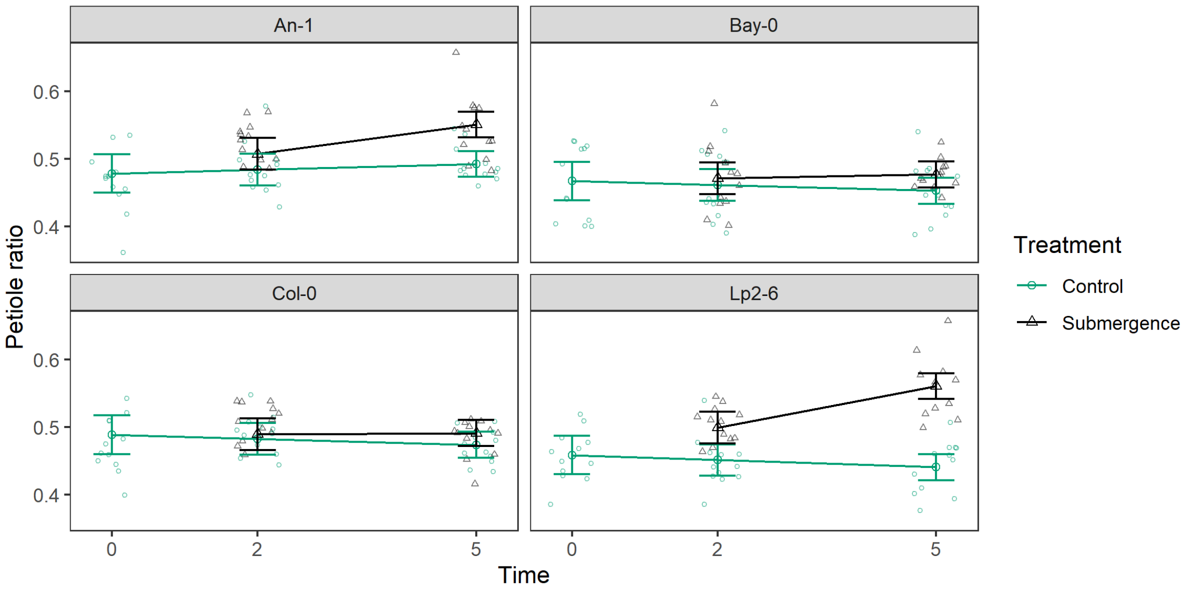


**Figure S12:** Measured petiole ratio in experiment II (phase a) and estimated means for each combination of accession (panel), treatment (dots and lines) and time (days) using the fitted linear mixed models. Whiskers represent standard errors of the means, dots in the background indicate individual measurements. Time = 0 corresponds to the 10 true leaf stage where treatments were initiated.

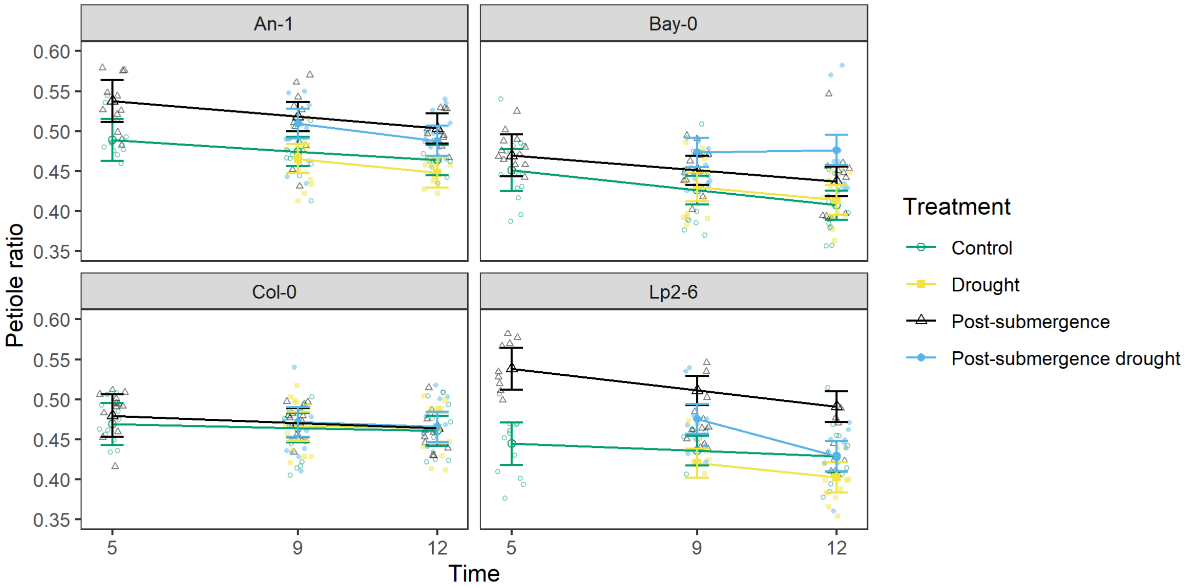


**Figure S13:** Measured petiole ratio in experiment II (phase b) and estimated means for each combination of accession (panel), treatment (dots and lines) and time (days) using the fitted linear mixed models. Whiskers represent standard errors of the means, dots in the background indicate individual measurements. Time = 5 corresponds to 5 days after the 10 true leaf stage was reached.

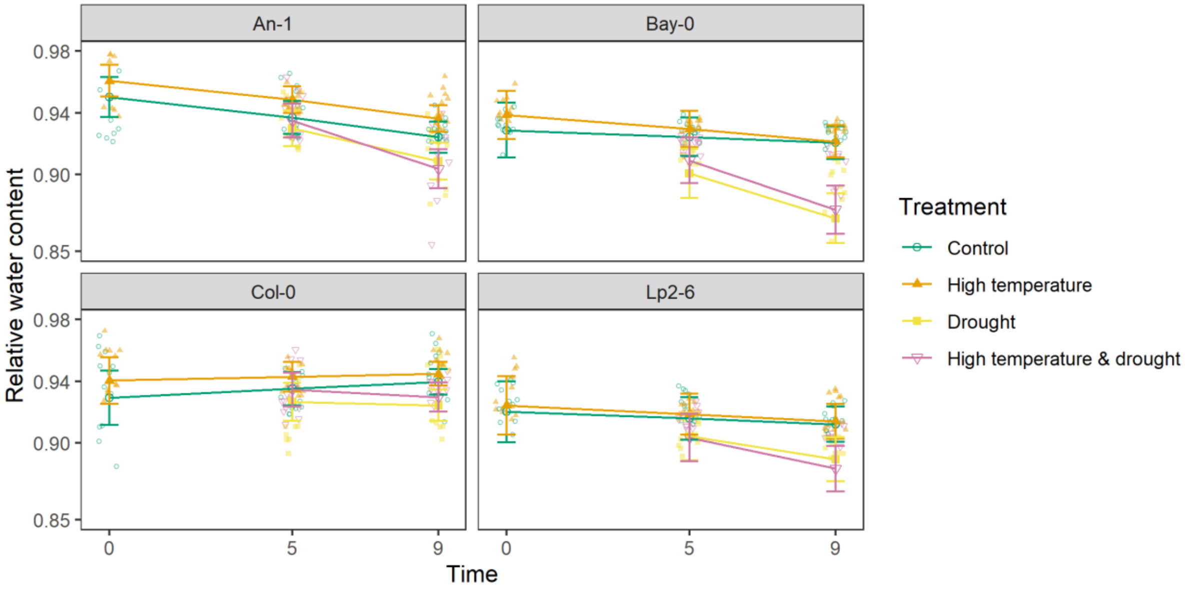


**Figure S14:** Measured relative water content in experiment I and estimated means for each combination of accession (panel), treatment (dots and lines) and time (days) using the fitted linear mixed models. Whiskers represent standard errors of the means, dots in the background indicate individual measurements. Time = 0 corresponds to the 10 true leaf stage where treatments were initiated.

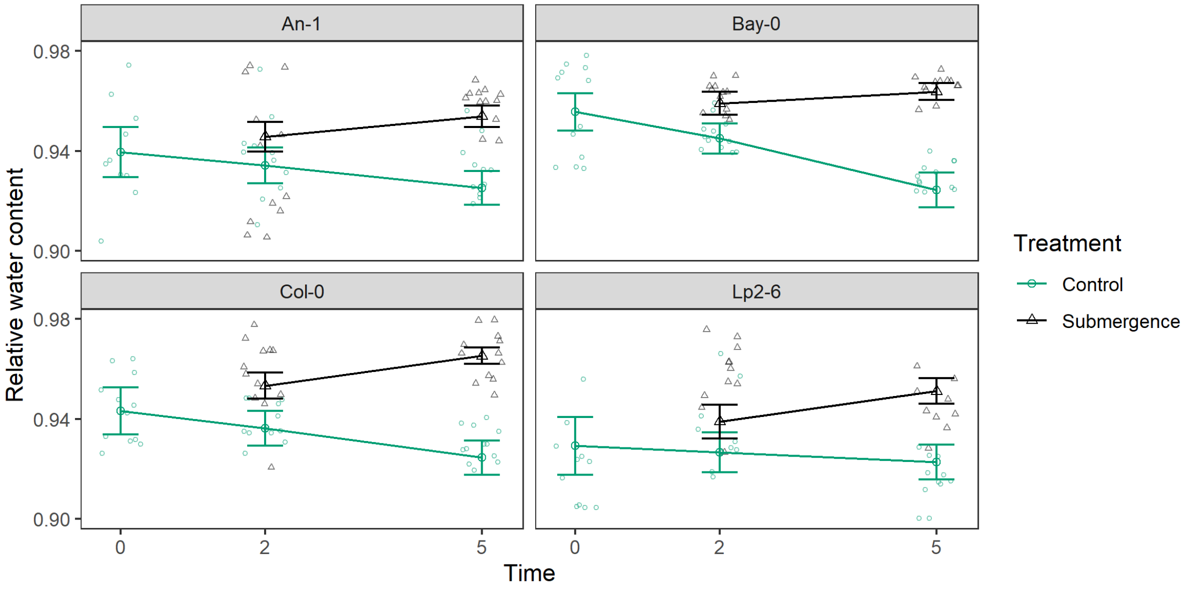


**Figure S15:** Measured relative water content in experiment II (phase a) and estimated means for each combination of accession (panel), treatment (dots and lines) and time (days) using the fitted linear mixed models. Whiskers represent standard errors of the means, dots in the background indicate individual measurements. Time = 0 corresponds to the 10 true leaf stage where treatments were initiated.

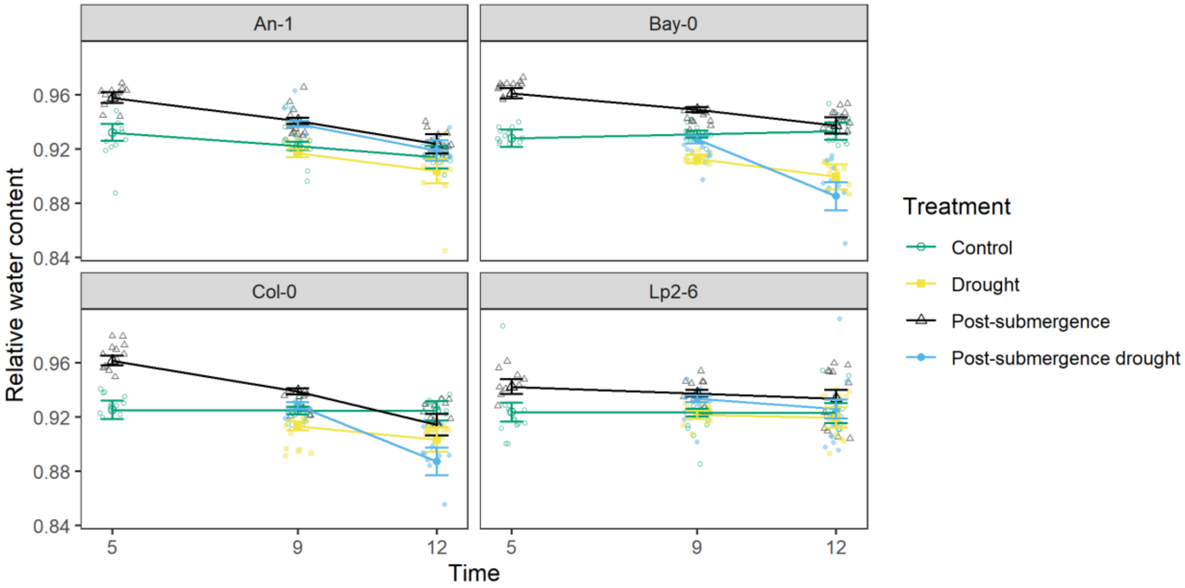


**Figure S16**: Measured relative water content in experiment II (phase b) and estimated means for each combination of accession (panel), treatment (dots and lines) and time (days) using the fitted linear mixed models. Whiskers represent standard errors of the means, dots in the background indicate individual measurements. Time = 5 corresponds to 5 days after the 10 true leaf stage was reached.

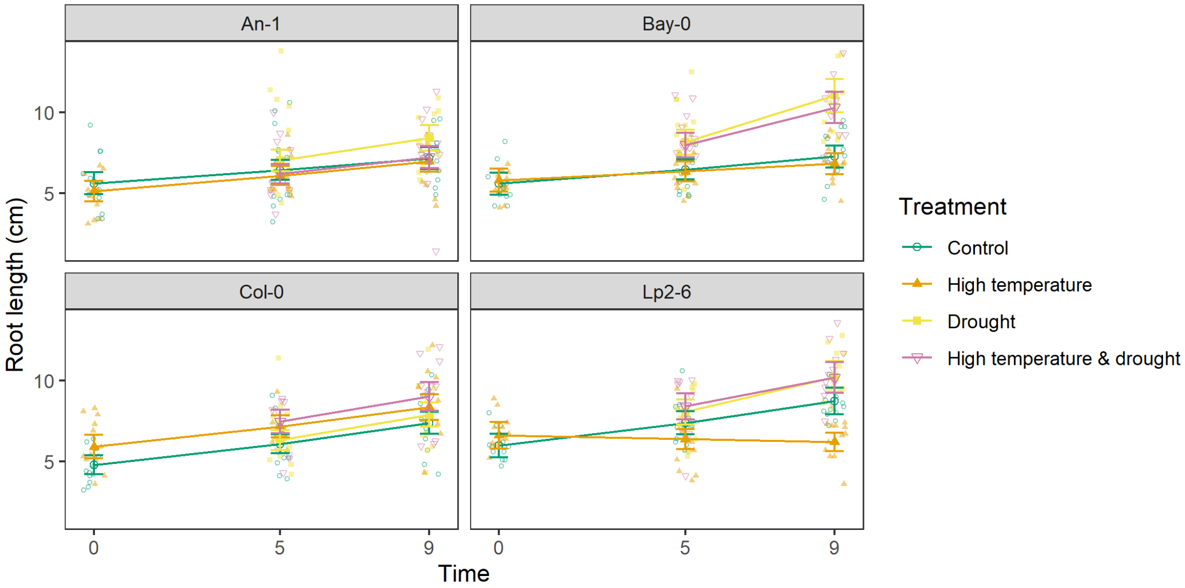


**Figure S17:** Measured root length in experiment I and estimated means for each combination of accession (panel), treatment (dots and lines) and time (days) using the fitted linear mixed models. Whiskers represent standard errors of the means, dots in the background indicate individual measurements. Time = 0 corresponds to the 10 true leaf stage where treatments were initiated.

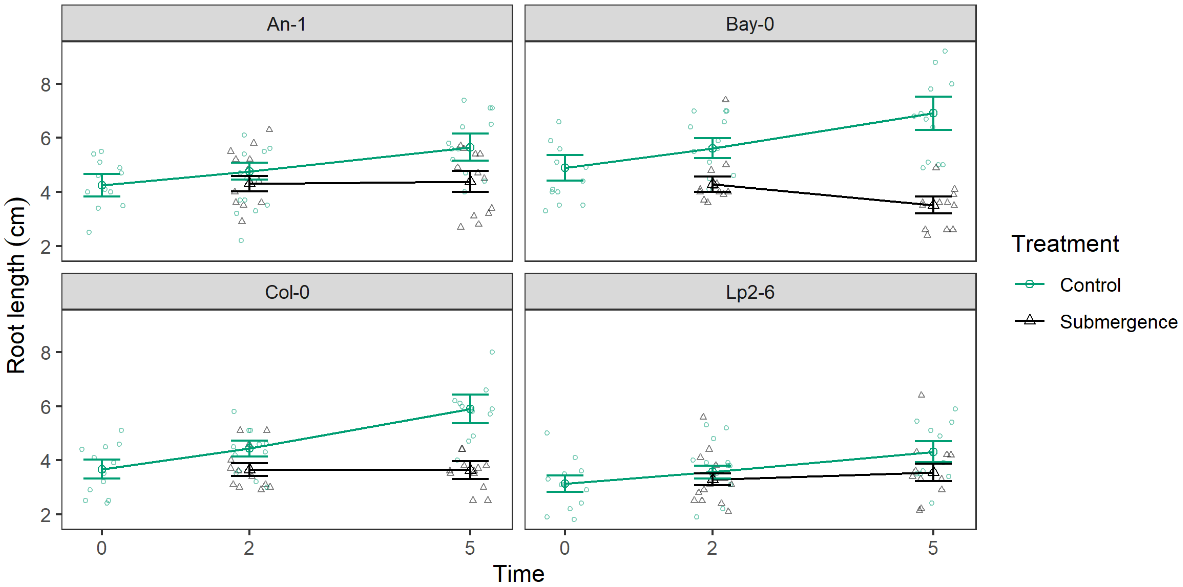


**Figure S18:** Measured root length in experiment II (phase a) and estimated means for each combination of accession (panel), treatment (dots and lines) and time (days) using the fitted linear mixed models. Whiskers represent standard errors of the means, dots in the background indicate individual measurements. Time = 0 corresponds to the 10 true leaf stage where treatments were initiated.

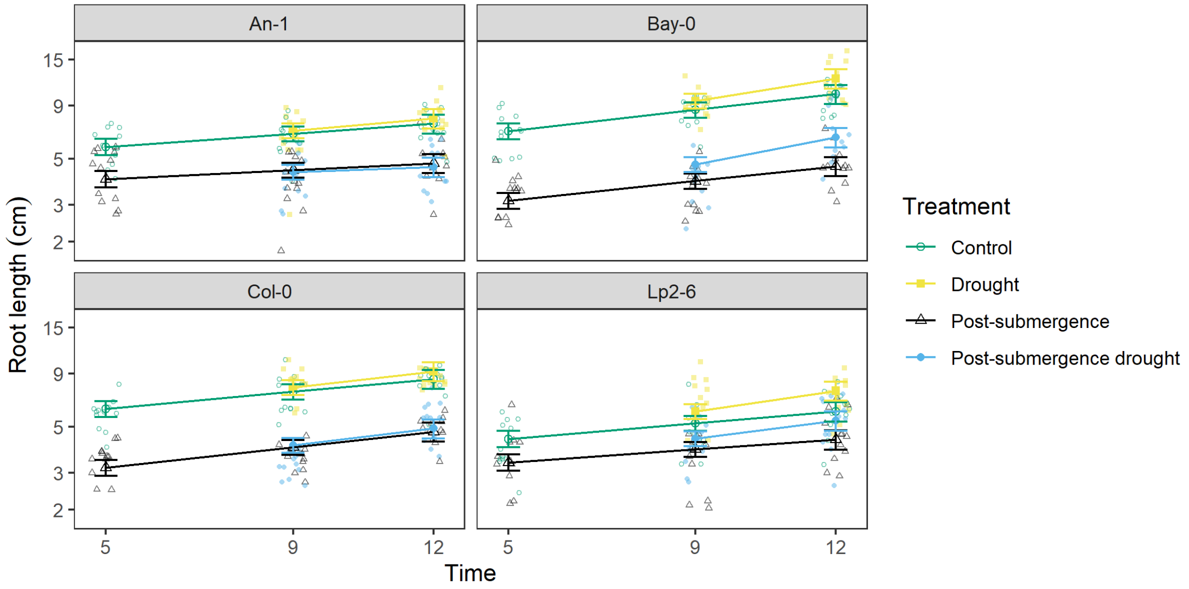


**Figure S19:** Measured root length in experiment II (phase b) and estimated means for each combination of accession (panel), treatment (dots and lines) and time (days) using the fitted linear mixed models. Whiskers represent standard errors of the means, dots in the background indicate individual measurements. Time = 5 corresponds to 5 days after the 10 true leaf stage was reached.

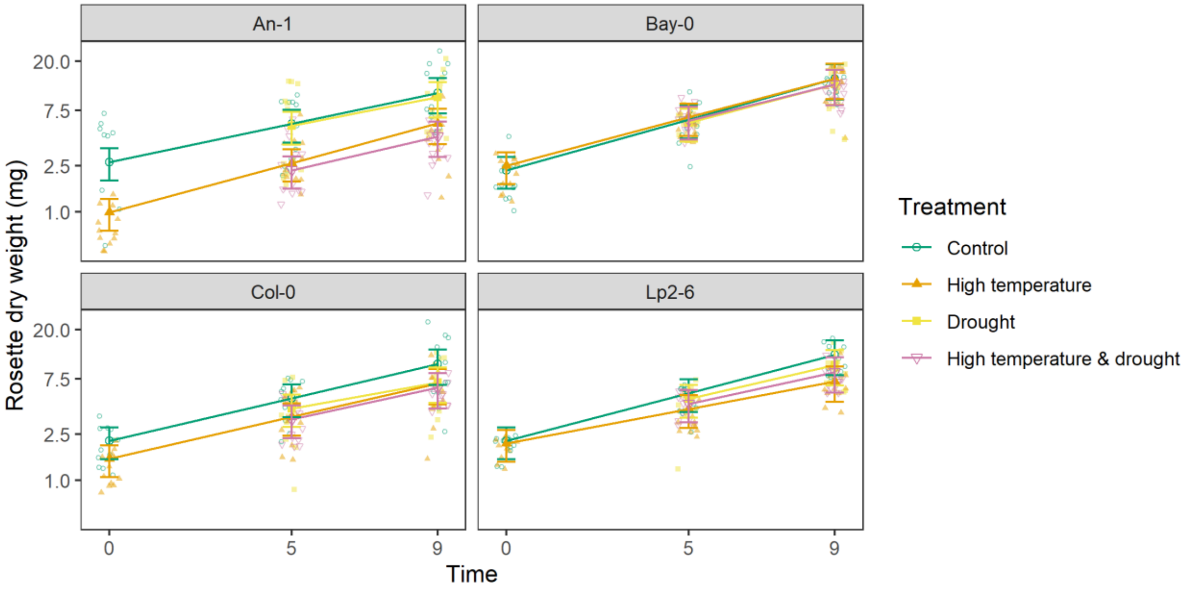


**Figure S20:** Measured rosette dry weight in experiment I and estimated means for each combination of accession (panel), treatment (dots and lines) and time (days) using the fitted linear mixed models. Whiskers represent standard errors of the means, dots in the background indicate individual measurements. Time = 0 corresponds to the 10 true leaf stage where treatments were initiated.

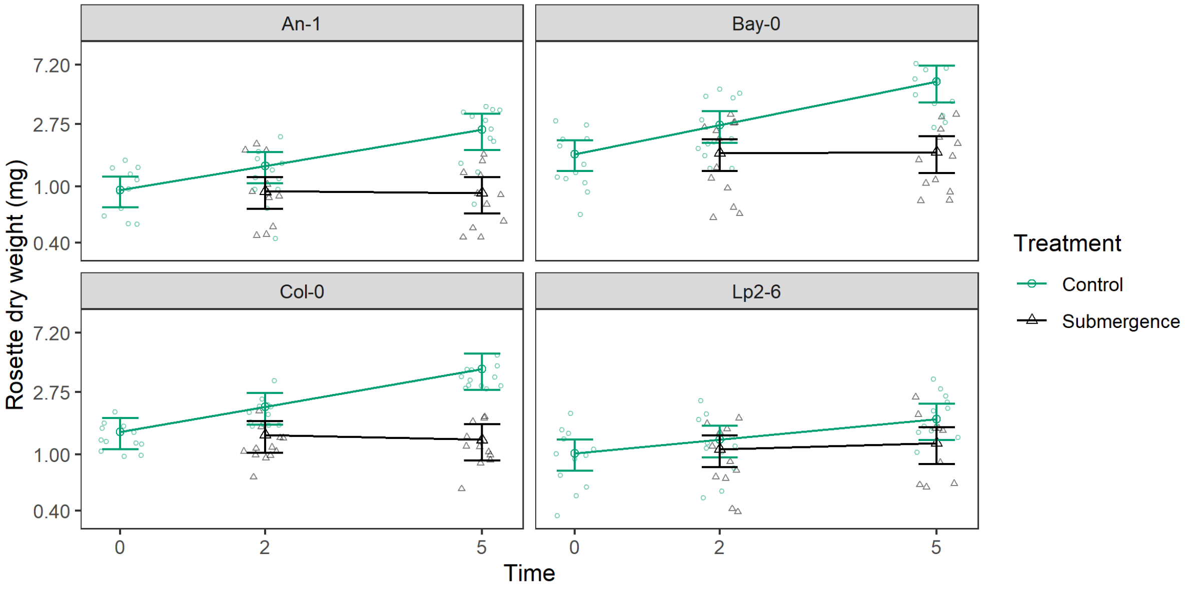


**Figure S21:** Measured rosette dry weight in experiment II (phase a) and estimated means for each combination of accession (panel), treatment (dots and lines) and time (days) using the fitted linear mixed models. Whiskers represent standard errors of the means, dots in the background indicate individual measurements. Time = 0 corresponds to the 10 true leaf stage where treatments were initiated.

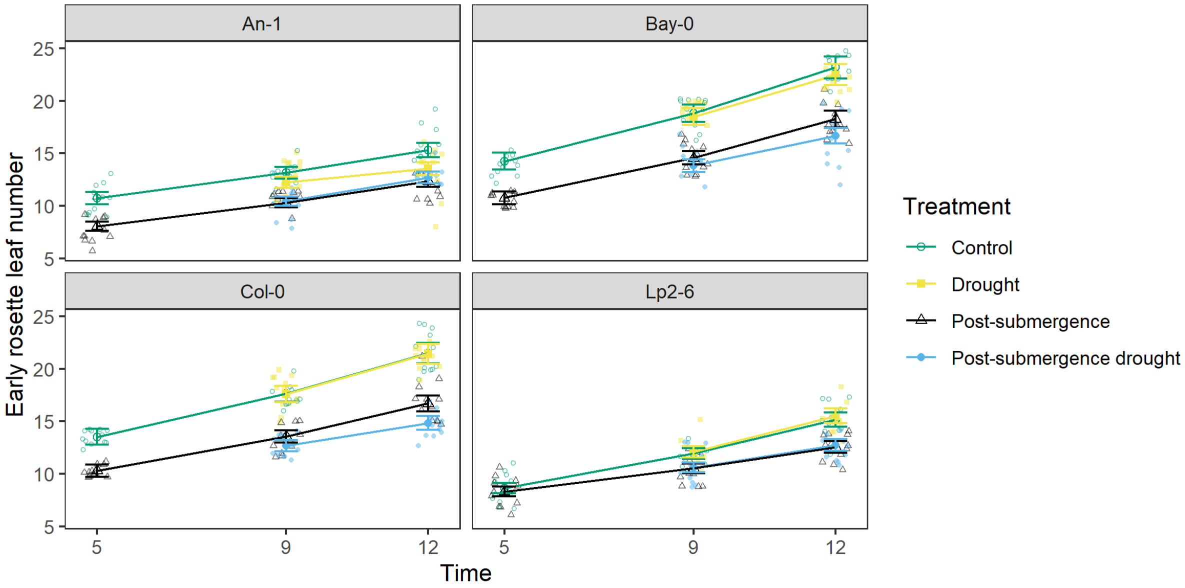


**Figure S22:** Measured rosette dry weight in experiment II (phase b) and estimated means for each combination of accession (panel), treatment (dots and lines) and time (days, X-axis) using the fitted linear mixed models. Whiskers represent standard errors of the means, dots in the background indicate individual measurements. Time = 5 corresponds to 5 days after the 10 true leaf stage was reached.

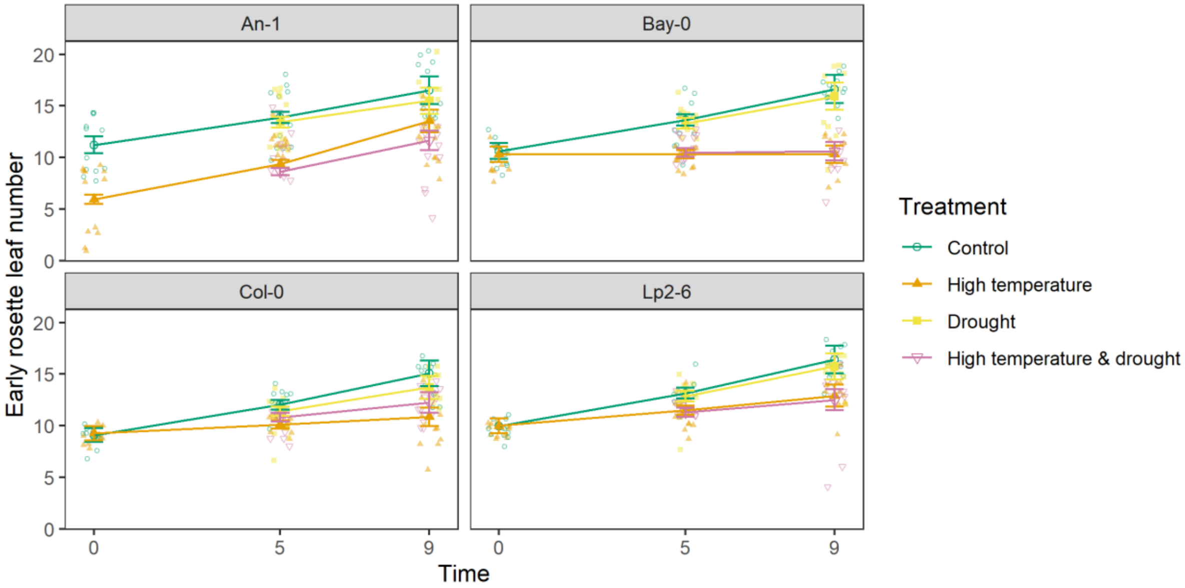


**Figure S23:** Measured early rosette leaf number in experiment I and estimated means for each combination of accession (panel), treatment (dots and lines) and time (days) using the fitted +linear mixed models. Whiskers represent standard errors of the means, dots in the background indicate individual measurements. Time = 0 corresponds to the 10 true leaf stage where treatments were initiated.

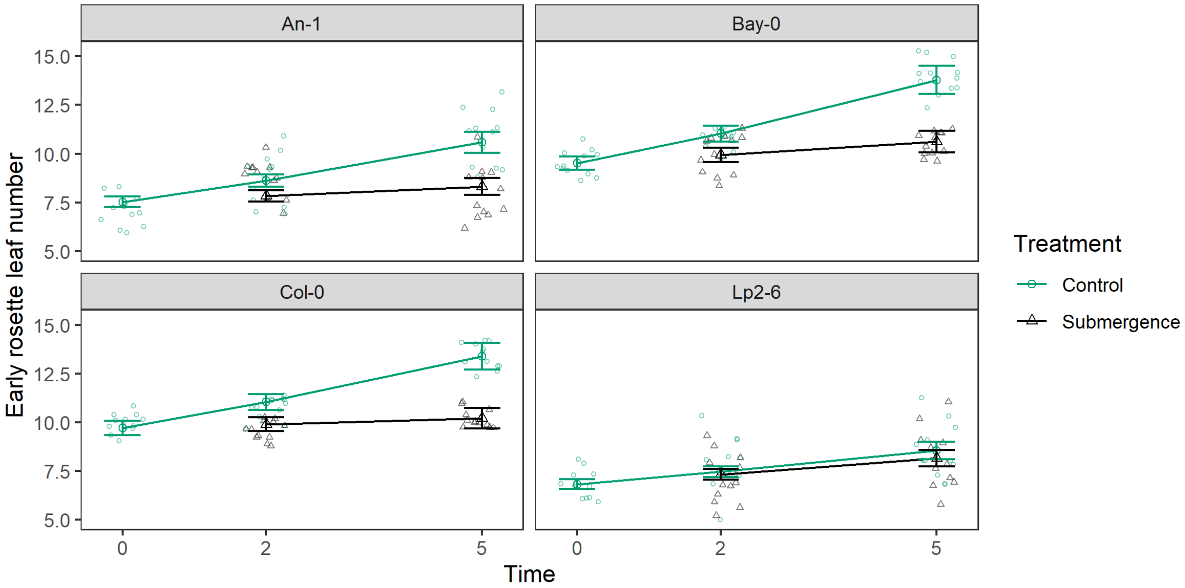


**Figure S24:** Measured early rosette leaf number in experiment II (phase a) and estimated means for each combination of accession (panel), treatment (dots and lines) and time (days) using the fitted linear mixed models. Whiskers represent standard errors of the means, dots in the background indicate individual measurements. Time = 0 corresponds to the 10 true leaf stage where treatments were initiated.

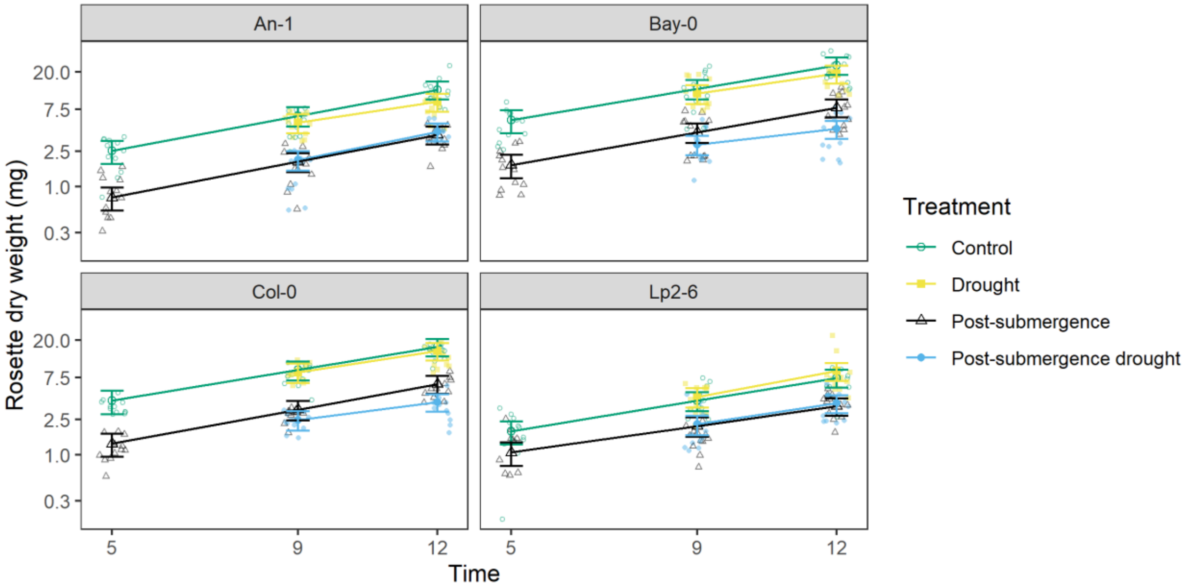


**Figure S25:** Measured early rosette leaf number in experiment II (phase b) and estimated means for each combination of accession (panel), treatment (dots and lines) and time (days) using the fitted linear mixed models. Whiskers represent standard errors of the means, dots in the background indicate individual measurements. Time = 5 corresponds to 5 days after the 10 true leaf stage was reached.

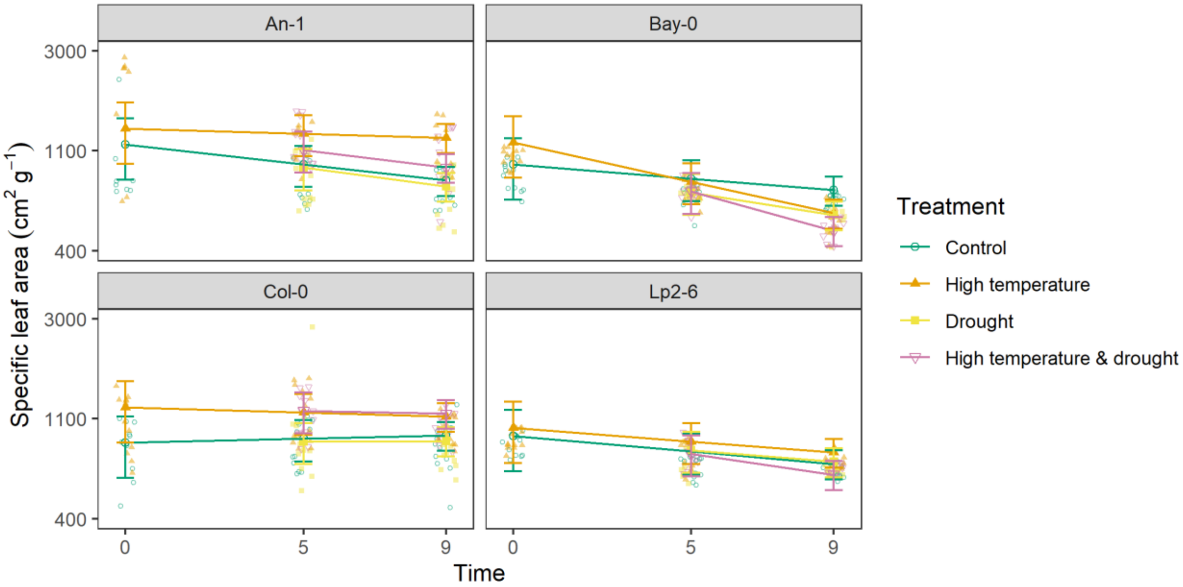


**Figure S26:** Measured specific leaf area in experiment I and estimated means for each combination of accession (panel), treatment (dots and lines) and time (days) using the fitted linear mixed models. Whiskers represent standard errors of the means, dots in the background indicate individual measurements. Time = 0 corresponds to the 10 true leaf stage where treatments were initiated.

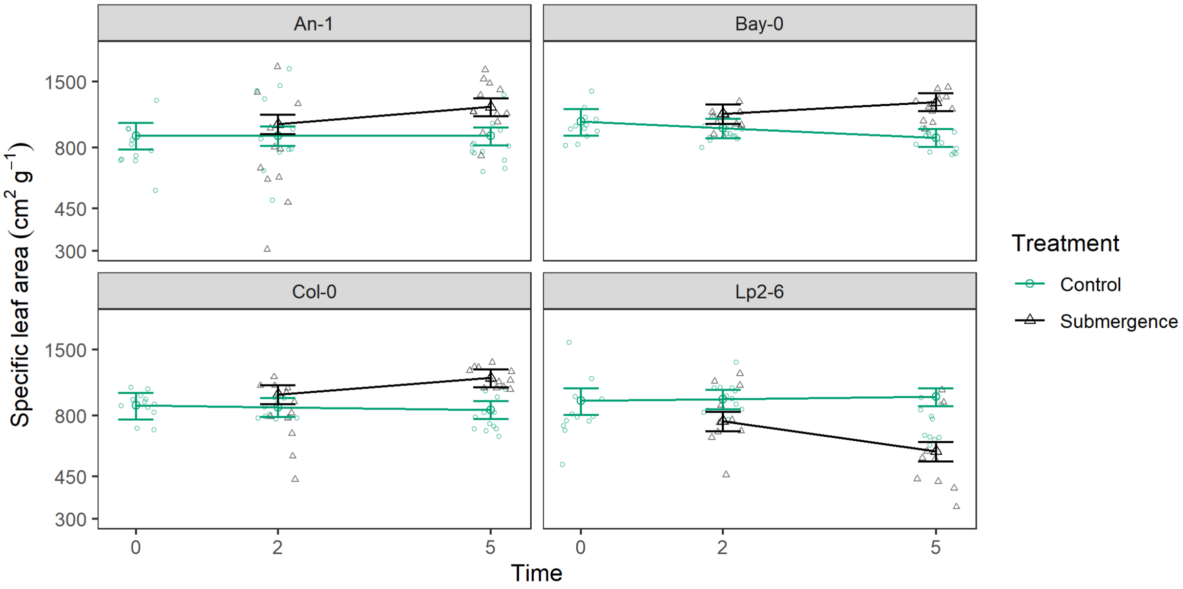


**Figure S27:** Measured specific leaf area in experiment II (phase a) and estimated means for each combination of accession (panel), treatment (dots and lines) and time (days) using the fitted linear mixed models. Whiskers represent standard errors of the means, dots in the background indicate individual measurements. Time = 0 corresponds to the 10 true leaf stage where treatments were initiated.

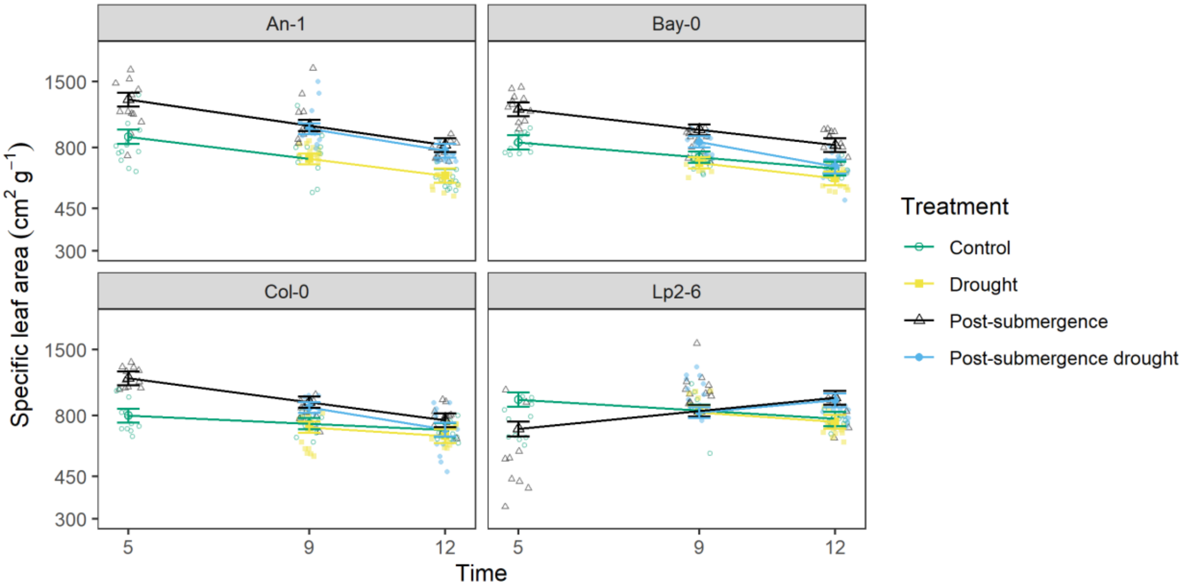


**Figure S28:** Measured specific leaf area in experiment II (phase b) and estimated means for each combination of accession (panel), treatment (dots and lines) and time (days) using the fitted linear mixed models. Whiskers represent standard errors of the means, dots in the background indicate individual measurements. Time = 5 corresponds to 5 days after the 10 true leaf stage was reached.
