## Supplementary material for "Effects of sub-lethal single, simultaneous, and sequential abiotic stresses on phenotypic traits of Arabidopsis thaliana": Data and R code: Metadata.md

# Experiment I_II

Each row corresponds to measurements on an Arabidopsis plant

**Genotype** = Arabidopsis accession being measured ("An-1", "Bay-0", "Col-0" or "Lp2-6")
**XP** = Identifier of this experiment (either "I" or "II")
**HighTemperature** = Growth temperature treatment (0 = control, 1 = high temperature)
**Drought** = Drought treatment (0 = No drought applied, 1 = Drought applied).
**Submergence** = Submergence treatment (0 = No submergence, 1 = Plants were submerged)
**DAS** = Days after sowing when the measurement was taken
**Timepoint** = Timepoint within the experimental protocol (from 1 to 5)
**Repetition** = Experimental repetition with experiment I (from 1 to 13)
**DW** = Rosette oven dry weight (mg)
**RootLength** = Maximum depth reached by the root system (cm)
**HypocotylLength** = Length of the hypocotyl (cm)
**Angle** = Angle of petiole insertion with respect to horizontal plane (°)
**RWC** = Relative water content of the rosette
**LeafNumber** = Total number of true leaves of the rosette
**PlantArea** = Total one-sided leaf (blade + petiole) area of the rosette (cm2)
**PetioleRatio** = Average ratio between petiole and leaf (petiole + blade) length
**ShapeBlade** = Average ratio between blade width and length assuming blades are shaped as ellipses
**SLA** = Specific leaf area including petioles (cm2/g)

# Experiment III

Each row corresponds to measurements on an Arabidopsis plant

**Genotype** = Arabidopsis accession being measured ("An-1", "Bay-0", "Col-0" or "Lp2-6")
**HighTemperature** = Growth temperature treatment (0 = control, 1 = high temperature)
**Drought** = Drought treatment (0 = No drought applied, 1 = Drought applied).
**Submergence** = Submergence treatment (0 = No submergence, 1 = Plants were submerged)
**Repetition** = Experimental repetition with experiment I (from 1 to 3)
**FT** = Flowering time in days after sowing
**LN** = Total number of rosette leaves produced
**Yield** = Seed yield of the plant (mg)
