## Supplementary material for "Effects of sub-lethal single, simultaneous, and sequential abiotic stresses on phenotypic traits of Arabidopsis thaliana": Data and R code: script.R

```
# INSTRUCTIONS
# In order to execute the script, please create a folder named Data with the contents of the zip file available as supplemental material to the article
# Also, crease folder named Results where the figures and tables will be generated
# The first time that the script runs it will install the necessary dependency, so a connection to the internet is required

# Load data and packages --------------------------------------------------

if(!require(easypackages)) install.packages("easypackages")

easypackages::packages("tidyverse", "nlme", "emmeans", "patchwork", 
                       "colorBlindness", prompt = FALSE)

# Some helper vectors
tnames = c("000" = "C", "001" = "D", "010" = "S", "011" = "PSD", "100" = "HT", 
           "101" = "HTD")
tnamesI = c("00" = "C", "01" = "D", "10" = "HT", "11" = "HTD")
tnamesII = c("00" = "C", "01" = "D", "10" = "S", "11" = "PSD")

labelsI = c(C = "Control", D = "Drought", HT = "High temperature", 
            HTD = "High temperature & drought")
colorsI = c(C = BLUISH_GREEN, HT = ORANGE, D = YELLOW, HTD = REDDISH_PURPLE)
shapesI = c(C = 1, HT = 17, D = 15, HTD = 6)

labelsIIa = c(C = "Control", S = "Submergence")
colorsIIa = c(C = BLUISH_GREEN, S = BLACK)
shapesIIa = c(C = 1, S = 2)

labelsIIb = c(C = "Control", D = "Drought", S = "Post-submergence", 
              PSD = "Post-submergence drought")
colorsIIb = c(C = BLUISH_GREEN, D = YELLOW, S = BLACK, PSD = SKY_BLUE)
shapesIIb = c(C = 1, D = 15, S = 2,  PSD = 16)


logit = function(x) log(x/(1 - x))


# Load the data
DryingCurve = read_csv("Data/DryingCurve.csv")
XP_I_II     = read_csv("Data/I_II.csv")
XP_III      = read_csv("Data/III.csv") %>% 
  mutate(Treatment = tnames[paste0(HighTemperature, Submergence, Drought)])


# Experiment I - Blade Shape ----------------------------------------------

# Filter the data
data_I_BS = XP_I_II %>% 
  filter(XP == 'I', !is.na(BladeShape)) %>% 
  mutate(Treatment = tnames[paste0(HighTemperature, Submergence, Drought)],
         Time = case_when(
           XP == "I" & Timepoint == 1 ~ 0,
           XP == "I" & Timepoint == 2 ~ 5,
           XP == "I" & Timepoint == 3 ~ 9))

# Fit linear mixed model
model_I_BS = lme(log(BladeShape)~ Genotype*HighTemperature*(Time + Drought:Time),
            random = ~ Time | Repetition, data = data_I_BS,
            control = lmeControl(maxIter = 1e4, msMaxIter = 1e4,
                                 opt = "optim"))

# F test on the fitted model
anova_I_BS = anova(model_I_BS, type = "marginal")
anova_I_BS = cbind(Term = row.names(anova_I_BS), anova_I_BS)
row.names(anova_I_BS) = NULL

# Marginal means
sigma = sqrt(sum(as.numeric(VarCorr(model_I_BS)[c(1,3),1])))
means_I_BS = bind_rows(
  # For timepoint 9
  ref_grid(model_I_BS, trans = "log", at = list(Time = 9)) %>% 
    summary(infer = c(F, F),  type = "response", 
            bias.adjust = TRUE, sigma = sigma),
  # For timepoint 5
  ref_grid(model_I_BS, trans = "log", at = list(Time = 5)) %>% 
    summary(infer = c(F, F),  type = "response", 
            bias.adjust = TRUE, sigma = sigma),
  # For timepoint 0
  ref_grid(model_I_BS, trans = "log", at = list(Time = 0)) %>% 
    summary(infer = c(F, F),  type = "response", 
            bias.adjust = TRUE, sigma = sigma)) %>% 
  mutate(Treatment = tnamesI[paste0(HighTemperature, Drought)]) %>% 
  filter(!(Time == 0 & Drought == 1))

# Figure S2
(ggplot(data = means_I_BS, aes(x = Time, y = response, colour = Treatment,
                               shape = Treatment)) +
  geom_point(data = data_I_BS, mapping = aes(y = BladeShape), alpha = 0.5,
               size = 0.8, position = position_jitter(width = 0.3),
             show.legend = FALSE) + 
  geom_point() + geom_line() +
  geom_errorbar(aes(ymax = response + SE, ymin = response - SE),
                width = 0.5) +
  facet_wrap(~Genotype) + 
  scale_y_continuous(limits = c(0.55,0.9)) + 
  scale_x_continuous(breaks = c(0,5,9), labels = c("0","5","9")) +
  scale_colour_manual(labels = labelsI, values = colorsI) +
  scale_shape_manual( labels = labelsI, values = shapesI) +
  labs(y = "Blade shape") +
  theme_bw() +
  theme(panel.grid = element_blank())) %>% 
  ggsave(filename = "Results/Figure_S2.png", plot = ., dpi = 300, 
         width = 16, height = 8, units = "cm",scale = 1.25)

# Experiment I - Early rosette leaf number --------------------------------

# Filter the data
data_I_LN = XP_I_II %>% 
  filter(XP == 'I', !is.na(LeafNumber)) %>% 
  mutate(Treatment = tnames[paste0(HighTemperature, Submergence, Drought)],
         Time = case_when(
           XP == "I" & Timepoint == 1 ~ 0,
           XP == "I" & Timepoint == 2 ~ 5,
           XP == "I" & Timepoint == 3 ~ 9))

# Fit linear mixed model
model_I_LN = lme(log(LeafNumber)~ Genotype*HighTemperature*(Time + Drought:Time),
                 random = ~ Time | Repetition, data = data_I_LN,
                 control = lmeControl(maxIter = 1e4, msMaxIter = 1e4,
                                      opt = "optim"))

# F test on the fitted model
anova_I_LN = anova(model_I_LN, type = "marginal")
anova_I_LN = cbind(Term = row.names(anova_I_LN), anova_I_LN)
row.names(anova_I_LN) = NULL

# Marginal means
sigma = sqrt(sum(as.numeric(VarCorr(model_I_LN)[c(1,3),1])))
means_I_LN = bind_rows(
  # For timepoint 9
  ref_grid(model_I_LN, trans = "log", at = list(Time = 9)) %>% 
    summary(infer = c(F, F),  type = "response", 
            bias.adjust = TRUE, sigma = sigma),
  # For timepoint 5
  ref_grid(model_I_LN, trans = "log", at = list(Time = 5)) %>% 
    summary(infer = c(F, F),  type = "response", 
            bias.adjust = TRUE, sigma = sigma),
  # For timepoint 0
  ref_grid(model_I_LN, trans = "log", at = list(Time = 0)) %>% 
    summary(infer = c(F, F),  type = "response", 
            bias.adjust = TRUE, sigma = sigma)) %>% 
  mutate(Treatment = tnamesI[paste0(HighTemperature, Drought)]) %>% 
  filter(!(Time == 0 & Drought == 1))

# Figure S23
(ggplot(data = means_I_LN, aes(x = Time, y = response, colour = Treatment,
                               shape = Treatment)) +
    geom_point(data = data_I_LN, mapping = aes(y = LeafNumber), alpha = 0.5,
               size = 0.8, position = position_jitter(width = 0.3),
               show.legend = FALSE) + 
    geom_point() + geom_line() +
    geom_errorbar(aes(ymax = response + SE, ymin = response - SE),
                  width = 0.5) +
    facet_wrap(~Genotype) + 
    scale_x_continuous(breaks = c(0,5,9), labels = c("0","5","9")) +
    scale_colour_manual(labels = labelsI, values = colorsI) +
    scale_shape_manual( labels = labelsI, values = shapesI) +
    labs(y = "Early rosette leaf number") +
    theme_bw() +
    theme(panel.grid = element_blank())) %>% 
  ggsave(filename = "Results/Figure_S23.png", plot = ., dpi = 300, 
         width = 16, height = 8, units = "cm",scale = 1.25)

# Experiment I - Leaf angle -----------------------------------------------

# Filter the data
data_I_Angle = XP_I_II %>% 
  filter(XP == 'I', !is.na(Angle), Angle != 0) %>% 
  mutate(Treatment = tnames[paste0(HighTemperature, Submergence, Drought)],
         Time = case_when(
           XP == "I" & Timepoint == 1 ~ 0,
           XP == "I" & Timepoint == 2 ~ 5,
           XP == "I" & Timepoint == 3 ~ 9),
         Angle = Angle/90)

# Fit linear mixed model
model_I_Angle = lme(logit(Angle)~ Genotype*HighTemperature*(Time + Drought:Time),
                 random = ~ Time | Repetition, data = data_I_Angle,
                 control = lmeControl(maxIter = 1e4, msMaxIter = 1e4,
                                      opt = "optim"))

# F test on the fitted model
anova_I_Angle = anova(model_I_Angle, type = "marginal")
anova_I_Angle = cbind(Term = row.names(anova_I_Angle), anova_I_Angle)
row.names(anova_I_Angle) = NULL

# Marginal means
sigma = sqrt(sum(as.numeric(VarCorr(model_I_Angle)[c(1,3),1])))
means_I_Angle = bind_rows(
  # For timepoint 9
  ref_grid(model_I_Angle, trans = "logit", at = list(Time = 9)) %>% 
    summary(infer = c(F, F),  type = "response", 
            bias.adjust = TRUE, sigma = sigma),
  # For timepoint 5
  ref_grid(model_I_Angle, trans = "logit", at = list(Time = 5)) %>% 
    summary(infer = c(F, F),  type = "response", 
            bias.adjust = TRUE, sigma = sigma),
  # For timepoint 0
  ref_grid(model_I_Angle, trans = "logit", at = list(Time = 0)) %>% 
    summary(infer = c(F, F),  type = "response", 
            bias.adjust = TRUE, sigma = sigma)) %>% 
  mutate(Treatment = tnamesI[paste0(HighTemperature, Drought)]) %>% 
  filter(!(Time == 0 & Drought == 1))

# Figure S5
(ggplot(data = means_I_Angle, aes(x = Time, y = response*90, colour = Treatment,
                               shape = Treatment)) +
    geom_point(data = data_I_Angle, mapping = aes(y = Angle*90), alpha = 0.5,
               size = 0.8, position = position_jitter(width = 0.3),
               show.legend = FALSE) + 
    geom_point() + geom_line() +
    geom_errorbar(aes(ymax = (response + SE)*90, ymin = (response - SE)*90),
                  width = 0.5) +
    facet_wrap(~Genotype) + 
    scale_x_continuous(breaks = c(0,5,9), labels = c("0","5","9")) +
    scale_colour_manual(labels = labelsI, values = colorsI) +
    scale_shape_manual( labels = labelsI, values = shapesI) +
    labs(y = quote(Leaf~angle~(degree))) +
    theme_bw() +
    theme(panel.grid = element_blank())) %>% 
  ggsave(filename = "Results/Figure_S5.png", plot = ., dpi = 300, 
         width = 16, height = 8, units = "cm",scale = 1.25)

# Experiment I - Leaf size ----------------------------------------

# Filter the data
data_I_LS = XP_I_II %>% 
  mutate(LS = PlantArea/LeafNumber) %>% 
  filter(XP == 'I', !is.na(LS)) %>% 
  mutate(Treatment = tnames[paste0(HighTemperature, Submergence, Drought)],
         Time = case_when(
           XP == "I" & Timepoint == 1 ~ 0,
           XP == "I" & Timepoint == 2 ~ 5,
           XP == "I" & Timepoint == 3 ~ 9))

# Fit linear mixed model
model_I_LS = lme(log(LS)~ Genotype*HighTemperature*(Time + Drought:Time),
                  random = ~ Time | Repetition, data = data_I_LS,
                  control = lmeControl(maxIter = 1e4, msMaxIter = 1e4,
                                       opt = "optim"))

# F test on the fitted model
anova_I_LS = anova(model_I_LS, type = "marginal")
anova_I_LS = cbind(Term = row.names(anova_I_LS), anova_I_LS)
row.names(anova_I_LS) = NULL

# Marginal means
sigma = sqrt(sum(as.numeric(VarCorr(model_I_LS)[c(1,3),1])))
means_I_LS = bind_rows(
  # For timepoint 9
  ref_grid(model_I_LS, trans = "log", at = list(Time = 9)) %>% 
    summary(infer = c(F, F),  type = "response", 
            bias.adjust = TRUE, sigma = sigma),
  # For timepoint 5
  ref_grid(model_I_LS, trans = "log", at = list(Time = 5)) %>% 
    summary(infer = c(F, F),  type = "response", 
            bias.adjust = TRUE, sigma = sigma),
  # For timepoint 0
  ref_grid(model_I_LS, trans = "log", at = list(Time = 0)) %>% 
    summary(infer = c(F, F),  type = "response", 
            bias.adjust = TRUE, sigma = sigma)) %>% 
  mutate(Treatment = tnamesI[paste0(HighTemperature, Drought)]) %>% 
  filter(!(Time == 0 & Drought == 1))

# Figure S8
(ggplot(data = means_I_LS, aes(x = Time, y = response, colour = Treatment,
                                shape = Treatment)) +
    geom_point(data = data_I_LS, mapping = aes(y = LS), alpha = 0.5,
               size = 0.8, position = position_jitter(width = 0.3),
               show.legend = FALSE) + 
    geom_point() + geom_line() +
    geom_errorbar(aes(ymax = response + SE, ymin = response - SE),
                  width = 0.5) +
    facet_wrap(~Genotype) + 
    scale_x_continuous(breaks = c(0,5,9), labels = c("0","5","9")) +
    scale_colour_manual(labels = labelsI, values = colorsI) +
    scale_shape_manual( labels = labelsI, values = shapesI) +
    labs(y = quote(Leaf~size~(cm^2))) +
    theme_bw() +
    theme(panel.grid = element_blank())) %>% 
  ggsave(filename = "Results/Figure_S8.png", plot = ., dpi = 300, 
         width = 16, height = 8, units = "cm",scale = 1.25)

# Experiment I - Petiole ratio --------------------------------------------

# Filter the data
data_I_PR = XP_I_II %>%
  filter(XP == 'I', !is.na(PetioleRatio)) %>% 
  mutate(Treatment = tnames[paste0(HighTemperature, Submergence, Drought)],
         Time = case_when(
           XP == "I" & Timepoint == 1 ~ 0,
           XP == "I" & Timepoint == 2 ~ 5,
           XP == "I" & Timepoint == 3 ~ 9))

# Fit linear mixed model
model_I_PR = lme(logit(PetioleRatio)~ Genotype*HighTemperature*(Time + Drought:Time),
                  random = ~ Time | Repetition, data = data_I_PR,
                  control = lmeControl(maxIter = 1e4, msMaxIter = 1e4,
                                       opt = "optim"))

# F test on the fitted model
anova_I_PR = anova(model_I_PR, type = "marginal")
anova_I_PR = cbind(Term = row.names(anova_I_PR), anova_I_PR)
row.names(anova_I_PR) = NULL

# Marginal means
sigma = sqrt(sum(as.numeric(VarCorr(model_I_PR)[c(1,3),1])))
means_I_PR = bind_rows(
  # For timepoint 9
  ref_grid(model_I_PR, trans = "logit", at = list(Time = 9)) %>% 
    summary(infer = c(F, F),  type = "response", 
            bias.adjust = TRUE, sigma = sigma),
  # For timepoint 5
  ref_grid(model_I_PR, trans = "logit", at = list(Time = 5)) %>% 
    summary(infer = c(F, F),  type = "response", 
            bias.adjust = TRUE, sigma = sigma),
  # For timepoint 0
  ref_grid(model_I_PR, trans = "logit", at = list(Time = 0)) %>% 
    summary(infer = c(F, F),  type = "response", 
            bias.adjust = TRUE, sigma = sigma)) %>% 
  mutate(Treatment = tnamesI[paste0(HighTemperature, Drought)]) %>% 
  filter(!(Time == 0 & Drought == 1))

# Figure S11
(ggplot(data = means_I_PR, aes(x = Time, y = response, colour = Treatment,
                                shape = Treatment)) +
    geom_point(data = data_I_PR, mapping = aes(y = PetioleRatio), alpha = 0.5,
               size = 0.8, position = position_jitter(width = 0.3),
               show.legend = FALSE) + 
    geom_point() + geom_line() +
    geom_errorbar(aes(ymax = response + SE, ymin = response - SE),
                  width = 0.5) +
    facet_wrap(~Genotype) + 
    scale_x_continuous(breaks = c(0,5,9), labels = c("0","5","9")) +
    scale_colour_manual(labels = labelsI, values = colorsI) +
    scale_shape_manual( labels = labelsI, values = shapesI) +
    labs(y = "Petiole ratio") +
    theme_bw() +
    theme(panel.grid = element_blank())) %>% 
  ggsave(filename = "Results/Figure_S11.png", plot = ., dpi = 300, 
         width = 16, height = 8, units = "cm",scale = 1.25)

# Experiment I - Relative water content -----------------------------------

# Filter the data
data_I_RWC = XP_I_II %>%
  filter(XP == 'I', !is.na(RWC)) %>% 
  mutate(Treatment = tnames[paste0(HighTemperature, Submergence, Drought)],
         Time = case_when(
           XP == "I" & Timepoint == 1 ~ 0,
           XP == "I" & Timepoint == 2 ~ 5,
           XP == "I" & Timepoint == 3 ~ 9))

# Fit linear mixed model
model_I_RWC = lme(logit(RWC)~ Genotype*HighTemperature*(Time + Drought:Time),
                 random = ~ Time | Repetition, data = data_I_RWC,
                 control = lmeControl(maxIter = 1e4, msMaxIter = 1e4,
                                      opt = "optim"))

# F test on the fitted model
anova_I_RWC = anova(model_I_RWC, type = "marginal")
anova_I_RWC = cbind(Term = row.names(anova_I_RWC), anova_I_RWC)
row.names(anova_I_RWC) = NULL

# Marginal means
sigma = sqrt(sum(as.numeric(VarCorr(model_I_RWC)[c(1,3),1])))
means_I_RWC = bind_rows(
  # For timepoint 9
  ref_grid(model_I_RWC, trans = "logit", at = list(Time = 9)) %>% 
    summary(infer = c(F, F),  type = "response", 
            bias.adjust = TRUE, sigma = sigma),
  # For timepoint 5
  ref_grid(model_I_RWC, trans = "logit", at = list(Time = 5)) %>% 
    summary(infer = c(F, F),  type = "response", 
            bias.adjust = TRUE, sigma = sigma),
  # For timepoint 0
  ref_grid(model_I_RWC, trans = "logit", at = list(Time = 0)) %>% 
    summary(infer = c(F, F),  type = "response", 
            bias.adjust = TRUE, sigma = sigma)) %>% 
  mutate(Treatment = tnamesI[paste0(HighTemperature, Drought)]) %>% 
  filter(!(Time == 0 & Drought == 1))

# Figure S14
(ggplot(data = means_I_RWC, aes(x = Time, y = response, colour = Treatment,
                               shape = Treatment)) +
    geom_point(data = data_I_RWC, mapping = aes(y = RWC), alpha = 0.5,
               size = 0.8, position = position_jitter(width = 0.3),
               show.legend = FALSE) + 
    geom_point() + geom_line() +
    geom_errorbar(aes(ymax = response + SE, ymin = response - SE),
                  width = 0.5) +
    facet_wrap(~Genotype) + 
    scale_x_continuous(breaks = c(0,5,9), labels = c("0","5","9")) +
    scale_y_continuous(limits = c(0.85,0.98), breaks = c(0.85,0.90,0.94,0.98)) +
    scale_colour_manual(labels = labelsI, values = colorsI) +
    scale_shape_manual( labels = labelsI, values = shapesI) +
    labs(y = "Relative water content") +
    theme_bw() +
    theme(panel.grid = element_blank())) %>% 
  ggsave(filename = "Results/Figure_S14.png", plot = ., dpi = 300, 
         width = 16, height = 8, units = "cm",scale = 1.25)

# Experiment I - Root length ----------------------------------------------

# Filter the data
data_I_RL = XP_I_II %>%
  filter(XP == 'I', !is.na(RootLength)) %>% 
  mutate(Treatment = tnames[paste0(HighTemperature, Submergence, Drought)],
         Time = case_when(
           XP == "I" & Timepoint == 1 ~ 0,
           XP == "I" & Timepoint == 2 ~ 5,
           XP == "I" & Timepoint == 3 ~ 9))

# Fit linear mixed model
model_I_RL = lme(log(RootLength)~ Genotype*HighTemperature*(Time + Drought:Time),
                  random = ~ Time | Repetition, data = data_I_RL,
                  control = lmeControl(maxIter = 1e4, msMaxIter = 1e4,
                                       opt = "optim"))

# F test on the fitted model
anova_I_RL = anova(model_I_RL, type = "marginal")
anova_I_RL = cbind(Term = row.names(anova_I_RL), anova_I_RL)
row.names(anova_I_RL) = NULL

# Marginal means
sigma = sqrt(sum(as.numeric(VarCorr(model_I_RL)[c(1,3),1])))
means_I_RL = bind_rows(
  # For timepoint 9
  ref_grid(model_I_RL, trans = "log", at = list(Time = 9)) %>% 
    summary(infer = c(F, F),  type = "response", 
            bias.adjust = TRUE, sigma = sigma(model_I_RL)),
  # For timepoint 5
  ref_grid(model_I_RL, trans = "log", at = list(Time = 5)) %>% 
    summary(infer = c(F, F),  type = "response", 
            bias.adjust = TRUE, sigma = sigma(model_I_RL)),
  # For timepoint 0
  ref_grid(model_I_RL, trans = "log", at = list(Time = 0)) %>% 
    summary(infer = c(F, F),  type = "response", 
            bias.adjust = TRUE, sigma = sigma(model_I_RL))) %>% 
  mutate(Treatment = tnamesI[paste0(HighTemperature, Drought)]) %>% 
  filter(!(Time == 0 & Drought == 1))

# Figure S17
(ggplot(data = means_I_RL, aes(x = Time, y = response, colour = Treatment,
                                shape = Treatment)) +
    geom_point(data = data_I_RL, mapping = aes(y = RootLength), alpha = 0.5,
               size = 0.8, position = position_jitter(width = 0.3),
               show.legend = FALSE) + 
    geom_point() + geom_line() +
    geom_errorbar(aes(ymax = response + SE, ymin = response - SE),
                  width = 0.5) +
    facet_wrap(~Genotype) + 
    scale_x_continuous(breaks = c(0,5,9), labels = c("0","5","9")) +
    scale_colour_manual(labels = labelsI, values = colorsI) +
    scale_shape_manual( labels = labelsI, values = shapesI) +
    labs(y = "Root length (cm)") +
    theme_bw() +
    theme(panel.grid = element_blank())) %>% 
  ggsave(filename = "Results/Figure_S17.png", plot = ., dpi = 300, 
         width = 16, height = 8, units = "cm",scale = 1.25)

# Experiment I - Rosette dry weight ---------------------------------------

# Filter the data
data_I_DW = XP_I_II %>%
  filter(XP == 'I', !is.na(DW)) %>% 
  mutate(Treatment = tnames[paste0(HighTemperature, Submergence, Drought)],
         Time = case_when(
           XP == "I" & Timepoint == 1 ~ 0,
           XP == "I" & Timepoint == 2 ~ 5,
           XP == "I" & Timepoint == 3 ~ 9))

# Fit linear mixed model
model_I_DW = lme(log(DW)~ Genotype*HighTemperature*(Time + Drought:Time),
                 random = ~ Time | Repetition, data = data_I_DW,
                 control = lmeControl(maxIter = 1e4, msMaxIter = 1e4,
                                      opt = "optim"))

# F test on the fitted model
anova_I_DW = anova(model_I_DW, type = "marginal")
anova_I_DW = cbind(Term = row.names(anova_I_DW), anova_I_DW)
row.names(anova_I_DW) = NULL

# Marginal means
sigma = sqrt(sum(as.numeric(VarCorr(model_I_DW)[c(1,3),1])))
means_I_DW = bind_rows(
  # For timepoint 9
  ref_grid(model_I_DW, trans = "log", at = list(Time = 9)) %>% 
    summary(infer = c(F, F),  type = "response", 
            bias.adjust = TRUE, sigma = sigma),
  # For timepoint 5
  ref_grid(model_I_DW, trans = "log", at = list(Time = 5)) %>% 
    summary(infer = c(F, F),  type = "response", 
            bias.adjust = TRUE, sigma = sigma),
  # For timepoint 0
  ref_grid(model_I_DW, trans = "log", at = list(Time = 0)) %>% 
    summary(infer = c(F, F),  type = "response", 
            bias.adjust = TRUE, sigma = sigma)) %>% 
  mutate(Treatment = tnamesI[paste0(HighTemperature, Drought)]) %>% 
  filter(!(Time == 0 & Drought == 1))

# Figure S20
(ggplot(data = means_I_DW, aes(x = Time, y = response, colour = Treatment,
                               shape = Treatment)) +
    geom_point(data = data_I_DW, mapping = aes(y = DW), alpha = 0.5,
               size = 0.8, position = position_jitter(width = 0.3),
               show.legend = FALSE) + 
    geom_point() + geom_line() +
    geom_errorbar(aes(ymax = response + SE, ymin = response - SE),
                  width = 0.5) +
    facet_wrap(~Genotype) + 
    scale_x_continuous(breaks = c(0,5,9), labels = c("0","5","9")) +
    scale_y_continuous(trans = "log", breaks = c(1,2.5,7.5,20)) + 
    scale_colour_manual(labels = labelsI, values = colorsI) +
    scale_shape_manual( labels = labelsI, values = shapesI) +
    labs(y = "Rosette dry weight (mg)") +
    theme_bw() +
    theme(panel.grid = element_blank())) %>% 
  ggsave(filename = "Results/Figure_S20.png", plot = ., dpi = 300, 
         width = 16, height = 8, units = "cm",scale = 1.25)


# Experiment I - Specific leaf area ---------------------------------------

# Filter the data
data_I_SLA = XP_I_II %>%
  filter(XP == 'I', !is.na(SLA)) %>% 
  mutate(Treatment = tnames[paste0(HighTemperature, Submergence, Drought)],
         Time = case_when(
           XP == "I" & Timepoint == 1 ~ 0,
           XP == "I" & Timepoint == 2 ~ 5,
           XP == "I" & Timepoint == 3 ~ 9))

# Fit linear mixed model
model_I_SLA = lme(log(SLA)~ Genotype*HighTemperature*(Time + Drought:Time),
                 random = ~ Time | Repetition, data = data_I_SLA,
                 control = lmeControl(maxIter = 1e4, msMaxIter = 1e4,
                                      opt = "optim"))

# F test on the fitted model
anova_I_SLA = anova(model_I_SLA, type = "marginal")
anova_I_SLA = cbind(Term = row.names(anova_I_SLA), anova_I_SLA)
row.names(anova_I_SLA) = NULL

# Marginal means
sigma = sqrt(sum(as.numeric(VarCorr(model_I_SLA)[c(1,3),1])))
means_I_SLA = bind_rows(
  # For timepoint 9
  ref_grid(model_I_SLA, trans = "log", at = list(Time = 9)) %>% 
    summary(infer = c(F, F),  type = "response", 
            bias.adjust = TRUE, sigma = sigma),
  # For timepoint 5
  ref_grid(model_I_SLA, trans = "log", at = list(Time = 5)) %>% 
    summary(infer = c(F, F),  type = "response", 
            bias.adjust = TRUE, sigma = sigma),
  # For timepoint 0
  ref_grid(model_I_SLA, trans = "log", at = list(Time = 0)) %>% 
    summary(infer = c(F, F),  type = "response", 
            bias.adjust = TRUE, sigma = sigma)) %>% 
  mutate(Treatment = tnamesI[paste0(HighTemperature, Drought)]) %>% 
  filter(!(Time == 0 & Drought == 1))

# Figure S26
(ggplot(data = means_I_SLA, aes(x = Time, y = response, colour = Treatment,
                               shape = Treatment)) +
    geom_point(data = data_I_SLA, mapping = aes(y = SLA), alpha = 0.5,
               size = 0.8, position = position_jitter(width = 0.3),
               show.legend = FALSE) + 
    geom_point() + geom_line() +
    geom_errorbar(aes(ymax = response + SE, ymin = response - SE),
                  width = 0.5) +
    facet_wrap(~Genotype) + 
    scale_x_continuous(breaks = c(0,5,9), labels = c("0","5","9")) +
    scale_y_continuous(trans = "log", breaks = c(400,1100,3000), 
                       limits = c(400,3000)) + 
    scale_colour_manual(labels = labelsI, values = colorsI) +
    scale_shape_manual( labels = labelsI, values = shapesI) +
    labs(y = quote(Specific~leaf~area~(cm^2~g^{-1}))) +
    theme_bw() +
    theme(panel.grid = element_blank())) %>% 
  ggsave(filename = "Results/Figure_S26.png", plot = ., dpi = 300, 
         width = 16, height = 8, units = "cm",scale = 1.25)

# Experiment IIa - Blade Shape --------------------------------------------

# Filter the data
data_IIa_BS = XP_I_II %>% 
  filter(XP == 'II', Timepoint <= 3, !is.na(BladeShape)) %>% 
  mutate(Treatment = tnames[paste0(HighTemperature, Submergence, Drought)],
         Time = case_when(
           XP == "II" & Timepoint == 1 ~ 0,
           XP == "II" & Timepoint == 2 ~ 2,
           XP == "II" & Timepoint == 3 ~ 5))

# Fit linear mixed model
model_IIa_BS = lme(log(BladeShape)~Genotype*(Time + Submergence:Time),
                 random = ~ Time | Repetition, data = data_IIa_BS,
                 control = lmeControl(maxIter = 1e4, msMaxIter = 1e4,
                                      opt = "optim"))

# F test on the fitted model
anova_IIa_BS = anova(model_IIa_BS, type = "marginal")
anova_IIa_BS = cbind(Term = row.names(anova_IIa_BS), anova_IIa_BS)
row.names(anova_IIa_BS) = NULL

# Marginal means
sigma = sqrt(sum(as.numeric(VarCorr(model_IIa_BS)[c(1,3),1])))
means_IIa_BS = bind_rows(
  # For timepoint 9
  ref_grid(model_IIa_BS, trans = "log", at = list(Time = 5)) %>% 
    summary(infer = c(F, F),  type = "response", 
            bias.adjust = TRUE, sigma = sigma),
  # For timepoint 5
  ref_grid(model_IIa_BS, trans = "log", at = list(Time = 2)) %>% 
    summary(infer = c(F, F),  type = "response", 
            bias.adjust = TRUE, sigma = sigma),
  # For timepoint 0
  ref_grid(model_IIa_BS, trans = "log", at = list(Time = 0)) %>% 
    summary(infer = c(F, F),  type = "response", 
            bias.adjust = TRUE, sigma = sigma)) %>% 
  mutate(Treatment = ifelse(Submergence, 'S', "C")) %>% 
  filter(!(Time == 0 & Submergence == 1))

# Figure S3
(ggplot(data = means_IIa_BS, aes(x = Time, y = response, colour = Treatment,
                               shape = Treatment)) +
    geom_point(data = data_IIa_BS, mapping = aes(y = BladeShape), alpha = 0.5,
               size = 0.8, position = position_jitter(width = 0.3),
               show.legend = FALSE) + 
    geom_point() + geom_line() +
    geom_errorbar(aes(ymax = response + SE, ymin = response - SE),
                  width = 0.5) +
    facet_wrap(~Genotype) + 
    scale_x_continuous(breaks = c(0,2,5), labels = c("0","2","5")) +
    scale_colour_manual(labels = labelsIIa, values = colorsIIa) +
    scale_shape_manual(labels = labelsIIa, values = shapesIIa) +
    labs(y = "Blade shape") +
    theme_bw() +
    theme(panel.grid = element_blank())) %>% 
  ggsave(filename = "Results/Figure_S3.png", plot = ., dpi = 300, 
         width = 16, height = 8, units = "cm",scale = 1.25)


# Experiment IIa - Early rosette leaf number ------------------------------

# Filter the data
data_IIa_LN = XP_I_II %>% 
  filter(XP == 'II', Timepoint <= 3, !is.na(LeafNumber)) %>% 
  mutate(Treatment = tnames[paste0(HighTemperature, Submergence, Drought)],
         Time = case_when(
           XP == "II" & Timepoint == 1 ~ 0,
           XP == "II" & Timepoint == 2 ~ 2,
           XP == "II" & Timepoint == 3 ~ 5))

# Fit linear mixed model
model_IIa_LN = lme(log(LeafNumber)~Genotype*(Time + Submergence:Time),
                   random = ~ Time | Repetition, data = data_IIa_LN,
                   control = lmeControl(maxIter = 1e4, msMaxIter = 1e4,
                                        opt = "optim"))

# F test on the fitted model
anova_IIa_LN = anova(model_IIa_LN, type = "marginal")
anova_IIa_LN = cbind(Term = row.names(anova_IIa_LN), anova_IIa_LN)
row.names(anova_IIa_LN) = NULL

# Marginal means
sigma = sqrt(sum(as.numeric(VarCorr(model_IIa_LN)[c(1,3),1])))
means_IIa_LN = bind_rows(
  # For timepoint 9
  ref_grid(model_IIa_LN, trans = "log", at = list(Time = 5)) %>% 
    summary(infer = c(F, F),  type = "response", 
            bias.adjust = TRUE, sigma = sigma),
  # For timepoint 5
  ref_grid(model_IIa_LN, trans = "log", at = list(Time = 2)) %>% 
    summary(infer = c(F, F),  type = "response", 
            bias.adjust = TRUE, sigma = sigma),
  # For timepoint 0
  ref_grid(model_IIa_LN, trans = "log", at = list(Time = 0)) %>% 
    summary(infer = c(F, F),  type = "response", 
            bias.adjust = TRUE, sigma = sigma)) %>% 
  mutate(Treatment = ifelse(Submergence, 'S', "C")) %>% 
  filter(!(Time == 0 & Submergence == 1))

# Figure S24
(ggplot(data = means_IIa_LN, aes(x = Time, y = response, colour = Treatment,
                                 shape = Treatment)) +
    geom_point(data = data_IIa_LN, mapping = aes(y = LeafNumber), alpha = 0.5,
               size = 0.8, position = position_jitter(width = 0.3),
               show.legend = FALSE) + 
    geom_point() + geom_line() +
    geom_errorbar(aes(ymax = response + SE, ymin = response - SE),
                  width = 0.5) +
    facet_wrap(~Genotype) + 
    scale_x_continuous(breaks = c(0,2,5), labels = c("0","2","5")) +
    scale_colour_manual(labels = labelsIIa, values = colorsIIa) +
    scale_shape_manual(labels = labelsIIa, values = shapesIIa) +
    labs(y = "Early rosette leaf number") +
    theme_bw() +
    theme(panel.grid = element_blank())) %>% 
  ggsave(filename = "Results/Figure_S24.png", plot = ., dpi = 300, 
         width = 16, height = 8, units = "cm",scale = 1.25)

# Experiment IIa - Leaf angle ---------------------------------------------

# Filter the data
data_IIa_Angle = XP_I_II %>% 
  filter(XP == 'II', Timepoint <= 3, !is.na(Angle), Angle != 0) %>% 
  mutate(Treatment = tnames[paste0(HighTemperature, Submergence, Drought)],
         Time = case_when(
           XP == "II" & Timepoint == 1 ~ 0,
           XP == "II" & Timepoint == 2 ~ 2,
           XP == "II" & Timepoint == 3 ~ 5),
         Angle = Angle/90)

# Fit linear mixed model
model_IIa_Angle = lme(logit(Angle)~Genotype*(Time + Submergence:Time),
                   random = ~ Time | Repetition, data = data_IIa_Angle,
                   control = lmeControl(maxIter = 1e4, msMaxIter = 1e4,
                                        opt = "optim"))

# F test on the fitted model
anova_IIa_Angle = anova(model_IIa_Angle, type = "marginal")
anova_IIa_Angle = cbind(Term = row.names(anova_IIa_Angle), anova_IIa_Angle)
row.names(anova_IIa_Angle) = NULL

# Marginal means
sigma = sqrt(sum(as.numeric(VarCorr(model_IIa_Angle)[c(1,3),1])))
means_IIa_Angle = bind_rows(
  # For timepoint 9
  ref_grid(model_IIa_Angle, trans = "logit", at = list(Time = 5)) %>% 
    summary(infer = c(F, F),  type = "response", 
            bias.adjust = TRUE, sigma = sigma),
  # For timepoint 5
  ref_grid(model_IIa_Angle, trans = "logit", at = list(Time = 2)) %>% 
    summary(infer = c(F, F),  type = "response", 
            bias.adjust = TRUE, sigma = sigma),
  # For timepoint 0
  ref_grid(model_IIa_Angle, trans = "logit", at = list(Time = 0)) %>% 
    summary(infer = c(F, F),  type = "response", 
            bias.adjust = TRUE, sigma = sigma)) %>% 
  mutate(Treatment = ifelse(Submergence, 'S', "C")) %>% 
  filter(!(Time == 0 & Submergence == 1))

# Figure S6
(ggplot(data = means_IIa_Angle, aes(x = Time, y = response*90, colour = Treatment,
                                 shape = Treatment)) +
    geom_point(data = data_IIa_Angle, mapping = aes(y = Angle*90), alpha = 0.5,
               size = 0.8, position = position_jitter(width = 0.3),
               show.legend = FALSE) + 
    geom_point() + geom_line() +
    geom_errorbar(aes(ymax = (response + SE)*90, ymin = (response - SE)*90),
                  width = 0.5) +
    facet_wrap(~Genotype) + 
    scale_x_continuous(breaks = c(0,2,5), labels = c("0","2","5")) +
    scale_colour_manual(labels = labelsIIa, values = colorsIIa) +
    scale_shape_manual(labels = labelsIIa, values = shapesIIa) +
    labs(y = quote(Leaf~angle~(degree))) +
    theme_bw() +
    theme(panel.grid = element_blank())) %>% 
  ggsave(filename = "Results/Figure_S6.png", plot = ., dpi = 300, 
         width = 16, height = 8, units = "cm",scale = 1.25)

# Experiment IIa - Leaf size --------------------------------------


# Filter the data
data_IIa_LS = XP_I_II %>% 
  mutate(LS = PlantArea/LeafNumber) %>% 
  filter(XP == 'II', Timepoint <= 3, !is.na(LS)) %>% 
  mutate(Treatment = tnames[paste0(HighTemperature, Submergence, Drought)],
         Time = case_when(
           XP == "II" & Timepoint == 1 ~ 0,
           XP == "II" & Timepoint == 2 ~ 2,
           XP == "II" & Timepoint == 3 ~ 5))

# Fit linear mixed model
model_IIa_LS = lme(log(LS)~Genotype*(Time + Submergence:Time),
                   random = ~ Time | Repetition, data = data_IIa_LS,
                   control = lmeControl(maxIter = 1e4, msMaxIter = 1e4,
                                        opt = "optim"))

# F test on the fitted model
anova_IIa_LS = anova(model_IIa_LS, type = "marginal")
anova_IIa_LS = cbind(Term = row.names(anova_IIa_LS), anova_IIa_LS)
row.names(anova_IIa_LS) = NULL

# Marginal means
sigma = sqrt(sum(as.numeric(VarCorr(model_IIa_LS)[c(1,3),1])))
means_IIa_LS = bind_rows(
  # For timepoint 9
  ref_grid(model_IIa_LS, trans = "log", at = list(Time = 5)) %>% 
    summary(infer = c(F, F),  type = "response", 
            bias.adjust = TRUE, sigma = sigma),
  # For timepoint 5
  ref_grid(model_IIa_LS, trans = "log", at = list(Time = 2)) %>% 
    summary(infer = c(F, F),  type = "response", 
            bias.adjust = TRUE, sigma = sigma),
  # For timepoint 0
  ref_grid(model_IIa_LS, trans = "log", at = list(Time = 0)) %>% 
    summary(infer = c(F, F),  type = "response", 
            bias.adjust = TRUE, sigma = sigma)) %>% 
  mutate(Treatment = ifelse(Submergence, 'S', "C")) %>% 
  filter(!(Time == 0 & Submergence == 1))

# Figure S9
(ggplot(data = means_IIa_LS, aes(x = Time, y = response, colour = Treatment,
                                 shape = Treatment)) +
    geom_point(data = data_IIa_LS, mapping = aes(y = LS), alpha = 0.5,
               size = 0.8, position = position_jitter(width = 0.3),
               show.legend = FALSE) + 
    geom_point() + geom_line() +
    geom_errorbar(aes(ymax = response + SE, ymin = response - SE),
                  width = 0.5) +
    facet_wrap(~Genotype) + 
    scale_x_continuous(breaks = c(0,2,5), labels = c("0","2","5")) +
    scale_colour_manual(labels = labelsIIa, values = colorsIIa) +
    scale_shape_manual(labels = labelsIIa, values = shapesIIa) +
    labs(y = quote(Leaf~size~(cm^2))) +
    theme_bw() +
    theme(panel.grid = element_blank())) %>% 
  ggsave(filename = "Results/Figure_S9.png", plot = ., dpi = 300, 
         width = 16, height = 8, units = "cm",scale = 1.25)

# Experiment IIa - Petiole ratio ------------------------------------------


# Filter the data
data_IIa_PR = XP_I_II %>% 
  filter(XP == 'II', Timepoint <= 3, !is.na(PetioleRatio)) %>% 
  mutate(Treatment = tnames[paste0(HighTemperature, Submergence, Drought)],
         Time = case_when(
           XP == "II" & Timepoint == 1 ~ 0,
           XP == "II" & Timepoint == 2 ~ 2,
           XP == "II" & Timepoint == 3 ~ 5))

# Fit linear mixed model
model_IIa_PR = lme(logit(PetioleRatio)~Genotype*(Time + Submergence:Time),
                    random = ~ Time | Repetition, data = data_IIa_PR,
                    control = lmeControl(maxIter = 1e4, msMaxIter = 1e4,
                                         opt = "optim"))

# F test on the fitted model
anova_IIa_PR = anova(model_IIa_PR, type = "marginal")
anova_IIa_PR = cbind(Term = row.names(anova_IIa_PR), anova_IIa_PR)
row.names(anova_IIa_PR) = NULL

# Marginal means
sigma = sqrt(sum(as.numeric(VarCorr(model_IIa_PR)[c(1,3),1])))
means_IIa_PR = bind_rows(
  # For timepoint 9
  ref_grid(model_IIa_PR, trans = "logit", at = list(Time = 5)) %>% 
    summary(infer = c(F, F),  type = "response", 
            bias.adjust = TRUE, sigma = sigma),
  # For timepoint 5
  ref_grid(model_IIa_PR, trans = "logit", at = list(Time = 2)) %>% 
    summary(infer = c(F, F),  type = "response", 
            bias.adjust = TRUE, sigma = sigma),
  # For timepoint 0
  ref_grid(model_IIa_PR, trans = "logit", at = list(Time = 0)) %>% 
    summary(infer = c(F, F),  type = "response", 
            bias.adjust = TRUE, sigma = sigma)) %>% 
  mutate(Treatment = ifelse(Submergence, 'S', "C")) %>% 
  filter(!(Time == 0 & Submergence == 1))

# Figure S12
(ggplot(data = means_IIa_PR, aes(x = Time, y = response, colour = Treatment,
                                  shape = Treatment)) +
    geom_point(data = data_IIa_PR, mapping = aes(y = PetioleRatio), alpha = 0.5,
               size = 0.8, position = position_jitter(width = 0.3),
               show.legend = FALSE) + 
    geom_point() + geom_line() +
    geom_errorbar(aes(ymax = response + SE, ymin = response - SE),
                  width = 0.5) +
    facet_wrap(~Genotype) + 
    scale_x_continuous(breaks = c(0,2,5), labels = c("0","2","5")) +
    scale_colour_manual(labels = labelsIIa, values = colorsIIa) +
    scale_shape_manual(labels = labelsIIa, values = shapesIIa) +
    labs(y = "Petiole ratio") +
    theme_bw() +
    theme(panel.grid = element_blank())) %>% 
  ggsave(filename = "Results/Figure_S12.png", plot = ., dpi = 300, 
         width = 16, height = 8, units = "cm",scale = 1.25)

# Experiment IIa - Relative water content ---------------------------------


# Filter the data
data_IIa_RWC = XP_I_II %>% 
  filter(XP == 'II', Timepoint <= 3, !is.na(RWC)) %>% 
  mutate(Treatment = tnames[paste0(HighTemperature, Submergence, Drought)],
         Time = case_when(
           XP == "II" & Timepoint == 1 ~ 0,
           XP == "II" & Timepoint == 2 ~ 2,
           XP == "II" & Timepoint == 3 ~ 5))

# Fit linear mixed model
model_IIa_RWC = lme(logit(RWC)~Genotype*(Time + Submergence:Time),
                   random = ~ Time | Repetition, data = data_IIa_RWC,
                   control = lmeControl(maxIter = 1e4, msMaxIter = 1e4,
                                        opt = "optim"))

# F test on the fitted model
anova_IIa_RWC = anova(model_IIa_RWC, type = "marginal")
anova_IIa_RWC = cbind(Term = row.names(anova_IIa_RWC), anova_IIa_RWC)
row.names(anova_IIa_RWC) = NULL

# Marginal means
sigma = sqrt(sum(as.numeric(VarCorr(model_IIa_RWC)[c(1,3),1])))
means_IIa_RWC = bind_rows(
  # For timepoint 9
  ref_grid(model_IIa_RWC, trans = "logit", at = list(Time = 5)) %>% 
    summary(infer = c(F, F),  type = "response", 
            bias.adjust = TRUE, sigma = sigma),
  # For timepoint 5
  ref_grid(model_IIa_RWC, trans = "logit", at = list(Time = 2)) %>% 
    summary(infer = c(F, F),  type = "response", 
            bias.adjust = TRUE, sigma = sigma),
  # For timepoint 0
  ref_grid(model_IIa_RWC, trans = "logit", at = list(Time = 0)) %>% 
    summary(infer = c(F, F),  type = "response", 
            bias.adjust = TRUE, sigma = sigma)) %>% 
  mutate(Treatment = ifelse(Submergence, 'S', "C")) %>% 
  filter(!(Time == 0 & Submergence == 1))

# Figure S15
(ggplot(data = means_IIa_RWC, aes(x = Time, y = response, colour = Treatment,
                                 shape = Treatment)) +
    geom_point(data = data_IIa_RWC, mapping = aes(y = RWC), alpha = 0.5,
               size = 0.8, position = position_jitter(width = 0.3),
               show.legend = FALSE) + 
    geom_point() + geom_line() +
    geom_errorbar(aes(ymax = response + SE, ymin = response - SE),
                  width = 0.5) +
    facet_wrap(~Genotype) + 
    scale_x_continuous(breaks = c(0,2,5), labels = c("0","2","5")) +
    scale_y_continuous(limits = c(0.90,0.98), breaks = c(0.90,0.94,0.98)) +
    scale_colour_manual(labels = labelsIIa, values = colorsIIa) +
    scale_shape_manual(labels = labelsIIa, values = shapesIIa) +
    labs(y = "Relative water content") +
    theme_bw() +
    theme(panel.grid = element_blank())) %>% 
  ggsave(filename = "Results/Figure_S15.png", plot = ., dpi = 300, 
         width = 16, height = 8, units = "cm",scale = 1.25)

# Experiment IIa - Root length --------------------------------------------

# Filter the data
data_IIa_RL = XP_I_II %>% 
  filter(XP == 'II', Timepoint <= 3, !is.na(RootLength)) %>% 
  mutate(Treatment = tnames[paste0(HighTemperature, Submergence, Drought)],
         Time = case_when(
           XP == "II" & Timepoint == 1 ~ 0,
           XP == "II" & Timepoint == 2 ~ 2,
           XP == "II" & Timepoint == 3 ~ 5))

# Fit linear mixed model
model_IIa_RL = lme(log(RootLength)~Genotype*(Time + Submergence:Time),
                    random = ~ Time | Repetition, data = data_IIa_RL,
                    control = lmeControl(maxIter = 1e4, msMaxIter = 1e4,
                                         opt = "optim"))

# F test on the fitted model
anova_IIa_RL = anova(model_IIa_RL, type = "marginal")
anova_IIa_RL = cbind(Term = row.names(anova_IIa_RL), anova_IIa_RL)
row.names(anova_IIa_RL) = NULL

# Marginal means
sigma = sqrt(sum(as.numeric(VarCorr(model_IIa_RL)[c(1,3),1])))
means_IIa_RL = bind_rows(
  # For timepoint 9
  ref_grid(model_IIa_RL, trans = "log", at = list(Time = 5)) %>% 
    summary(infer = c(F, F),  type = "response", 
            bias.adjust = TRUE, sigma = sigma),
  # For timepoint 5
  ref_grid(model_IIa_RL, trans = "log", at = list(Time = 2)) %>% 
    summary(infer = c(F, F),  type = "response", 
            bias.adjust = TRUE, sigma = sigma),
  # For timepoint 0
  ref_grid(model_IIa_RL, trans = "log", at = list(Time = 0)) %>% 
    summary(infer = c(F, F),  type = "response", 
            bias.adjust = TRUE, sigma = sigma)) %>% 
  mutate(Treatment = ifelse(Submergence, 'S', "C")) %>% 
  filter(!(Time == 0 & Submergence == 1))

# Figure S18
(ggplot(data = means_IIa_RL, aes(x = Time, y = response, colour = Treatment,
                                  shape = Treatment)) +
    geom_point(data = data_IIa_RL, mapping = aes(y = RootLength), alpha = 0.5,
               size = 0.8, position = position_jitter(width = 0.3),
               show.legend = FALSE) + 
    geom_point() + geom_line() +
    geom_errorbar(aes(ymax = response + SE, ymin = response - SE),
                  width = 0.5) +
    facet_wrap(~Genotype) + 
    scale_x_continuous(breaks = c(0,2,5), labels = c("0","2","5")) +
    scale_colour_manual(labels = labelsIIa, values = colorsIIa) +
    scale_shape_manual(labels = labelsIIa, values = shapesIIa) +
    labs(y = quote(Root~length~(cm))) +
    theme_bw() +
    theme(panel.grid = element_blank())) %>% 
  ggsave(filename = "Results/Figure_S18.png", plot = ., dpi = 300, 
         width = 16, height = 8, units = "cm",scale = 1.25)

# Experiment IIa - Rosette dry weight -------------------------------------

# Filter the data
data_IIa_DW = XP_I_II %>% 
  filter(XP == 'II', Timepoint <= 3, !is.na(DW)) %>% 
  mutate(Treatment = tnames[paste0(HighTemperature, Submergence, Drought)],
         Time = case_when(
           XP == "II" & Timepoint == 1 ~ 0,
           XP == "II" & Timepoint == 2 ~ 2,
           XP == "II" & Timepoint == 3 ~ 5))

# Fit linear mixed model
model_IIa_DW = lme(log(DW)~Genotype*(Time + Submergence:Time),
                   random = ~ Time | Repetition, data = data_IIa_DW,
                   control = lmeControl(maxIter = 1e4, msMaxIter = 1e4,
                                        opt = "optim"))

# F test on the fitted model
anova_IIa_DW = anova(model_IIa_DW, type = "marginal")
anova_IIa_DW = cbind(Term = row.names(anova_IIa_DW), anova_IIa_DW)
row.names(anova_IIa_DW) = NULL

# Marginal means
sigma = sqrt(sum(as.numeric(VarCorr(model_IIa_DW)[c(1,3),1])))
means_IIa_DW = bind_rows(
  # For timepoint 9
  ref_grid(model_IIa_DW, trans = "log", at = list(Time = 5)) %>% 
    summary(infer = c(F, F),  type = "response", 
            bias.adjust = TRUE, sigma = sigma),
  # For timepoint 5
  ref_grid(model_IIa_DW, trans = "log", at = list(Time = 2)) %>% 
    summary(infer = c(F, F),  type = "response", 
            bias.adjust = TRUE, sigma = sigma),
  # For timepoint 0
  ref_grid(model_IIa_DW, trans = "log", at = list(Time = 0)) %>% 
    summary(infer = c(F, F),  type = "response", 
            bias.adjust = TRUE, sigma = sigma)) %>% 
  mutate(Treatment = ifelse(Submergence, 'S', "C")) %>% 
  filter(!(Time == 0 & Submergence == 1))

# Figure S21
(ggplot(data = means_IIa_DW, aes(x = Time, y = response, colour = Treatment,
                                 shape = Treatment)) +
    geom_point(data = data_IIa_DW, mapping = aes(y = DW), alpha = 0.5,
               size = 0.8, position = position_jitter(width = 0.3),
               show.legend = FALSE) + 
    geom_point() + geom_line() +
    geom_errorbar(aes(ymax = response + SE, ymin = response - SE),
                  width = 0.5) +
    facet_wrap(~Genotype) + 
    scale_x_continuous(breaks = c(0,2,5), labels = c("0","2","5")) +
    scale_y_continuous(trans = "log", breaks = c(0.4,1,2.75,7.2),
                       limits = c(0.35,9)) + 
    scale_colour_manual(labels = labelsIIa, values = colorsIIa) +
    scale_shape_manual(labels = labelsIIa, values = shapesIIa) +
    labs(y = "Rosette dry weight (mg)") +
    theme_bw() +
    theme(panel.grid = element_blank())) %>% 
  ggsave(filename = "Results/Figure_S21.png", plot = ., dpi = 300, 
         width = 16, height = 8, units = "cm",scale = 1.25)


# Experiment IIa - Specific leaf area -------------------------------------

# Filter the data
data_IIa_SLA = XP_I_II %>% 
  filter(XP == 'II', Timepoint <= 3, !is.na(SLA)) %>% 
  mutate(Treatment = tnames[paste0(HighTemperature, Submergence, Drought)],
         Time = case_when(
           XP == "II" & Timepoint == 1 ~ 0,
           XP == "II" & Timepoint == 2 ~ 2,
           XP == "II" & Timepoint == 3 ~ 5))

# Fit linear mixed model
model_IIa_SLA = lme(log(SLA)~Genotype*(Time + Submergence:Time),
                   random = ~ Time | Repetition, data = data_IIa_SLA,
                   control = lmeControl(maxIter = 1e4, msMaxIter = 1e4,
                                        opt = "optim"))

# F test on the fitted model
anova_IIa_SLA = anova(model_IIa_SLA, type = "marginal")
anova_IIa_SLA = cbind(Term = row.names(anova_IIa_SLA), anova_IIa_SLA)
row.names(anova_IIa_SLA) = NULL

# Marginal means
sigma = sqrt(sum(as.numeric(VarCorr(model_IIa_SLA)[c(1,3),1])))
means_IIa_SLA = bind_rows(
  # For timepoint 9
  ref_grid(model_IIa_SLA, trans = "log", at = list(Time = 5)) %>% 
    summary(infer = c(F, F),  type = "response", 
            bias.adjust = TRUE, sigma = sigma),
  # For timepoint 5
  ref_grid(model_IIa_SLA, trans = "log", at = list(Time = 2)) %>% 
    summary(infer = c(F, F),  type = "response", 
            bias.adjust = TRUE, sigma = sigma),
  # For timepoint 0
  ref_grid(model_IIa_SLA, trans = "log", at = list(Time = 0)) %>% 
    summary(infer = c(F, F),  type = "response", 
            bias.adjust = TRUE, sigma = sigma)) %>% 
  mutate(Treatment = ifelse(Submergence, 'S', "C")) %>% 
  filter(!(Time == 0 & Submergence == 1))

# Figure S27
(ggplot(data = means_IIa_SLA, aes(x = Time, y = response, colour = Treatment,
                                 shape = Treatment)) +
    geom_point(data = data_IIa_SLA, mapping = aes(y = SLA), alpha = 0.5,
               size = 0.8, position = position_jitter(width = 0.3),
               show.legend = FALSE) + 
    geom_point() + geom_line() +
    geom_errorbar(aes(ymax = response + SE, ymin = response - SE),
                  width = 0.5) +
    facet_wrap(~Genotype) + 
    scale_x_continuous(breaks = c(0,2,5), labels = c("0","2","5")) +
    scale_y_continuous(trans = "log", limits = c(300,2000),
                       breaks = c(300,450,800,1500)) + 
    scale_colour_manual(labels = labelsIIa, values = colorsIIa) +
    scale_shape_manual(labels = labelsIIa, values = shapesIIa) +
    labs(y = quote(Specific~leaf~area~(cm^2~g^{-1}))) +
    theme_bw() +
    theme(panel.grid = element_blank())) %>% 
  ggsave(filename = "Results/Figure_S27.png", plot = ., dpi = 300, 
         width = 16, height = 8, units = "cm",scale = 1.25)

# Experiment IIb - Blade Shape --------------------------------------------

# Filter the data
data_IIb_BS = XP_I_II %>% 
  filter(XP == 'II', Timepoint >= 3, !is.na(BladeShape)) %>% 
  mutate(Treatment = tnames[paste0(HighTemperature, Submergence, Drought)],
         Time = case_when(
           XP == "II" & Timepoint == 3 ~ 0,
           XP == "II" & Timepoint == 4 ~ 4,
           XP == "II" & Timepoint == 5 ~ 7))

# Fit linear mixed model
model_IIb_BS = lme(log(BladeShape)~Genotype*Submergence*(Time + Drought:Time),
                   random = ~ Time | Repetition, data = data_IIb_BS,
                   control = lmeControl(maxIter = 1e4, msMaxIter = 1e4,
                                        opt = "optim"))

# F test on the fitted model
anova_IIb_BS = anova(model_IIb_BS, type = "marginal")
anova_IIb_BS = cbind(Term = row.names(anova_IIb_BS), anova_IIb_BS)
row.names(anova_IIb_BS) = NULL

# Marginal means
sigma = sqrt(sum(as.numeric(VarCorr(model_IIb_BS)[c(1,3),1])))
means_IIb_BS = bind_rows(
  # For timepoint 9
  ref_grid(model_IIb_BS, trans = "log", at = list(Time = 7)) %>% 
    summary(infer = c(F, F),  type = "response", 
            bias.adjust = TRUE, sigma = sigma),
  # For timepoint 5
  ref_grid(model_IIb_BS, trans = "log", at = list(Time = 4)) %>% 
    summary(infer = c(F, F),  type = "response", 
            bias.adjust = TRUE, sigma = sigma),
  # For timepoint 0
  ref_grid(model_IIb_BS, trans = "log", at = list(Time = 0)) %>% 
    summary(infer = c(F, F),  type = "response", 
            bias.adjust = TRUE, sigma = sigma)) %>% 
  mutate(Treatment = tnamesII[paste0(Submergence, Drought)]) %>% 
  filter(!(Time == 0 & Drought == 1))

# Figure S4
(ggplot(data = means_IIb_BS, aes(x = Time, y = response, colour = Treatment,
                                 shape = Treatment)) +
    geom_point(data = data_IIb_BS, mapping = aes(y = BladeShape), alpha = 0.5,
               size = 0.8, position = position_jitter(width = 0.3),
               show.legend = FALSE) + 
    geom_point() + geom_line() +
    geom_errorbar(aes(ymax = response + SE, ymin = response - SE),
                  width = 0.5) +
    facet_wrap(~Genotype) + 
    scale_x_continuous(breaks = c(0,4,7), labels = c("5","9","12")) +
    scale_colour_manual(labels = labelsIIb, values = colorsIIb) +
    scale_shape_manual(labels = labelsIIb, values = shapesIIb) +
    labs(y = "Blade shape") +
    theme_bw() +
    theme(panel.grid = element_blank())) %>% 
  ggsave(filename = "Results/Figure_S4.png", plot = ., dpi = 300, 
         width = 16, height = 8, units = "cm",scale = 1.25)

# Experiment IIb - Early rosette leaf number ------------------------------

# Filter the data
data_IIb_LN = XP_I_II %>% 
  filter(XP == 'II', Timepoint >= 3, !is.na(LeafNumber)) %>% 
  mutate(Treatment = tnames[paste0(HighTemperature, Submergence, Drought)],
         Time = case_when(
           XP == "II" & Timepoint == 3 ~ 0,
           XP == "II" & Timepoint == 4 ~ 4,
           XP == "II" & Timepoint == 5 ~ 7))

# Fit linear mixed model
model_IIb_LN = lme(log(LeafNumber)~Genotype*Submergence*(Time + Drought:Time),
                   random = ~ Time | Repetition, data = data_IIb_LN,
                   control = lmeControl(maxIter = 1e4, msMaxIter = 1e4,
                                        opt = "optim"))

# F test on the fitted model
anova_IIb_LN = anova(model_IIb_LN, type = "marginal")
anova_IIb_LN = cbind(Term = row.names(anova_IIb_LN), anova_IIb_LN)
row.names(anova_IIb_LN) = NULL

# Marginal means
sigma = sqrt(sum(as.numeric(VarCorr(model_IIb_LN)[c(1,3),1])))
means_IIb_LN = bind_rows(
  # For timepoint 9
  ref_grid(model_IIb_LN, trans = "log", at = list(Time = 7)) %>% 
    summary(infer = c(F, F),  type = "response", 
            bias.adjust = TRUE, sigma = sigma),
  # For timepoint 5
  ref_grid(model_IIb_LN, trans = "log", at = list(Time = 4)) %>% 
    summary(infer = c(F, F),  type = "response", 
            bias.adjust = TRUE, sigma = sigma),
  # For timepoint 0
  ref_grid(model_IIb_LN, trans = "log", at = list(Time = 0)) %>% 
    summary(infer = c(F, F),  type = "response", 
            bias.adjust = TRUE, sigma = sigma)) %>% 
  mutate(Treatment = tnamesII[paste0(Submergence, Drought)]) %>% 
  filter(!(Time == 0 & Drought == 1))

# Figure S22
(ggplot(data = means_IIb_LN, aes(x = Time, y = response, colour = Treatment,
                                 shape = Treatment)) +
    geom_point(data = data_IIb_LN, mapping = aes(y = LeafNumber), alpha = 0.5,
               size = 0.8, position = position_jitter(width = 0.3),
               show.legend = FALSE) + 
    geom_point() + geom_line() +
    geom_errorbar(aes(ymax = response + SE, ymin = response - SE),
                  width = 0.5) +
    facet_wrap(~Genotype) + 
    scale_x_continuous(breaks = c(0,4,7), labels = c("5","9","12")) +
    scale_colour_manual(labels = labelsIIb, values = colorsIIb) +
    scale_shape_manual(labels = labelsIIb, values = shapesIIb) +
    labs(y = "Early rosette leaf number") +
    theme_bw() +
    theme(panel.grid = element_blank())) %>% 
  ggsave(filename = "Results/Figure_S22.png", plot = ., dpi = 300, 
         width = 16, height = 8, units = "cm",scale = 1.25)


# Experiment IIb - Leaf angle ---------------------------------------------

# Filter the data
data_IIb_Angle = XP_I_II %>% 
  filter(XP == 'II', Timepoint >= 3, !is.na(Angle), Angle != 0) %>% 
  mutate(Treatment = tnames[paste0(HighTemperature, Submergence, Drought)],
         Time = case_when(
           XP == "II" & Timepoint == 3 ~ 0,
           XP == "II" & Timepoint == 4 ~ 4,
           XP == "II" & Timepoint == 5 ~ 7)) %>% 
  mutate(Angle = Angle/90)

# Fit linear mixed model
model_IIb_Angle = lme(logit(Angle)~Genotype*Submergence*(Time + Drought:Time),
                   random = ~ Time | Repetition, data = data_IIb_Angle,
                   control = lmeControl(maxIter = 1e4, msMaxIter = 1e4,
                                        opt = "optim"))

# F test on the fitted model
anova_IIb_Angle = anova(model_IIb_Angle, type = "marginal")
anova_IIb_Angle = cbind(Term = row.names(anova_IIb_Angle), anova_IIb_Angle)
row.names(anova_IIb_Angle) = NULL

# Marginal means
sigma = sqrt(sum(as.numeric(VarCorr(model_IIb_Angle)[c(1,3),1])))
means_IIb_Angle = bind_rows(
  # For timepoint 9
  ref_grid(model_IIb_Angle, trans = "logit", at = list(Time = 7)) %>% 
    summary(infer = c(F, F),  type = "response", 
            bias.adjust = TRUE, sigma = sigma),
  # For timepoint 5
  ref_grid(model_IIb_Angle, trans = "logit", at = list(Time = 4)) %>% 
    summary(infer = c(F, F),  type = "response", 
            bias.adjust = TRUE, sigma = sigma),
  # For timepoint 0
  ref_grid(model_IIb_Angle, trans = "logit", at = list(Time = 0)) %>% 
    summary(infer = c(F, F),  type = "response", 
            bias.adjust = TRUE, sigma = sigma)) %>% 
  mutate(Treatment = tnamesII[paste0(Submergence, Drought)]) %>% 
  filter(!(Time == 0 & Drought == 1))

# Figure S7
(ggplot(means_IIb_Angle, aes(x = Time, y = response*90, 
                colour = Treatment, shape = Treatment)) +
    geom_point(data = data_IIb_Angle, mapping = aes(y = Angle*90), alpha = 0.5,
               size = 0.8, position = position_jitter(width = 0.3),
               show.legend = FALSE) + 
    geom_point() + geom_line() +
    geom_errorbar(aes(ymax = (response + SE)*90, 
                      ymin = (response - SE)*90),
                  width = 0.5) +
    facet_wrap(~Genotype) + 
    scale_x_continuous(breaks = c(0,4,7), labels = c("5","9","12")) +
    scale_colour_manual(labels = labelsIIb, values = colorsIIb) +
    scale_shape_manual(labels = labelsIIb, values = shapesIIb) +
    labs(y = quote(Leaf~angle~(degree))) +
    theme_bw() +
    theme(panel.grid = element_blank())) %>% 
  ggsave(filename = "Results/Figure_S7.png", plot = ., dpi = 300, 
         width = 16, height = 8, units = "cm",scale = 1.25)


# Experiment IIb - Leaf size --------------------------------------

# Filter the data
data_IIb_LS = XP_I_II %>% 
  mutate(LS = PlantArea/LeafNumber) %>% 
  filter(XP == 'II', Timepoint >= 3, !is.na(LS)) %>% 
  mutate(Treatment = tnames[paste0(HighTemperature, Submergence, Drought)],
         Time = case_when(
           XP == "II" & Timepoint == 3 ~ 0,
           XP == "II" & Timepoint == 4 ~ 4,
           XP == "II" & Timepoint == 5 ~ 7))

# Fit linear mixed model
model_IIb_LS = lme(log(LS)~Genotype*Submergence*(Time + Drought:Time),
                   random = ~ Time | Repetition, data = data_IIb_LS,
                   control = lmeControl(maxIter = 1e4, msMaxIter = 1e4,
                                        opt = "optim"))

# F test on the fitted model
anova_IIb_LS = anova(model_IIb_LS, type = "marginal")
anova_IIb_LS = cbind(Term = row.names(anova_IIb_LS), anova_IIb_LS)
row.names(anova_IIb_LS) = NULL

# Marginal means
sigma = sqrt(sum(as.numeric(VarCorr(model_IIb_LS)[c(1,3),1])))
means_IIb_LS = bind_rows(
  # For timepoint 9
  ref_grid(model_IIb_LS, trans = "log", at = list(Time = 7)) %>% 
    summary(infer = c(F, F),  type = "response", 
            bias.adjust = TRUE, sigma = sigma),
  # For timepoint 5
  ref_grid(model_IIb_LS, trans = "log", at = list(Time = 4)) %>% 
    summary(infer = c(F, F),  type = "response", 
            bias.adjust = TRUE, sigma = sigma),
  # For timepoint 0
  ref_grid(model_IIb_LS, trans = "log", at = list(Time = 0)) %>% 
    summary(infer = c(F, F),  type = "response", 
            bias.adjust = TRUE, sigma = sigma)) %>% 
  mutate(Treatment = tnamesII[paste0(Submergence, Drought)]) %>% 
  filter(!(Time == 0 & Drought == 1))

# Figure S10
(ggplot(data = means_IIb_LS, aes(x = Time, y = response, colour = Treatment,
                                 shape = Treatment)) +
    geom_point(data = data_IIb_LS, mapping = aes(y = LS), alpha = 0.5,
               size = 0.8, position = position_jitter(width = 0.3),
               show.legend = FALSE) + 
    geom_point() + geom_line() +
    geom_errorbar(aes(ymax = response + SE, ymin = response - SE),
                  width = 0.5) +
    facet_wrap(~Genotype) + 
    scale_x_continuous(breaks = c(0,4,7), labels = c("5","9","12")) +
    scale_y_continuous(trans = "log", breaks = c(0.05,0.15,0.35,1)) + 
    scale_colour_manual(labels = labelsIIb, values = colorsIIb) +
    scale_shape_manual(labels = labelsIIb, values = shapesIIb) +
    labs(y = quote(Leaf~size~(cm^2))) +
    theme_bw() +
    theme(panel.grid = element_blank())) %>% 
  ggsave(filename = "Results/Figure_S10.png", plot = ., dpi = 300, 
         width = 16, height = 8, units = "cm",scale = 1.25)


# Experiment IIb - Petiole ratio ------------------------------------------

# Filter the data
data_IIb_PR = XP_I_II %>% 
  filter(XP == 'II', Timepoint >= 3, !is.na(PetioleRatio)) %>% 
  mutate(Treatment = tnames[paste0(HighTemperature, Submergence, Drought)],
         Time = case_when(
           XP == "II" & Timepoint == 3 ~ 0,
           XP == "II" & Timepoint == 4 ~ 4,
           XP == "II" & Timepoint == 5 ~ 7))

# Fit linear mixed model
model_IIb_PR = lme(logit(PetioleRatio)~Genotype*Submergence*(Time + Drought:Time),
                      random = ~ Time | Repetition, data = data_IIb_PR,
                      control = lmeControl(maxIter = 1e4, msMaxIter = 1e4,
                                           opt = "optim"))

# F test on the fitted model
anova_IIb_PR = anova(model_IIb_PR, type = "marginal")
anova_IIb_PR = cbind(Term = row.names(anova_IIb_PR), anova_IIb_PR)
row.names(anova_IIb_PR) = NULL

# Marginal means
sigma = sqrt(sum(as.numeric(VarCorr(model_IIb_PR)[c(1,3),1])))
means_IIb_PR = bind_rows(
  # For timepoint 9
  ref_grid(model_IIb_PR, trans = "logit", at = list(Time = 7)) %>% 
    summary(infer = c(F, F),  type = "response", 
            bias.adjust = TRUE, sigma = sigma),
  # For timepoint 5
  ref_grid(model_IIb_PR, trans = "logit", at = list(Time = 4)) %>% 
    summary(infer = c(F, F),  type = "response", 
            bias.adjust = TRUE, sigma = sigma),
  # For timepoint 0
  ref_grid(model_IIb_PR, trans = "logit", at = list(Time = 0)) %>% 
    summary(infer = c(F, F),  type = "response", 
            bias.adjust = TRUE, sigma = sigma)) %>% 
  mutate(Treatment = tnamesII[paste0(Submergence, Drought)]) %>% 
  filter(!(Time == 0 & Drought == 1))

# Figure S13
(ggplot(means_IIb_PR, aes(x = Time, y = response, 
                             colour = Treatment, shape = Treatment)) +
    geom_point(data = data_IIb_PR, mapping = aes(y = PetioleRatio), alpha = 0.5,
               size = 0.8, position = position_jitter(width = 0.3),
               show.legend = FALSE) + 
    geom_point() + geom_line() +
    geom_errorbar(aes(ymax = (response + SE), 
                      ymin = (response - SE)),
                  width = 0.5) +
    facet_wrap(~Genotype) + 
    scale_x_continuous(breaks = c(0,4,7), labels = c("5","9","12")) +
    scale_y_continuous(limits = c(0.35,0.6)) + 
    scale_colour_manual(labels = labelsIIb, values = colorsIIb) +
    scale_shape_manual(labels = labelsIIb, values = shapesIIb) +
    labs(y = "Petiole ratio") +
    theme_bw() +
    theme(panel.grid = element_blank())) %>% 
  ggsave(filename = "Results/Figure_S13.png", plot = ., dpi = 300, 
         width = 16, height = 8, units = "cm",scale = 1.25)

# Experiment IIb - Relative water content ---------------------------------

# Filter the data
data_IIb_RWC = XP_I_II %>% 
  filter(XP == 'II', Timepoint >= 3, !is.na(RWC)) %>% 
  mutate(Treatment = tnames[paste0(HighTemperature, Submergence, Drought)],
         Time = case_when(
           XP == "II" & Timepoint == 3 ~ 0,
           XP == "II" & Timepoint == 4 ~ 4,
           XP == "II" & Timepoint == 5 ~ 7))

# Fit linear mixed model
model_IIb_RWC = lme(logit(RWC)~Genotype*Submergence*(Time + Drought:Time),
                   random = ~ Time | Repetition, data = data_IIb_RWC,
                   control = lmeControl(maxIter = 1e4, msMaxIter = 1e4,
                                        opt = "optim"))

# F test on the fitted model
anova_IIb_RWC = anova(model_IIb_RWC, type = "marginal")
anova_IIb_RWC = cbind(Term = row.names(anova_IIb_RWC), anova_IIb_RWC)
row.names(anova_IIb_RWC) = NULL

# Marginal means
sigma = sqrt(sum(as.numeric(VarCorr(model_IIb_RWC)[c(1,3),1])))
means_IIb_RWC = bind_rows(
  # For timepoint 9
  ref_grid(model_IIb_RWC, trans = "logit", at = list(Time = 7)) %>% 
    summary(infer = c(F, F),  type = "response", 
            bias.adjust = TRUE, sigma = sigma),
  # For timepoint 5
  ref_grid(model_IIb_RWC, trans = "logit", at = list(Time = 4)) %>% 
    summary(infer = c(F, F),  type = "response", 
            bias.adjust = TRUE, sigma = sigma),
  # For timepoint 0
  ref_grid(model_IIb_RWC, trans = "logit", at = list(Time = 0)) %>% 
    summary(infer = c(F, F),  type = "response", 
            bias.adjust = TRUE, sigma = sigma)) %>% 
  mutate(Treatment = tnamesII[paste0(Submergence, Drought)]) %>% 
  filter(!(Time == 0 & Drought == 1))

# Figure S16
(ggplot(means_IIb_RWC, aes(x = Time, y = response, 
                          colour = Treatment, shape = Treatment)) +
    geom_point(data = data_IIb_RWC, mapping = aes(y = RWC), alpha = 0.5,
               size = 0.8, position = position_jitter(width = 0.3),
               show.legend = FALSE) + 
    geom_point() + geom_line() +
    geom_errorbar(aes(ymax = (response + SE), 
                      ymin = (response - SE)),
                  width = 0.5) +
    facet_wrap(~Genotype) + 
    scale_x_continuous(breaks = c(0,4,7), labels = c("5","9","12")) +
    scale_colour_manual(labels = labelsIIb, values = colorsIIb) +
    scale_shape_manual(labels = labelsIIb, values = shapesIIb) +
    labs(y = "Relative water content") +
    theme_bw() +
    theme(panel.grid = element_blank())) %>% 
  ggsave(filename = "Results/Figure_S16.png", plot = ., dpi = 300, 
         width = 16, height = 8, units = "cm",scale = 1.25)

# Experiment IIb - Root length --------------------------------------------

# Filter the data
data_IIb_RL = XP_I_II %>% 
  filter(XP == 'II', Timepoint >= 3, !is.na(RootLength)) %>% 
  mutate(Treatment = tnames[paste0(HighTemperature, Submergence, Drought)],
         Time = case_when(
           XP == "II" & Timepoint == 3 ~ 0,
           XP == "II" & Timepoint == 4 ~ 4,
           XP == "II" & Timepoint == 5 ~ 7))

# Fit linear mixed model
model_IIb_RL = lme(log(RootLength)~Genotype*Submergence*(Time + Drought:Time),
                    random = ~ Time | Repetition, data = data_IIb_RL,
                    control = lmeControl(maxIter = 1e4, msMaxIter = 1e4,
                                         opt = "optim"))

# F test on the fitted model
anova_IIb_RL = anova(model_IIb_RL, type = "marginal")
anova_IIb_RL = cbind(Term = row.names(anova_IIb_RL), anova_IIb_RL)
row.names(anova_IIb_RL) = NULL

# Marginal means
sigma = sqrt(sum(as.numeric(VarCorr(model_IIb_RL)[c(1,3),1])))
means_IIb_RL = bind_rows(
  # For timepoint 9
  ref_grid(model_IIb_RL, trans = "log", at = list(Time = 7)) %>% 
    summary(infer = c(F, F),  type = "response", 
            bias.adjust = TRUE, sigma = sigma),
  # For timepoint 5
  ref_grid(model_IIb_RL, trans = "log", at = list(Time = 4)) %>% 
    summary(infer = c(F, F),  type = "response", 
            bias.adjust = TRUE, sigma = sigma),
  # For timepoint 0
  ref_grid(model_IIb_RL, trans = "log", at = list(Time = 0)) %>% 
    summary(infer = c(F, F),  type = "response", 
            bias.adjust = TRUE, sigma = sigma)) %>% 
  mutate(Treatment = tnamesII[paste0(Submergence, Drought)]) %>% 
  filter(!(Time == 0 & Drought == 1))

# Figure S19
(ggplot(data = means_IIb_RL, aes(x = Time, y = response, colour = Treatment,
                                  shape = Treatment)) +
    geom_point(data = data_IIb_RL, mapping = aes(y = RootLength),alpha=0.5,
               size = 0.8, position = position_jitter(width = 0.3),
               show.legend = FALSE) + 
    geom_point() + geom_line() +
    geom_errorbar(aes(ymax = response + SE, ymin = response - SE),
                  width = 0.5) +
    facet_wrap(~Genotype) + 
    scale_x_continuous(breaks = c(0,4,7), labels = c("5","9","12")) +
    scale_y_continuous(trans = "log", breaks = c(2,3,5,9,15)) + 
    scale_colour_manual(labels = labelsIIb, values = colorsIIb) +
    scale_shape_manual(labels = labelsIIb, values = shapesIIb) +
    labs(y = quote(Root~length~(cm))) +
    theme_bw() +
    theme(panel.grid = element_blank())) %>% 
  ggsave(filename = "Results/Figure_S19.png", plot = ., dpi = 300, 
         width = 16, height = 8, units = "cm",scale = 1.25)

# Experiment IIb - Rosette dry weight -------------------------------------

# Filter the data
data_IIb_DW = XP_I_II %>% 
  filter(XP == 'II', Timepoint >= 3, !is.na(DW)) %>% 
  mutate(Treatment = tnames[paste0(HighTemperature, Submergence, Drought)],
         Time = case_when(
           XP == "II" & Timepoint == 3 ~ 0,
           XP == "II" & Timepoint == 4 ~ 4,
           XP == "II" & Timepoint == 5 ~ 7))

# Fit linear mixed model
model_IIb_DW = lme(log(DW)~Genotype*Submergence*(Time + Drought:Time),
                   random = ~ Time | Repetition, data = data_IIb_DW,
                   control = lmeControl(maxIter = 1e4, msMaxIter = 1e4,
                                        opt = "optim"))

# F test on the fitted model
anova_IIb_DW = anova(model_IIb_DW, type = "marginal")
anova_IIb_DW = cbind(Term = row.names(anova_IIb_DW), anova_IIb_DW)
row.names(anova_IIb_DW) = NULL

# Marginal means
sigma = sqrt(sum(as.numeric(VarCorr(model_IIb_DW)[c(1,3),1])))
means_IIb_DW = bind_rows(
  # For timepoint 9
  ref_grid(model_IIb_DW, trans = "log", at = list(Time = 7)) %>% 
    summary(infer = c(F, F),  type = "response", 
            bias.adjust = TRUE, sigma = sigma),
  # For timepoint 5
  ref_grid(model_IIb_DW, trans = "log", at = list(Time = 4)) %>% 
    summary(infer = c(F, F),  type = "response", 
            bias.adjust = TRUE, sigma = sigma),
  # For timepoint 0
  ref_grid(model_IIb_DW, trans = "log", at = list(Time = 0)) %>% 
    summary(infer = c(F, F),  type = "response", 
            bias.adjust = TRUE, sigma = sigma)) %>% 
  mutate(Treatment = tnamesII[paste0(Submergence, Drought)]) %>% 
  filter(!(Time == 0 & Drought == 1))

# Figure S25
(ggplot(data = means_IIb_DW, aes(x = Time, y = response, colour = Treatment,
                                 shape = Treatment)) +
    geom_point(data = data_IIb_DW, mapping = aes(y = DW),alpha=0.5,
               size = 0.8, position = position_jitter(width = 0.3),
               show.legend = FALSE) + 
    geom_point() + geom_line() +
    geom_errorbar(aes(ymax = response + SE, ymin = response - SE),
                  width = 0.5) +
    facet_wrap(~Genotype) + 
    scale_x_continuous(breaks = c(0,4,7), labels = c("5","9","12")) +
    scale_y_continuous(trans = "log", breaks = c(0.3,1,2.5,7.5,20)) +
    scale_colour_manual(labels = labelsIIb, values = colorsIIb) +
    scale_shape_manual(labels = labelsIIb, values = shapesIIb) +
    labs(y = "Rosette dry weight (mg)") +
    theme_bw() +
    theme(panel.grid = element_blank())) %>% 
  ggsave(filename = "Results/Figure_S25.png", plot = ., dpi = 300, 
         width = 16, height = 8, units = "cm",scale = 1.25)

# Experiment IIb - Specific leaf area -------------------------------------

# Filter the data
data_IIb_SLA = XP_I_II %>% 
  filter(XP == 'II', Timepoint >= 3, !is.na(SLA)) %>% 
  mutate(Treatment = tnames[paste0(HighTemperature, Submergence, Drought)],
         Time = case_when(
           XP == "II" & Timepoint == 3 ~ 0,
           XP == "II" & Timepoint == 4 ~ 4,
           XP == "II" & Timepoint == 5 ~ 7))

# Fit linear mixed model
model_IIb_SLA = lme(log(SLA)~Genotype*Submergence*(Time + Drought:Time),
                   random = ~ Time | Repetition, data = data_IIb_SLA,
                   control = lmeControl(maxIter = 1e4, msMaxIter = 1e4,
                                        opt = "optim"))

# F test on the fitted model
anova_IIb_SLA = anova(model_IIb_SLA, type = "marginal")
anova_IIb_SLA = cbind(Term = row.names(anova_IIb_SLA), anova_IIb_SLA)
row.names(anova_IIb_SLA) = NULL

# Marginal means
sigma = sqrt(sum(as.numeric(VarCorr(model_IIb_SLA)[c(1,3),1])))
means_IIb_SLA = bind_rows(
  # For timepoint 9
  ref_grid(model_IIb_SLA, trans = "log", at = list(Time = 7)) %>% 
    summary(infer = c(F, F),  type = "response", 
            bias.adjust = TRUE, sigma = sigma),
  # For timepoint 5
  ref_grid(model_IIb_SLA, trans = "log", at = list(Time = 4)) %>% 
    summary(infer = c(F, F),  type = "response", 
            bias.adjust = TRUE, sigma = sigma),
  # For timepoint 0
  ref_grid(model_IIb_SLA, trans = "log", at = list(Time = 0)) %>% 
    summary(infer = c(F, F),  type = "response", 
            bias.adjust = TRUE, sigma = sigma)) %>% 
  mutate(Treatment = tnamesII[paste0(Submergence, Drought)]) %>% 
  filter(!(Time == 0 & Drought == 1))

# Figure S28
(ggplot(data = means_IIb_SLA, aes(x = Time, y = response, colour = Treatment,
                                 shape = Treatment)) +
    geom_point(data = data_IIb_SLA, mapping = aes(y = SLA),alpha=0.5,
               size = 0.8, position = position_jitter(width = 0.3),
               show.legend = FALSE) + 
    geom_point() + geom_line() +
    geom_errorbar(aes(ymax = response + SE, ymin = response - SE),
                  width = 0.5) +
    facet_wrap(~Genotype) + 
    scale_x_continuous(breaks = c(0,4,7), labels = c("5","9","12")) +
    scale_y_continuous(trans = "log", limits = c(300,2000),
                       breaks = c(300,450,800,1500)) + 
    scale_colour_manual(labels = labelsIIb, values = colorsIIb) +
    scale_shape_manual(labels = labelsIIb, values = shapesIIb) +
    labs(y = quote(Specific~leaf~area~(cm^2~g^{-1}))) +
    theme_bw() +
    theme(panel.grid = element_blank())) %>% 
  ggsave(filename = "Results/Figure_S28.png", plot = ., dpi = 300, 
         width = 16, height = 8, units = "cm",scale = 1.25)

# Experiment III - Flowering time -----------------------------------------

# Filter the data
data_III_FT = XP_III %>% filter(!is.na(FT))

# Fit linear mixed model
model_III_FT = lme(log(FT)~Genotype*Drought*(HighTemperature + Submergence),
                   random = ~ 1 | Repetition, data = data_III_FT,
                   control = lmeControl(maxIter = 1e4, msMaxIter = 1e4, 
                                        opt = "optim"))

# F test on the fitted model
anova_III_FT = anova(model_III_FT, type = "marginal")
anova_III_FT = cbind(Term = row.names(anova_III_FT), anova_III_FT)
row.names(anova_III_FT) = NULL

# Experiment III - Total rosette leaf number ------------------------------

# Filter the data
data_III_TLN = XP_III %>% filter(!is.na(LN))

# Fit linear mixed model
model_III_TLN = lme(log(LN)~Genotype*Drought*(HighTemperature + Submergence),
                    random = ~ 1 | Repetition, data = data_III_TLN,
                    control = lmeControl(maxIter = 1e4, msMaxIter = 1e4,
                                 opt = "optim"))

# F test on the fitted model
anova_III_TLN = anova(model_III_TLN, type = "marginal")
anova_III_TLN = cbind(Term = row.names(anova_III_TLN), anova_III_TLN)
row.names(anova_III_TLN) = NULL


# Experiment III - Yield per plant ----------------------------------------

# Filter the data
data_III_Y = XP_III %>% filter(!is.na(Yield))

# Fit linear mixed model
model_III_Y = lme(log(Yield)~Genotype*Drought*(HighTemperature + Submergence),
                    random = ~ 1 | Repetition, data = data_III_Y,
                    control = lmeControl(maxIter = 1e4, msMaxIter = 1e4,
                                         opt = "optim"))

# F test on the fitted model
anova_III_Y = anova(model_III_Y, type = "marginal")
anova_III_Y = cbind(Term = row.names(anova_III_Y), anova_III_Y)
row.names(anova_III_Y) = NULL


# Figure 2 ----------------------------------------------------------------

# LS
int_I_LS = model_I_LS %>% 
        ref_grid(trans = "log", at = list(Time = 0, Drought = 0)) %>% 
              emmeans:::cld.emmGrid(type = "response", Letters = letters, 
                                    by = "Genotype", sort = FALSE, bias.adjust = TRUE, 
                                    sigma = sigma(model_I_LS)) %>% 
        as_tibble %>% 
        rename(group = .group, intercept = response) %>% 
        mutate(Treatment = ifelse(HighTemperature == 1, "HT", "C"),
               group = str_replace_all(string=group, pattern=" ", repl="")) %>% 
        mutate(Trait = "LS")
              
# Angle
int_I_Angle = model_I_Angle %>% 
  ref_grid(trans = "logit", at = list(Time = 0, Drought = 0)) %>% 
  emmeans:::cld.emmGrid(type = "response", Letters = letters, 
                        by = "Genotype", sort = FALSE, bias.adjust = TRUE, 
                        sigma = sigma(model_I_Angle)) %>% 
  as_tibble %>% 
  rename(group = .group, intercept = response) %>% 
  mutate(Treatment = ifelse(HighTemperature == 1, "HT", "C"),
         group = str_replace_all(string=group, pattern=" ", repl=""),
         intercept = intercept*90,
         SE = SE*90) %>% 
  mutate(Trait = "Angle")

# DW
int_I_DW = model_I_DW %>% 
  ref_grid(trans = "log", at = list(Time = 0, Drought = 0)) %>% 
  emmeans:::cld.emmGrid(type = "response", Letters = letters, 
                        by = "Genotype", sort = FALSE, bias.adjust = TRUE, 
                        sigma = sigma(model_I_DW)) %>% 
  as_tibble %>% 
  rename(group = .group, intercept = response) %>% 
  mutate(Treatment = ifelse(HighTemperature == 1, "HT", "C"),
         group = str_replace_all(string=group, pattern=" ", repl="")) %>% 
  mutate(Trait = "DW")

# LN
int_I_LN = model_I_LN %>% 
  ref_grid(trans = "log", at = list(Time = 0, Drought = 0)) %>% 
  emmeans:::cld.emmGrid(type = "response", Letters = letters, 
                        by = "Genotype", sort = FALSE, bias.adjust = TRUE, 
                        sigma = sigma(model_I_LN)) %>% 
  as_tibble %>% 
  rename(group = .group, intercept = response) %>% 
  mutate(Treatment = ifelse(HighTemperature == 1, "HT", "C"),
         group = str_replace_all(string=group, pattern=" ", repl="")) %>% 
  mutate(Trait = "LN")

# PR
int_I_PR = model_I_PR %>% 
  ref_grid(trans = "logit", at = list(Time = 0, Drought = 0)) %>% 
  emmeans:::cld.emmGrid(type = "response", Letters = letters, 
                        by = "Genotype", sort = FALSE, bias.adjust = TRUE, 
                        sigma = sigma(model_I_PR)) %>% 
  as_tibble %>% 
  rename(group = .group, intercept = response) %>% 
  mutate(Treatment = ifelse(HighTemperature == 1, "HT", "C"),
         group = str_replace_all(string=group, pattern=" ", repl="")) %>% 
  mutate(Trait = "PR")

# RL
int_I_RL = model_I_RL %>% 
  ref_grid(trans = "log", at = list(Time = 0, Drought = 0)) %>% 
  emmeans:::cld.emmGrid(type = "response", Letters = letters, 
                        by = "Genotype", sort = FALSE, bias.adjust = TRUE, 
                        sigma = sigma(model_I_RL)) %>% 
  as_tibble %>% 
  rename(group = .group, intercept = response) %>% 
  mutate(Treatment = ifelse(HighTemperature == 1, "HT", "C"),
         group = str_replace_all(string=group, pattern=" ", repl="")) %>% 
  mutate(Trait = "RL")

# RWC
int_I_RWC = model_I_RWC %>% 
  ref_grid(trans = "logit", at = list(Time = 0, Drought = 0)) %>% 
  emmeans:::cld.emmGrid(type = "response", Letters = letters, 
                        by = "Genotype", sort = FALSE, bias.adjust = TRUE, 
                        sigma = sigma(model_I_RWC)) %>% 
  as_tibble %>% 
  rename(group = .group, intercept = response) %>% 
  mutate(Treatment = ifelse(HighTemperature == 1, "HT", "C"),
         group = str_replace_all(string=group, pattern=" ", repl="")) %>% 
  mutate(Trait = "RWC")

# BS
int_I_BS = model_I_BS %>% 
  ref_grid(trans = "log", at = list(Time = 0, Drought = 0)) %>% 
  emmeans:::cld.emmGrid(type = "response", Letters = letters, 
                        by = "Genotype", sort = FALSE, bias.adjust = TRUE, 
                        sigma = sigma(model_I_BS)) %>% 
  as_tibble %>% 
  rename(group = .group, intercept = response) %>% 
  mutate(Treatment = ifelse(HighTemperature == 1, "HT", "C"),
         group = str_replace_all(string=group, pattern=" ", repl="")) %>% 
  mutate(Trait = "BS")

# SLA
int_I_SLA = model_I_SLA %>% 
  ref_grid(trans = "log", at = list(Time = 0, Drought = 0)) %>% 
  emmeans:::cld.emmGrid(type = "response", Letters = letters, 
                        by = "Genotype", sort = FALSE, bias.adjust = TRUE, 
                        sigma = sigma(model_I_SLA)) %>% 
  as_tibble %>% 
  rename(group = .group, intercept = response) %>% 
  mutate(Treatment = ifelse(HighTemperature == 1, "HT", "C"),
         group = str_replace_all(string=group, pattern=" ", repl="")) %>% 
  mutate(Trait = "SLA")

intercepts_I = bind_rows(int_I_Angle, int_I_DW, int_I_LN, int_I_LS,
                         int_I_PR, int_I_RL, int_I_RWC, int_I_BS, int_I_SLA)

intercepts_I = intercepts_I %>% 
  group_by(Trait) %>% 
  mutate(ylabel = max(intercept + SE)*1.05)

vlines = intercepts_I %>% 
  group_by(Trait) %>% 
  summarise(xintercept = c(01.5,2.5,3.5))

rm(int_I_LS, int_I_Angle, int_I_DW, int_I_LN, int_I_PR, int_I_RL, int_I_RWC,
   int_I_BS, int_I_SLA)

pretty_labels = c(
  Angle = "Leaf~angle~(degree)",
  DW = "Rosette~dry~weight~(mg)",
  LN = "Early~rosette~leaf~number",
  LS = "Leaf~size~(cm^2)",
  PR = "Petiole~ratio",
  RL = "Root~length~(cm)",
  RWC = "Relative~water~content",
  BS = "Blade~shape",
  SLA = "Specific~leaf~area~(cm^2~g^{-1})")

intercepts_I = intercepts_I %>% 
  mutate(TraitPlot = pretty_labels[Trait])


(ggplot(data = intercepts_I, aes(x = Genotype, y = intercept, 
                                colour = Treatment, shape = Treatment)) +
  geom_point(position = position_dodge(width = 1)) +
  geom_errorbar(aes(ymax = intercept + SE, ymin = intercept - SE),
                position = position_dodge(width = 1)) +
  geom_text(aes(y = ylabel, label = group), size = 3,
            position = position_dodge(width = 1), show.legend = FALSE) +
  geom_vline(data = vlines, aes(xintercept = xintercept), col = BLACK) +
  facet_wrap(~TraitPlot, scales = "free",ncol = 3, labeller = label_parsed) +
  scale_colour_manual(labels = labelsI[c("C","HT")],
                      values = colorsI[c("C","HT")])  +
  scale_shape_manual(labels = labelsI[c("C","HT")],
                     values = shapesI[c("C","HT")])  +
  theme_bw() +
  theme(panel.grid = element_blank(),
        legend.position = "bottom",
        axis.title = element_blank())) %>% 
  ggsave("Results/Figure_2.png", plot = ., dpi = 300, 
         width = 16, height = 16, units = "cm",scale = 1.2)

# Figure 3 ----------------------------------------------------------------

# LS
slp_I_LS = model_I_LS %>% 
  emtrends(~ Genotype*HighTemperature*Drought,
           var = "Time", by = "Genotype") %>% 
  emmeans:::cld.emmGrid(by = "Genotype",
                        Letters = letters, sort = FALSE) %>% 
  as_tibble %>% 
  rename(slope = Time.trend, group = .group) %>% 
  mutate(Treatment = tnamesI[paste0(HighTemperature, Drought)],
         group = str_replace_all(string=group, pattern=" ", repl="")) %>% 
  mutate(Trait = "LS")

# Angle
slp_I_Angle = model_I_Angle %>% 
  emtrends(~ Genotype*HighTemperature*Drought,
           var = "Time", by = "Genotype") %>% 
  emmeans:::cld.emmGrid(by = "Genotype",
                        Letters = letters, sort = FALSE) %>% 
  as_tibble %>% 
  rename(slope = Time.trend, group = .group) %>% 
  mutate(Treatment = tnamesI[paste0(HighTemperature, Drought)],
         group = str_replace_all(string=group, pattern=" ", repl="")) %>% 
  mutate(Trait = "Angle")

# DW
slp_I_DW = model_I_DW %>% 
  emtrends(~ Genotype*HighTemperature*Drought,
           var = "Time", by = "Genotype") %>% 
  emmeans:::cld.emmGrid(by = "Genotype",
                        Letters = letters, sort = FALSE) %>% 
  as_tibble %>% 
  rename(slope = Time.trend, group = .group) %>% 
  mutate(Treatment = tnamesI[paste0(HighTemperature, Drought)],
         group = str_replace_all(string=group, pattern=" ", repl="")) %>% 
  mutate(Trait = "DW")

# LN
slp_I_LN = model_I_LN %>% 
  emtrends(~ Genotype*HighTemperature*Drought,
           var = "Time", by = "Genotype") %>% 
  emmeans:::cld.emmGrid(by = "Genotype",
                        Letters = letters, sort = FALSE) %>% 
  as_tibble %>% 
  rename(slope = Time.trend, group = .group) %>% 
  mutate(Treatment = tnamesI[paste0(HighTemperature, Drought)],
         group = str_replace_all(string=group, pattern=" ", repl="")) %>% 
  mutate(Trait = "LN")

# PR
slp_I_PR = model_I_PR %>% 
  emtrends(~ Genotype*HighTemperature*Drought,
           var = "Time", by = "Genotype") %>% 
  emmeans:::cld.emmGrid(by = "Genotype",
                        Letters = letters, sort = FALSE) %>% 
  as_tibble %>% 
  rename(slope = Time.trend, group = .group) %>% 
  mutate(Treatment = tnamesI[paste0(HighTemperature, Drought)],
         group = str_replace_all(string=group, pattern=" ", repl="")) %>% 
  mutate(Trait = "PR")

# RL
slp_I_RL = model_I_RL %>% 
  emtrends(~ Genotype*HighTemperature*Drought,
           var = "Time", by = "Genotype") %>% 
  emmeans:::cld.emmGrid(by = "Genotype",
                        Letters = letters, sort = FALSE) %>% 
  as_tibble %>% 
  rename(slope = Time.trend, group = .group) %>% 
  mutate(Treatment = tnamesI[paste0(HighTemperature, Drought)],
         group = str_replace_all(string=group, pattern=" ", repl="")) %>% 
  mutate(Trait = "RL")

# RWC
slp_I_RWC = model_I_RWC %>% 
  emtrends(~ Genotype*HighTemperature*Drought,
           var = "Time", by = "Genotype") %>% 
  emmeans:::cld.emmGrid(by = "Genotype",
                        Letters = letters, sort = FALSE) %>% 
  as_tibble %>% 
  rename(slope = Time.trend, group = .group) %>% 
  mutate(Treatment = tnamesI[paste0(HighTemperature, Drought)],
         group = str_replace_all(string=group, pattern=" ", repl="")) %>% 
  mutate(Trait = "RWC")

# BS
slp_I_BS = model_I_BS %>% 
  emtrends(~ Genotype*HighTemperature*Drought,
           var = "Time", by = "Genotype") %>% 
  emmeans:::cld.emmGrid(by = "Genotype",
                        Letters = letters, sort = FALSE) %>% 
  as_tibble %>% 
  rename(slope = Time.trend, group = .group) %>% 
  mutate(Treatment = tnamesI[paste0(HighTemperature, Drought)],
         group = str_replace_all(string=group, pattern=" ", repl="")) %>% 
  mutate(Trait = "BS")

# SLA
slp_I_SLA = model_I_SLA %>% 
  emtrends(~ Genotype*HighTemperature*Drought,
           var = "Time", by = "Genotype") %>% 
  emmeans:::cld.emmGrid(by = "Genotype",
                        Letters = letters, sort = FALSE) %>% 
  as_tibble %>% 
  rename(slope = Time.trend, group = .group) %>% 
  mutate(Treatment = tnamesI[paste0(HighTemperature, Drought)],
         group = str_replace_all(string=group, pattern=" ", repl="")) %>% 
  mutate(Trait = "SLA")


slopes_I = rbind(slp_I_Angle, slp_I_DW, slp_I_LN, slp_I_LS, slp_I_PR,
                 slp_I_RL, slp_I_RWC, slp_I_BS, slp_I_SLA) %>% 
  mutate(ymax = slope + SE, ymin = slope - SE)

slopes_I = slopes_I %>% 
  group_by(Trait) %>% 
  mutate(ylabel = max(ymax) + (max(ymax) - min(ymin))*0.1)

vlines = slopes_I %>%
  group_by(Trait) %>%
  summarise(xintercept = c(01.5,2.5,3.5))

rm(slp_I_LS, slp_I_Angle, slp_I_DW, slp_I_LN, slp_I_PR,
   slp_I_RL, slp_I_RWC, slp_I_BS, slp_I_SLA)

pretty_labels = c(
  Angle = "Leaf~angle",
  DW = "Rosette~dry~weight",
  HL = "Hypocotyl~length",
  LN = "Early~rosette~leaf~number",
  LS = "Leaf~size",
  PR = "Petiole~ratio",
  RL = "Root~length",
  RWC = "Relative~water~content",
  BS = "Blade~shape",
  SLA = "Specific~leaf~area")

slopes_I = slopes_I %>% 
  mutate(TraitPlot = pretty_labels[Trait])


(ggplot(slopes_I, aes(x = Genotype, y = slope, 
                     colour = Treatment, shape = Treatment)) +
  geom_point(position = position_dodge(width = 1)) +
  geom_errorbar(aes(ymax = ymax, ymin = ymin),
                position = position_dodge(width = 1)) +
  geom_text(aes(y = ylabel, label = group),
            position = position_dodge(width = 1), show.legend = FALSE) +
  geom_vline(data = vlines, aes(xintercept = xintercept), col = BLACK) +
  facet_wrap(~TraitPlot, scales = "free", ncol = 2, labeller = label_parsed) +
  scale_colour_manual(labels = labelsI, values = colorsI)  +
  scale_shape_manual(labels = labelsI, values = shapesI)  +
  theme_bw() +
  theme(panel.grid = element_blank(),
        legend.position = c(0.8, 0.01),
        legend.justification = c(1, 0),
        axis.title = element_blank())) %>% 
  ggsave("Results/Figure_3.png", plot = ., dpi = 300, 
         width = 14.5, height = 20, units = "cm",scale = 1.2)

# Figure 4 ----------------------------------------------------------------

# LS
slp_IIa_LS = model_IIa_LS %>% 
  emtrends(~ Genotype*Submergence,
           var = "Time", by = "Genotype") %>% 
  emmeans:::cld.emmGrid(by = "Genotype",
                        Letters = letters, sort = FALSE) %>% 
  as_tibble %>% 
  rename(slope = Time.trend, group = .group) %>% 
  mutate(Treatment = ifelse(Submergence == 1, "S", "C"),
         group = str_replace_all(string=group, pattern=" ", repl="")) %>% 
  mutate(Trait = "LS")

# Angle
slp_IIa_Angle = model_IIa_Angle %>% 
  emtrends(~ Genotype*Submergence,
           var = "Time", by = "Genotype") %>% 
  emmeans:::cld.emmGrid(by = "Genotype",
                        Letters = letters, sort = FALSE) %>% 
  as_tibble %>% 
  rename(slope = Time.trend, group = .group) %>% 
  mutate(Treatment = ifelse(Submergence == 1, "S", "C"),
         group = str_replace_all(string=group, pattern=" ", repl="")) %>% 
  mutate(Trait = "Angle")

# DW
slp_IIa_DW = model_IIa_DW %>% 
  emtrends(~ Genotype*Submergence,
           var = "Time", by = "Genotype") %>% 
  emmeans:::cld.emmGrid(by = "Genotype",
                        Letters = letters, sort = FALSE) %>% 
  as_tibble %>% 
  rename(slope = Time.trend, group = .group) %>% 
  mutate(Treatment = ifelse(Submergence == 1, "S", "C"),
         group = str_replace_all(string=group, pattern=" ", repl="")) %>% 
  mutate(Trait = "DW")

# LN
slp_IIa_LN = model_IIa_LN %>% 
  emtrends(~ Genotype*Submergence,
           var = "Time", by = "Genotype") %>% 
  emmeans:::cld.emmGrid(by = "Genotype",
                        Letters = letters, sort = FALSE) %>% 
  as_tibble %>% 
  rename(slope = Time.trend, group = .group) %>% 
  mutate(Treatment = ifelse(Submergence == 1, "S", "C"),
         group = str_replace_all(string=group, pattern=" ", repl="")) %>% 
  mutate(Trait = "LN")

# PR
slp_IIa_PR = model_IIa_PR %>% 
  emtrends(~ Genotype*Submergence,
           var = "Time", by = "Genotype") %>% 
  emmeans:::cld.emmGrid(by = "Genotype",
                        Letters = letters, sort = FALSE) %>% 
  as_tibble %>% 
  rename(slope = Time.trend, group = .group) %>% 
  mutate(Treatment = ifelse(Submergence == 1, "S", "C"),
         group = str_replace_all(string=group, pattern=" ", repl="")) %>% 
  mutate(Trait = "PR")

# RL
slp_IIa_RL = model_IIa_RL %>% 
  emtrends(~ Genotype*Submergence,
           var = "Time", by = "Genotype") %>% 
  emmeans:::cld.emmGrid(by = "Genotype",
                        Letters = letters, sort = FALSE) %>% 
  as_tibble %>% 
  rename(slope = Time.trend, group = .group) %>% 
  mutate(Treatment = ifelse(Submergence == 1, "S", "C"),
         group = str_replace_all(string=group, pattern=" ", repl="")) %>% 
  mutate(Trait = "RL")

# RWC
slp_IIa_RWC = model_IIa_RWC %>% 
  emtrends(~ Genotype*Submergence,
           var = "Time", by = "Genotype") %>% 
  emmeans:::cld.emmGrid(by = "Genotype",
                        Letters = letters, sort = FALSE) %>% 
  as_tibble %>% 
  rename(slope = Time.trend, group = .group) %>% 
  mutate(Treatment = ifelse(Submergence == 1, "S", "C"),
         group = str_replace_all(string=group, pattern=" ", repl="")) %>% 
  mutate(Trait = "RWC")

# BS
slp_IIa_BS = model_IIa_BS %>% 
  emtrends(~ Genotype*Submergence,
           var = "Time", by = "Genotype") %>% 
  emmeans:::cld.emmGrid(by = "Genotype",
                        Letters = letters, sort = FALSE) %>% 
  as_tibble %>% 
  rename(slope = Time.trend, group = .group) %>% 
  mutate(Treatment = ifelse(Submergence == 1, "S", "C"),
         group = str_replace_all(string=group, pattern=" ", repl="")) %>% 
  mutate(Trait = "BS")

# SLA
slp_IIa_SLA = model_IIa_SLA %>% 
  emtrends(~ Genotype*Submergence,
           var = "Time", by = "Genotype") %>% 
  emmeans:::cld.emmGrid(by = "Genotype",
                        Letters = letters, sort = FALSE) %>% 
  as_tibble %>% 
  rename(slope = Time.trend, group = .group) %>% 
  mutate(Treatment = ifelse(Submergence == 1, "S", "C"),
         group = str_replace_all(string=group, pattern=" ", repl="")) %>% 
  mutate(Trait = "SLA")


slopes_IIa = rbind(slp_IIa_Angle, slp_IIa_DW, slp_IIa_LN, slp_IIa_LS, slp_IIa_PR,
                 slp_IIa_RL, slp_IIa_RWC, slp_IIa_BS, slp_IIa_SLA) %>% 
  mutate(ymax = slope + SE, ymin = slope - SE)

slopes_IIa = slopes_IIa %>% 
  group_by(Trait) %>% 
  mutate(ylabel = max(ymax) + (max(ymax) - min(ymin))*0.1)

vlines = slopes_IIa %>%
  group_by(Trait) %>%
  summarise(xintercept = c(01.5,2.5,3.5))

rm(slp_IIa_LS, slp_IIa_Angle, slp_IIa_DW, slp_IIa_LN, slp_IIa_PR,
   slp_IIa_RL, slp_IIa_RWC, slp_IIa_BS, slp_IIa_SLA)

pretty_labels = c(
  Angle = "Leaf~angle",
  DW = "Rosette~dry~weight",
  HL = "Hypocotyl~length",
  LN = "Early~rosette~leaf~number",
  LS = "Leaf~size",
  PR = "Petiole~ratio",
  RL = "Root~length",
  RWC = "Relative~water~content",
  BS = "Blade~shape",
  SLA = "Specific~leaf~area")

slopes_IIa = slopes_IIa %>% 
  mutate(TraitPlot = pretty_labels[Trait])


(ggplot(slopes_IIa, aes(x = Genotype, y = slope, 
                      colour = Treatment, shape = Treatment)) +
    geom_point(position = position_dodge(width = 1)) +
    geom_errorbar(aes(ymax = ymax, ymin = ymin),
                  position = position_dodge(width = 1)) +
    geom_text(aes(y = ylabel, label = group),
              position = position_dodge(width = 1), show.legend = FALSE) +
    geom_vline(data = vlines, aes(xintercept = xintercept), col = BLACK) +
    facet_wrap(~TraitPlot, scales = "free", ncol = 3, labeller = label_parsed) +
    scale_colour_manual(labels = labelsIIa, values = colorsIIa)  +
    scale_shape_manual(labels = labelsIIa, values = shapesIIa)  +  
    theme_bw() +
    theme(panel.grid = element_blank(),
          legend.position = "bottom",
          axis.title = element_blank())) %>% 
  ggsave("Results/Figure_4.png", plot = ., dpi = 300, 
         width = 16, height = 16, units = "cm", scale = 1.2)

# Figure 5 ----------------------------------------------------------------

# LS
slp_IIb_LS = model_IIb_LS %>% 
  emtrends(~ Genotype*Submergence*Drought,
           var = "Time", by = "Genotype") %>% 
  emmeans:::cld.emmGrid(by = "Genotype",
                        Letters = letters, sort = FALSE) %>% 
  as_tibble %>% 
  rename(slope = Time.trend, group = .group) %>% 
  mutate(Treatment = tnamesII[paste0(Submergence, Drought)],
         group = str_replace_all(string=group, pattern=" ", repl="")) %>% 
  mutate(Trait = "LS")

# Angle
slp_IIb_Angle = model_IIb_Angle %>% 
  emtrends(~ Genotype*Submergence*Drought,
           var = "Time", by = "Genotype") %>% 
  emmeans:::cld.emmGrid(by = "Genotype",
                        Letters = letters, sort = FALSE) %>% 
  as_tibble %>% 
  rename(slope = Time.trend, group = .group) %>% 
  mutate(Treatment = tnamesII[paste0(Submergence, Drought)],
         group = str_replace_all(string=group, pattern=" ", repl="")) %>% 
  mutate(Trait = "Angle")

# DW
slp_IIb_DW = model_IIb_DW %>% 
  emtrends(~ Genotype*Submergence*Drought,
           var = "Time", by = "Genotype") %>% 
  emmeans:::cld.emmGrid(by = "Genotype",
                        Letters = letters, sort = FALSE) %>% 
  as_tibble %>% 
  rename(slope = Time.trend, group = .group) %>% 
  mutate(Treatment = tnamesII[paste0(Submergence, Drought)],
         group = str_replace_all(string=group, pattern=" ", repl="")) %>% 
  mutate(Trait = "DW")

# LN
slp_IIb_LN = model_IIb_LN %>% 
  emtrends(~ Genotype*Submergence*Drought,
           var = "Time", by = "Genotype") %>% 
  emmeans:::cld.emmGrid(by = "Genotype",
                        Letters = letters, sort = FALSE) %>% 
  as_tibble %>% 
  rename(slope = Time.trend, group = .group) %>% 
  mutate(Treatment = tnamesII[paste0(Submergence, Drought)],
         group = str_replace_all(string=group, pattern=" ", repl="")) %>% 
  mutate(Trait = "LN")

# PR
slp_IIb_PR = model_IIb_PR %>% 
  emtrends(~ Genotype*Submergence*Drought,
           var = "Time", by = "Genotype") %>% 
  emmeans:::cld.emmGrid(by = "Genotype",
                        Letters = letters, sort = FALSE) %>% 
  as_tibble %>% 
  rename(slope = Time.trend, group = .group) %>% 
  mutate(Treatment = tnamesII[paste0(Submergence, Drought)],
         group = str_replace_all(string=group, pattern=" ", repl="")) %>% 
  mutate(Trait = "PR")

# RL
slp_IIb_RL = model_IIb_RL %>% 
  emtrends(~ Genotype*Submergence*Drought,
           var = "Time", by = "Genotype") %>% 
  emmeans:::cld.emmGrid(by = "Genotype",
                        Letters = letters, sort = FALSE) %>% 
  as_tibble %>% 
  rename(slope = Time.trend, group = .group) %>% 
  mutate(Treatment = tnamesII[paste0(Submergence, Drought)],
         group = str_replace_all(string=group, pattern=" ", repl="")) %>% 
  mutate(Trait = "RL")

# RWC
slp_IIb_RWC = model_IIb_RWC %>% 
  emtrends(~ Genotype*Submergence*Drought,
           var = "Time", by = "Genotype") %>% 
  emmeans:::cld.emmGrid(by = "Genotype",
                        Letters = letters, sort = FALSE) %>% 
  as_tibble %>% 
  rename(slope = Time.trend, group = .group) %>% 
  mutate(Treatment = tnamesII[paste0(Submergence, Drought)],
         group = str_replace_all(string=group, pattern=" ", repl="")) %>% 
  mutate(Trait = "RWC")

# BS
slp_IIb_BS = model_IIb_BS %>% 
  emtrends(~ Genotype*Submergence*Drought,
           var = "Time", by = "Genotype") %>% 
  emmeans:::cld.emmGrid(by = "Genotype",
                        Letters = letters, sort = FALSE) %>% 
  as_tibble %>% 
  rename(slope = Time.trend, group = .group) %>% 
  mutate(Treatment = tnamesII[paste0(Submergence, Drought)],
         group = str_replace_all(string=group, pattern=" ", repl="")) %>% 
  mutate(Trait = "BS")

# SLA
slp_IIb_SLA = model_IIb_SLA %>% 
  emtrends(~ Genotype*Submergence*Drought,
           var = "Time", by = "Genotype") %>% 
  emmeans:::cld.emmGrid(by = "Genotype",
                        Letters = letters, sort = FALSE) %>% 
  as_tibble %>% 
  rename(slope = Time.trend, group = .group) %>% 
  mutate(Treatment = tnamesII[paste0(Submergence, Drought)],
         group = str_replace_all(string=group, pattern=" ", repl="")) %>% 
  mutate(Trait = "SLA")


slopes_IIb = rbind(slp_IIb_Angle, slp_IIb_DW, slp_IIb_LN, slp_IIb_LS, slp_IIb_PR,
                   slp_IIb_RL, slp_IIb_RWC, slp_IIb_BS, slp_IIb_SLA) %>% 
  mutate(ymax = slope + SE, ymin = slope - SE)

slopes_IIb = slopes_IIb %>% 
  group_by(Trait) %>% 
  mutate(ylabel = max(ymax) + (max(ymax) - min(ymin))*0.1)

vlines = slopes_IIb %>%
  group_by(Trait) %>%
  summarise(xintercept = c(01.5,2.5,3.5))

rm(slp_IIb_LS, slp_IIb_Angle, slp_IIb_DW, slp_IIb_LN, slp_IIb_PR,
   slp_IIb_RL, slp_IIb_RWC, slp_IIb_BS, slp_IIb_SLA)

pretty_labels = c(
  Angle = "Leaf~angle",
  DW = "Rosette~dry~weight",
  HL = "Hypocotyl~length",
  LN = "Early~rosette~leaf~number",
  LS = "Leaf~size",
  PR = "Petiole~ratio",
  RL = "Root~length",
  RWC = "Relative~water~content",
  BS = "Blade~shape",
  SLA = "Specific~leaf~area")

slopes_IIb = slopes_IIb %>% 
  mutate(TraitPlot = pretty_labels[Trait])


(ggplot(slopes_IIb, aes(x = Genotype, y = slope, 
                        colour = Treatment, shape = Treatment)) +
    geom_point(position = position_dodge(width = 1)) +
    geom_errorbar(aes(ymax = ymax, ymin = ymin),
                  position = position_dodge(width = 1)) +
    geom_text(aes(y = ylabel, label = group),
              position = position_dodge(width = 1), show.legend = FALSE) +
    geom_vline(data = vlines, aes(xintercept = xintercept), col = BLACK) +
    facet_wrap(~TraitPlot, scales = "free", ncol = 2, labeller = label_parsed) +
    scale_colour_manual(labels = labelsIIb, values = colorsIIb)  +
    scale_shape_manual(labels = labelsIIb, values = shapesIIb)  +  
    theme_bw() +
    theme(panel.grid = element_blank(),
          legend.position = c(0.8, 0.01),
          legend.justification = c(1, 0),
          axis.title = element_blank())) %>% 
  ggsave("Results/Figure_5.png", plot = ., dpi = 300, 
         width = 14.5, height = 20, units = "cm", scale = 1.2)


# Figure 6 ----------------------------------------------------------------

# Yield
sigma = sqrt(sum(as.numeric(VarCorr(model_III_Y)[,1])))

Yield_grid = ref_grid(model_III_Y) 

Yield_means = Yield_grid %>% 
  summary(infer = c(F, F),  type = "response", bias.adjust = TRUE, sigma = sigma) %>% 
  mutate(Treatment = tnames[paste0(HighTemperature, Submergence, Drought)]) %>% 
  filter(!is.na(Treatment)) %>% 
  mutate(Treatment = factor(Treatment, levels = c("C", "D", "HT", "HTD", "S", "PSD")))

Yield_letters = Yield_grid %>% 
  emmeans:::cld.emmGrid(by = "Genotype", Letters = letters, sort = FALSE) %>% 
  as_tibble %>% 
  mutate(Treatment = tnames[paste0(HighTemperature, Submergence, Drought)],
         .group = str_replace_all(string=.group, pattern=" ", repl="")) %>% 
  filter(!is.na(Treatment)) %>% 
  mutate(Treatment = factor(Treatment, levels = c("C", "D", "HT", "HTD", "S", "PSD")))

data_III_Y = data_III_Y %>% 
  mutate(Treatment = factor(Treatment, levels = c("C", "D", "HT", "HTD", "S", "PSD")))

Yield_plot = ggplot(data = Yield_means, aes(x = Genotype, y = response,
                                            colour = Treatment, shape = Treatment)) +
  geom_point(position = position_dodge(width = 1), show.legend = F) + 
  geom_point(data = data_III_Y, mapping = aes(y = Yield), size = 0.75,
             position = position_jitterdodge(dodge.width = 1, jitter.width = 0.3)) +
  geom_errorbar(aes(ymax = response + SE, ymin = response - SE), 
                position = position_dodge(width = 1)) +
  geom_vline(data = data.frame(), aes(xintercept = c(01.5,2.5,3.5))) +
  geom_text(data = Yield_letters, aes(y = 1100, label = .group),
            position = position_dodge(width = 1), show.legend = FALSE) +
  scale_colour_manual(values = c(C = BLUISH_GREEN, S = BLACK, D = YELLOW,
                                 PSD = SKY_BLUE,  HT = ORANGE, HTD = REDDISH_PURPLE)) +
  scale_shape_manual(values = c(C = 1, S = 2, D = 15,PSD = 16,  HT = 17, HTD = 6)) +
  scale_y_continuous(trans = "log", breaks = c(2.5,50,1000)) + 
  labs(y = quote(Yield~per~plant~(mg))) +
  guides(colour = "none", shape = "none") +
  theme_bw() +
  theme(panel.grid = element_blank())


# FT
sigma = sqrt(sum(as.numeric(VarCorr(model_III_FT)[,1])))

FT_grid = ref_grid(model_III_FT) 

FT_means = FT_grid %>% 
  summary(infer = c(F, F),  type = "response", bias.adjust = TRUE, sigma = sigma) %>% 
  mutate(Treatment = tnames[paste0(HighTemperature, Submergence, Drought)]) %>% 
  filter(!is.na(Treatment)) %>% 
  mutate(Treatment = factor(Treatment, levels = c("C", "D", "HT", "HTD", "S", "PSD")))

FT_letters = FT_grid %>% 
  emmeans:::cld.emmGrid(by = "Genotype", Letters = letters, sort = FALSE) %>% 
  as_tibble %>% 
  mutate(Treatment = tnames[paste0(HighTemperature, Submergence, Drought)],
         .group = str_replace_all(string=.group, pattern=" ", repl="")) %>% 
  filter(!is.na(Treatment)) %>% 
  mutate(Treatment = factor(Treatment, levels = c("C", "D", "HT", "HTD", "S", "PSD")))

data_III_FT = data_III_FT %>% 
  mutate(Treatment = factor(Treatment, levels = c("C", "D", "HT", "HTD", "S", "PSD")))

FT_plot = ggplot(data = FT_means, aes(x = Genotype, y = response,
                                            colour = Treatment, shape = Treatment)) +
  geom_point(position = position_dodge(width = 1), show.legend = F) + 
  geom_point(data = data_III_FT, mapping = aes(y = FT), size = 0.75,
             position = position_jitterdodge(dodge.width = 1, jitter.width = 0.3)) +
  geom_errorbar(aes(ymax = response + SE, ymin = response - SE), 
                position = position_dodge(width = 1)) +
  geom_vline(data = data.frame(), aes(xintercept = c(01.5,2.5,3.5))) +
  geom_text(data = FT_letters, aes(y = 180, label = .group),
            position = position_dodge(width = 1), show.legend = FALSE) +
  scale_colour_manual(values = c(C = BLUISH_GREEN, S = BLACK, D = YELLOW,
                                 PSD = SKY_BLUE,  HT = ORANGE, HTD = REDDISH_PURPLE)) +
  scale_shape_manual(values = c(C = 1, S = 2, D = 15,PSD = 16,  HT = 17, HTD = 6)) +
  scale_y_continuous(trans = "log", breaks = c(40,60,100,160)) + 
  labs(y = quote(Flowering~time~(days))) +
  guides(colour = "none", shape = "none") +
  theme_bw() +
  theme(panel.grid = element_blank())


# TLN
sigma = sqrt(sum(as.numeric(VarCorr(model_III_TLN)[,1])))

TLN_grid = ref_grid(model_III_TLN) 

TLN_means = TLN_grid %>% 
  summary(infer = c(F, F),  type = "response", bias.adjust = TRUE, sigma = sigma) %>% 
  mutate(Treatment = tnames[paste0(HighTemperature, Submergence, Drought)]) %>% 
  filter(!is.na(Treatment)) %>% 
  mutate(Treatment = factor(Treatment, levels = c("C", "D", "HT", "HTD", "S", "PSD")))

TLN_letters = TLN_grid %>% 
  emmeans:::cld.emmGrid(by = "Genotype", Letters = letters, sort = FALSE) %>% 
  as_tibble %>% 
  mutate(Treatment = tnames[paste0(HighTemperature, Submergence, Drought)],
         .group = str_replace_all(string=.group, pattern=" ", repl="")) %>% 
  filter(!is.na(Treatment)) %>% 
  mutate(Treatment = factor(Treatment, levels = c("C", "D", "HT", "HTD", "S", "PSD")))

data_III_TLN = data_III_TLN %>% 
  mutate(Treatment = factor(Treatment, levels = c("C", "D", "HT", "HTD", "S", "PSD")))

TLN_plot = ggplot(data = TLN_means, aes(x = Genotype, y = response,
                                      colour = Treatment, shape = Treatment)) +
  geom_point(position = position_dodge(width = 1), show.legend = F) + 
  geom_point(data = data_III_TLN, mapping = aes(y = LN), size = 0.75,
             position = position_jitterdodge(dodge.width = 1, jitter.width = 0.3)) +
  geom_errorbar(aes(ymax = response + SE, ymin = response - SE), 
                position = position_dodge(width = 1)) +
  geom_vline(data = data.frame(), aes(xintercept = c(01.5,2.5,3.5))) +
  geom_text(data = TLN_letters, aes(y = 95, label = .group),
            position = position_dodge(width = 1), show.legend = F) +
  scale_colour_manual(values = c(C = BLUISH_GREEN, S = BLACK, D = YELLOW,
                                 PSD = SKY_BLUE,  HT = ORANGE, HTD = REDDISH_PURPLE),
                      labels = c(C = "Control", S = "Post-submergence",
                                 D = "Drought", PSD = "Post-submergence drought",
                                 HT = "High temperature", HTD = "High temperature & drought")) +
  scale_shape_manual(values = c(C = 1, S = 2, D = 15,PSD = 16,  HT = 17, HTD = 6),
                     labels = c(C = "Control", S = "Post-submergence",
                                D = "Drought", PSD = "Post-submergence drought",
                                HT = "High temperature", HTD = "High temperature & drought")) +
  scale_y_continuous(trans = "log", breaks = c(7.5,20,55)) + 
  labs(y = quote(Total~rosette~leaf~number), colour = "", shape = "") +
  theme_bw() +
  theme(panel.grid = element_blank())


(Yield_plot + FT_plot + TLN_plot +
  plot_annotation(tag_levels = "A") + 
  plot_layout(ncol = 1, guides = "collect") &
  theme(axis.title = element_text(size = 10),
        legend.position = "bottom", 
        legend.direction = "horizontal")) %>% 
  ggsave("Results/Figure_6.png", plot = ., dpi = 300, 
         width = 16, height = 16, units = "cm",scale = 1.3)


# Table S1 ----------------------------------------------------------------

# Construct the basic table
traits_I_II = c("Blade shape", "Early rosette leaf number", "Leaf angle",
               "Leaf average area", "Petiole ratio", "Relative water content",
               "Root length", "Rosette dry weight", "Specific leaf area")

transforms_I_II = c("log", "log", "logit", "log", "logit", "logit",
                    "log", "log", "log")

traits_III = c("Flowering time", "Total rosette leaf number", "Yield")
transforms_III = c("log", "log", "log")

TableS1 = tibble(Trait = c(rep(traits_I_II, 3), traits_III),
                 XP    = rep(paste0("Experiment_",c("I","IIa","IIb","III")), 
                             times = c(9,9,9,3)),
                 Transform = c(rep(transforms_I_II, 3), transforms_III))

# Add Ntotal and Nbatch
TableS1 = mutate(TableS1,
         Ntotal = c(nrow(data_I_BS), nrow(data_I_LN), nrow(data_I_Angle),
                    nrow(data_I_ALA), nrow(data_I_PR), nrow(data_I_RWC),
                    nrow(data_I_RL), nrow(data_I_DW), nrow(data_I_SLA),
                    nrow(data_IIa_BS), nrow(data_IIa_LN), nrow(data_IIa_Angle),
                    nrow(data_IIa_ALA), nrow(data_IIa_PR), nrow(data_IIa_RWC),
                    nrow(data_IIa_RL), nrow(data_IIa_DW), nrow(data_IIa_SLA),
                    nrow(data_IIb_BS), nrow(data_IIb_LN), nrow(data_IIb_Angle),
                    nrow(data_IIb_ALA), nrow(data_IIb_PR), nrow(data_IIb_RWC),
                    nrow(data_IIb_RL), nrow(data_IIb_DW), nrow(data_IIb_SLA),
                    nrow(data_III_FT), nrow(data_III_TLN), nrow(data_III_Y)),
         Nbatch = c(rep(8,27), rep(3,3)))

# Summary of datasets
summarise_dataset = function(data, XP3 = FALSE) {
  if(XP3) {
    datag = data %>% group_by(Repetition, Genotype, Treatment)  
  } else {
    datag = data %>% group_by(Repetition, Genotype, Treatment, Timepoint)  
  }
  datag %>% 
    summarise(n = n()) %>% 
    ungroup() %>% 
    summarise(mu = format(mean(n), digits = 1, nsmall= 1),
              sd = format(sd(n), digits = 1, nsmall= 1),
              mx = max(n),
              mn = min(n))
}
summaries = map2_dfr(list(data_I_BS, data_I_LN, data_I_Angle, data_I_ALA,
                      data_I_PR, data_I_RWC, data_I_RL, data_I_DW, data_I_SLA,
                      data_IIa_BS, data_IIa_LN, data_IIa_Angle, data_IIa_ALA,
                      data_IIa_PR, data_IIa_RWC, data_IIa_RL, data_IIa_DW, 
                      data_IIa_SLA, data_IIb_BS, data_IIb_LN, data_IIb_Angle, 
                      data_IIb_ALA, data_IIb_PR, data_IIb_RWC, data_IIb_RL, 
                      data_IIb_DW, data_IIb_SLA, data_III_FT, data_III_TLN,
                      data_III_Y), rep(c(F,T), c(27,3)),
                     summarise_dataset)

TableS1 = bind_cols(TableS1, summaries)

write_csv(TableS1, file = "Results/TableS1.csv")

# Table S2 ----------------------------------------------------------------

TableS2 = mutate(anova_I_BS, Trait = "Blade shape") %>% 
       bind_rows(mutate(anova_I_LN, Trait = "Early rosette leaf number")) %>% 
       bind_rows(mutate(anova_I_Angle, Trait = "Leaf angle")) %>% 
       bind_rows(mutate(anova_I_ALA, Trait = "Leaf average area")) %>% 
       bind_rows(mutate(anova_I_PR, Trait = "Petiole ratio")) %>% 
       bind_rows(mutate(anova_I_RWC, Trait = "Relative water content")) %>% 
       bind_rows(mutate(anova_I_RL, Trait = "Root length")) %>% 
       bind_rows(mutate(anova_I_DW, Trait = "Rosette dry weight")) %>% 
       bind_rows(mutate(anova_I_SLA, Trait = "Specific leaf area"))

# Shorter treatment names
shorterm = c(Genotype = "i_G",
             HighTemperature = "i_HT",
             `Genotype:HighTemperature` = "i_HT:G",
             `Genotype:Time` = "d_G",
             `HighTemperature:Time` = "d_HT",
             `Time:Drought` = "d_D",
             `Genotype:HighTemperature:Time` = "d_HT:G",
             `HighTemperature:Time:Drought` = "d_HT:D",
             `Genotype:Time:Drought` = "d_D:G",
             `Genotype:HighTemperature:Time:Drought` = "d_HT:D:G",
             `Time` = "d")

# Construct p-value table
TableS2 = TableS2 %>% 
  mutate(Term = ifelse(Term %in% names(shorterm), shorterm[Term],NA)) %>% 
  filter(!is.na(Term)) %>% 
  dplyr::select(Term, `p-value`, Trait) %>% 
  rename(pval = "p-value") %>% 
  mutate(pval = log10(pval),
         pval = ifelse(is.finite(pval), pval, -16),
         pval = sprintf(pval, fmt = '%#.1f')) %>% 
  pivot_wider(values_from = pval, names_from = Term)


write_csv(TableS2, file = "Results/TableS2.csv")

# Table S3 ----------------------------------------------------------------

TableS3 = mutate(anova_IIa_BS, Trait = "Blade shape") %>% 
  bind_rows(mutate(anova_IIa_LN, Trait = "Early rosette leaf number")) %>% 
  bind_rows(mutate(anova_IIa_Angle, Trait = "Leaf angle")) %>% 
  bind_rows(mutate(anova_IIa_ALA, Trait = "Leaf average area")) %>% 
  bind_rows(mutate(anova_IIa_PR, Trait = "Petiole ratio")) %>% 
  bind_rows(mutate(anova_IIa_RWC, Trait = "Relative water content")) %>% 
  bind_rows(mutate(anova_IIa_RL, Trait = "Root length")) %>% 
  bind_rows(mutate(anova_IIa_DW, Trait = "Rosette dry weight")) %>% 
  bind_rows(mutate(anova_IIa_SLA, Trait = "Specific leaf area"))

# Shorter treatment names
shorterm = c(Genotype = "i_G",
             Time = "d",
             `Genotype:Time` = "d_G",
             `Time:Submergence` = "d_S",
             `Genotype:Time:Submergence` = "d_S:G")

# Construct p-value table
TableS3 = TableS3 %>% 
  mutate(Term = ifelse(Term %in% names(shorterm), shorterm[Term],NA)) %>% 
  filter(!is.na(Term)) %>% 
  dplyr::select(Term, `p-value`, Trait) %>% 
  rename(pval = "p-value") %>% 
  mutate(pval = log10(pval),
         pval = ifelse(is.finite(pval), pval, -16),
         pval = sprintf(pval, fmt = '%#.1f')) %>% 
  pivot_wider(values_from = pval, names_from = Term)


write_csv(TableS3, file = "Results/TableS3.csv")


# Table S4 ----------------------------------------------------------------

TableS4 = mutate(anova_IIb_BS, Trait = "Blade shape") %>% 
  bind_rows(mutate(anova_IIb_LN, Trait = "Early rosette leaf number")) %>% 
  bind_rows(mutate(anova_IIb_Angle, Trait = "Leaf angle")) %>% 
  bind_rows(mutate(anova_IIb_ALA, Trait = "Leaf average area")) %>% 
  bind_rows(mutate(anova_IIb_PR, Trait = "Petiole ratio")) %>% 
  bind_rows(mutate(anova_IIb_RWC, Trait = "Relative water content")) %>% 
  bind_rows(mutate(anova_IIb_RL, Trait = "Root length")) %>% 
  bind_rows(mutate(anova_IIb_DW, Trait = "Rosette dry weight")) %>% 
  bind_rows(mutate(anova_IIb_SLA, Trait = "Specific leaf area"))

# Shorter treatment names
shorterm = c(Genotype = "i_G",
             Submergence = "i_S",
             `Genotype:Submergence` = "i_S:G",
             Time = "d",
             `Genotype:Time` = "d_G",
             `Submergence:Time` = "d_S",
             `Time:Drought` = "d_D",
             `Genotype:Submergence:Time` = "d_S:G",
             `Submergence:Time:Drought` = "d_S:D",
             `Genotype:Time:Drought` = "d_D:G",
             `Genotype:Submergence:Time:Drought` = "d_S:D:G")

# Construct p-value table
TableS4 = TableS4 %>% 
  mutate(Term = ifelse(Term %in% names(shorterm), shorterm[Term],NA)) %>% 
  filter(!is.na(Term)) %>% 
  dplyr::select(Term, `p-value`, Trait) %>% 
  rename(pval = "p-value") %>% 
  mutate(pval = log10(pval),
         pval = ifelse(is.finite(pval), pval, -16),
         pval = sprintf(pval, fmt = '%#.1f')) %>% 
  pivot_wider(values_from = pval, names_from = Term)


write_csv(TableS4, file = "Results/TableS4.csv")

# Table S5 ----------------------------------------------------------------


TableS5 = mutate(anova_III_FT, Trait = "Flowering time") %>% 
  bind_rows(mutate(anova_III_TLN, Trait = "Total rosette leaf number")) %>% 
  bind_rows(mutate(anova_III_Y, Trait = "Yield"))

# Shorter treatment names
shorterm = c(Drought = "D", 
             `Drought:Submergence` = "S:D",
             Genotype = "G",
             `Genotype:Drought` = "D:G",
             `Genotype:Drought:Submergence` = "S:D:G",
             `Genotype:HighTemperature` = "HT:G",
             `Genotype:HighTemperature:Drought` = "HT:D:G",
             `Genotype:Drought:HighTemperature` = "HT:D:G",
             `Genotype:Submergence` = "S:G",
             HighTemperature = "HT",
             `HighTemperature:Drought` = "HT:D",
             `Drought:HighTemperature` = "HT:D",
             Submergence = "S")

# Construct p-value table
TableS5 = TableS5 %>% 
  mutate(Term = ifelse(Term %in% names(shorterm), shorterm[Term],NA)) %>% 
  filter(!is.na(Term)) %>% 
  dplyr::select(Term, `p-value`, Trait) %>% 
  rename(pval = "p-value") %>% 
  mutate(pval = log10(pval),
         pval = ifelse(is.finite(pval), pval, -16),
         pval = sprintf(pval, fmt = '%#.1f')) %>% 
  pivot_wider(values_from = pval, names_from = Term)


write_csv(TableS5, file = "Results/TableS5.csv")


# Drying curve - Figure S1 ------------------------------------------------

# Identify the treatment from the final relative pellet weight
FinalRW = filter(DryingCurve, Time == 12)
Control = filter(FinalRW, RelativeWeight > 0.8)$ID
Drought = filter(FinalRW, RelativeWeight < 0.3)$ID

DryingCurve = mutate(DryingCurve, 
                     Treatment = case_when(
                       ID %in% Control ~ "Control",
                       ID %in% Drought ~ "Drought",
                       TRUE            ~ "Oher"
                     ))

# Calculate median of the individual traces
sum_DryingCurve = group_by(DryingCurve, Treatment, Time) %>%
  summarise(mu = median(RelativeWeight, na.rm = T),
            upr = quantile(RelativeWeight, probs = 0.90, na.rm = T),
            lwr = quantile(RelativeWeight, probs = 0.10, na.rm = T))

# Check the individual traces
pellet_weights = ggplot(data = filter(sum_DryingCurve, Treatment %in% c("Control", "Drought")), 
                        aes(x = Time, y = mu, col = Treatment, linetype = Treatment, shape = Treatment)) + 
  geom_line() + 
  geom_point() +
  geom_ribbon(aes(ymax = upr, ymin = lwr, fill = Treatment), col = NA, alpha = 0.2) +
  ylim(0,1.05) +
  labs(y = "Relative pellet weight", x = "Time (days)", fill = "", col = "", 
       shape = "", linetype = "") + 
  scale_color_manual(values = c(Control = "Blue", Drought = "Red")) + 
  scale_fill_manual(values = c(Control = "Blue", Drought = "Red")) + 
  scale_x_continuous(breaks = c(0,2,4,6,8,10,12), limits = c(0,12)) +
  theme_bw() +
  theme(panel.grid = element_blank(), 
        legend.position = c(0.2,0.2))

ggsave(filename = "Results/Figure_S1.png", plot = pellet_weights,
       width = 8, height = 8, units = "cm",device = "png", dpi = 300)

rm(DryingCurve, FinalRW, pellet_weights, sum_DryingCurve, Control, Drought)
```
